# *GEfetch2R*: fetching single-cell/bulk RNA-seq data from public repositories to R and benchmarking the subsequent format conversion tools

**DOI:** 10.1101/2023.11.18.567507

**Authors:** Yabing Song, Jianbin Wang, Jiaxin Gao

## Abstract

**Background:** Downloading and reanalyzing the existing single-cell RNA sequencing (scRNA-seq) data provides an efficient choice to gain clues and new insights. However, no tool can fetch the diverse scRNA-seq data types (raw data, count matrix, and processed object) distributed in various repositories, process and load the downloaded data to R, convert formats between scRNA-seq objects, and benchmark the format conversion tools.

**Findings:** Here, we present *GEfetch2R*, an R package with Docker image to (i) download diverse scRNA-seq data types, including raw data (SRA and ENA), count matrices (GEO, UCSC Cell Browser, and PanglaoDB), and processed objects (Zenodo, CELLxGENE, and HCA); (ii) process the downloaded data, load output/downloaded count matrices and annotations to R (*SeuratObject*/*DESeqDataSet*), filter the *SeuratObject* based on cell metadata and genes, and merge multiple *SeuratObjects* if applicable; (iii) convert formats between the widely used scRNA-seq objects, including *SeuratObject*, *AnnData*, *SingleCellExperiment*, *CellDataSet*/*cell_data_set*, and *loom*, and benchmark format conversion tools in terms of information kept, usability, running time, and scalability to guide the tool selection. Furthermore, *GEfetch2R* can also download, process, and load bulk RNA-seq raw data (SRA and ENA) and count matrices (GEO) to R (*DESeqDataSet*).

**Conclusions:** *GEfetch2R* is an R package dedicated to facilitating researchers to access and explore the existing gene expression data from various public repositories. It can function as a data downloader (supports all three scRNA-seq and two bulk RNA-seq data types), a data processor (processes and loads the output/downloaded count matrices and annotations to R), and an object format converter (between the widely used scRNA-seq objects).

## Introduction

In recent years, single-cell RNA sequencing (scRNA-seq) has emerged as a powerful tool to reveal cellular heterogeneity and fundamental characteristics of gene expression [1]. As a result, thousands of scRNA-seq datasets have been generated and uploaded to public repositories. Downloading and reanalyzing the existing scRNA-seq data provides an efficient choice to gain clues and new insights. For example, rare cell type identification [2] and pan-cancer scRNA-seq analysis [3] usually require a huge number of samples and cells, which is extremely expensive and time-consuming if not integrating the available scRNA-seq datasets. As well, exploring existing relevant scRNA-seq data can accelerate the progress of one’s own research.

The existing scRNA-seq data are in three types: raw data (*sra*/*fastq*/*bam*), count matrix, and processed object (e.g. *rds* and *h5ad*), which are stored in different repositories. The raw data are mainly stored in Sequence Read Archive (SRA) [4] and European Nucleotide Archive (ENA) [5], the count matrices can be found in Gene Expression Omnibus (GEO) [6] and some scRNA-seq databases (e.g. UCSC Cell Browser [7] and PanglaoDB [8]), and a majority of processed objects are provided by Zenodo [9], Human Cell Atlas (HCA) [10], and CELLxGENE [11]. The diversity of data types and their distribution in specialized repositories make scRNA-seq data download difficult. Secondly, as many scRNA-seq protocols and preprocessing tools have been developed, the data they generated and stored differ. Representatively, there are differences between 10x Genomics (10x) [12] and Smart-seq2 [13] in terms of sequencing reads and output count matrix form. How to process and load these different types of downloaded files into a unified object that can be analyzed matter. Thirdly, the formats and programming languages of downloaded processed objects are varied due to the prosperity in scRNA-seq analysis tools, such as *SeuratObject* (*Seurat*, R) [14], *AnnData* (*Scanpy*, Python) [15], *CellDataSet*/*cell_data_set* (*Monocle2*/*3*, R) [16, 17], and *SingleCellExperiment* (*scater*, R) [18]. Diverse object formats and programming languages hinder the integration of scRNA-seq data and the interoperability between different analysis tools, thus the object format conversion is vital.

Currently available tools mainly focus on downloading raw data (SRA and ENA) and count matrices (GEO) (Table 1). *ffq* [19] is designed to fetch sample metadata including the download links, therefore third-party tools are needed to perform downloading. *GEOfastq* [20], *SRA-Toolkit* [21], *enaBrowserTools* [22], *fastq-dl* [23], *pysradb* [24], and *GEOfetch* [25] supports downloading raw data from SRA and/or ENA. Among them, *GEOfastq*, *fastq-dl*, and *GEOfetch* can only download *fastq*, *sra*/*fastq*, and *sra* files respectively, and *GEOfetch* supports splitting *sra* into *fastq*/*bam* files. *GEOfastq*, *enaBrowserTools*, *fastq-dl* (version <1.2.0), and *pysradb* support downloading data via Aspera, while *fastq-dl* and *pysradb* parallelize the download process. *pysradb*, *GEOfetch*, and *GEOquery* [26] are able to download the supplementary files from GEO, and *GEOquery* can extract the count matrix from *ExpressionSet*. *rPanglaoDB* [27] can be used to download scRNA-seq count matrices and annotations from PanglaoDB, which only contains datasets originating from human and mouse, has relatively small sample numbers, and is no longer maintained. The above generalized tools, except *rPanglaoDB*, support downloading raw data and/or count matrices from limited databases. And this download is regardless of whether the data are from scRNA-seq or bulk RNA-seq, thus lacking support for unique characteristics of scRNA-seq data (such as 10x-style *fastq* files and *bam* files with custom tags). *rPanglaoDB* specializes in downloading scRNA-seq count matrices and annotations, but the support is quite insufficient. As for format conversion, many tools have been developed for the same conversion stream. From *SeuratObject* to *AnnData* (Seu2AD), available tools are *SeuratDisk*, *sceasy*, and *scDIOR*, while the above tools and *schard* are suitable for the reverse stream (AD2Seu). From *SingleCellExperiment* to *AnnData* (SCE2AD), available tools are *sceasy*, *scDIOR*, and *zellkonverter*, while *scDIOR*, *zellkonverter*, and *schard* are suitable for the reverse stream (AD2SCE). Multiple available tools for the same stream confused the format conversion process, and currently lacking a benchmark of format conversion tools to guide the tool selection. Besides, no tool can cover all the conversion streams between the widely used scRNA-seq objects.

To address the above issues, we present *GEfetch2R*, an R package to (i) fetch scRNA-seq/bulk RNA-seq raw data, count matrices, and processed objects from diverse public repositories; (ii) process the downloaded data, load the count matrices and annotations to R (*SeuratObject*/*DESeqDataSet*), filter the *SeuratObject* based on cell metadata and genes, and merge multiple *SeuratObjects* if applicable; (iii) convert formats between the widely used scRNA-seq objects, and benchmark format conversion tools to guide the tool selection.

## Materials and Methods

### Download, process and load raw data

With GEO accessions as input, *GEfetch2R* can download raw data (*sra*/*fastq*/*bam*) from SRA and ENA (Supplementary Fig. S1). Firstly, *GEfetch2R* extracts all sample metadata under given GEO accessions, users can skip this step and provide a data frame containing interested samples. The output sample metadata is used as input for subsequent raw data download. For *sra* files, *GEfetch2R* uses *prefetch* command to download them from SRA, and uses *ascp* and *download.file* command in parallel to download them from ENA.

For *fastq* files in SRA, *GEfetch2R* uses *parallel-fastq-dump* (parallel), *fasterq-dump* (parallel), and *fastq-dump* commands to split the downloaded *sra* files into *fastq* files. If the data are from 10x, the *--split-files* parameter is added and at least two *fastq* files are generated. *GEfetch2R* automatically distinguishes read1 and read2 based on read length, and renames them according to the *CellRanger* required format. As for data from other scRNA-seq protocols or bulk RNA-seq, the *--split-3* parameter is used. For *fastq* files in ENA, *GEfetch2R* supports splitting the downloaded *sra* files as above, alternatively, if there are *fastq* files in ENA, *GEfetch2R* uses *ascp* and *download.file* command in parallel to download them directly.

For *bam* files in SRA, if they are from 10x, *GEfetch2R* uses *prefetch* command with *--type TenX* parameter to download the original uploaded *bam* files directly to keep the custom tags required to reconstruct the original *fastq* files. As for *bam* files from other scRNA-seq protocols or bulk RNA-seq, *GEfetch2R* uses *sam-dump* commands to split the downloaded *sra* files into *bam* files. For *bam* files in ENA, *GEfetch2R* uses *ascp* and *download.file* command in parallel to download them directly. If *fastq* files are needed, *GEfetch2R* uses *bamtofastq* command developed by 10x to convert the downloaded 10x-generated *bam* files to *fastq* files, and uses *samtools* [28] command to convert the downloaded *bam* files generated by other scRNA-seq protocols or bulk RNA-seq to *fastq* files.

With prepared *fastq* files, if they are from 10x, *GEfetch2R* uses *CellRanger* to perform read alignment and counting with default parameters. Users can pass custom parameters to *st.paras* parameter in *Fastq2R* function. The output count matrix is then loaded to R using *Seurat*. If multiple count matrices are available, *GEfetch2R* merges multiple *SeuratObjects* if applicable. As for *fastq* files from Smart-seq2 or bulk RNA-seq, *GEfetch2R* uses *STAR* to perform read alignment and counting with *--quantMode GeneCounts* parameter. Users can also pass custom parameters to *st.paras* parameter in *Fastq2R* function. The output count matrix is then loaded to R using *DESeq2*.

### Download, process and load count matrices and annotations

As in Supplementary Fig. S1, *GEfetch2R* first extracts detailed dataset metadata, such as description, source/tissue, organism, protocol, and related publication. Users can filter datasets based on these attributes, and the filtered metadata is used as subsequent input. When fetching count matrices from GEO, *GEfetch2R* tries to extract the count matrix from *ExpressionSet* first. If the extracted count matrix is empty or contains non-integer values, *GEfetch2R* generates the count matrix from supplementary files. For supplementary files in *CellRanger* output format, *GEfetch2R* automatically categorizes the downloaded files based on sample names. For supplementary files composed of the count matrix of every single cell/bulk sample, *GEfetch2R* creates a count matrix containing all cells/samples. Users can also add *down.supp = TRUE* parameter in *ParseGEO* function to generate the count matrix directly from supplementary files. For UCSC Cell Browser, *GEfetch2R* accesses count matrices online instead of downloading them to save time and disk usage. In addition to count matrices, UCSC Cell Browser contains diverse annotations, such as cell type annotation, cell type composition, and dimensionality reduction coordinates, *GEfetch2R* supports extracting all these annotations. For PanglaoDB, similar to UCSC Cell Browser, it also contains various annotations (e.g. cell type annotation and cell type composition), *GEfetch2R* uses *rPanglaoDB* to extract count matrices and corresponding annotations.

With available scRNA-seq count matrices and various annotations, *GEfetch2R* loads the count matrices to R using *Seurat*, adds available annotations to the *SeuratObjects* (UCSC Cell Browser and PanglaoDB), filters the *SeuratObjects* based on cell metadata and genes (UCSC Cell Browser and PanglaoDB), and merges *SeuratObjects* if applicable. For the bulk RNA-seq count matrix containing all samples, *GEfetch2R* loads it to R using *DESeq2* (GEO).

### Download and load processed objects

Similar to downloading count matrices, *GEfetch2R* first extracts detailed dataset metadata (dataset description in Zenodo) (Supplementary Fig. S1). Users can select the dataset of interest based on the dataset information and use it as input for subsequent processed object download. For processed objects in Zenodo, with provided DOIs or selected dataset description table, and the file extensions (e.g. *rds*, *rdata*, *loom*, and *h5ad*), *GEfetch2R* downloads the corresponding processed objects. For processed objects in CELLxGENE, *GEfetch2R* provides two methods to download them. *GEfetch2R* uses *cellxgene-census* [29] developed by the CELLxGENE team to efficiently access the cloud-hosted Census single-cell data and filter the data based on cell metadata and genes. However, some newly uploaded data that can be explored on the web page may be missing due to the delay of cloud hosting (e.g. the stable Census release is 2025-01-30 when the current date is 2025-02-26). To overcome this, *GEfetch2R* takes advantage of CZ CELLxGENE Discover API to get the download links of processed objects (*rds* and *h5ad* files) directly, the same as clicking the download button on the web page. For processed objects in HCA, with filtered metadata and provided file extensions (e.g. *rdata*, *rds*, *h5*, *h5ad*, *loom*, and *tsv*), *GEfetch2R* downloads the corresponding processed objects. With processed objects in *rds* format, *GEfetch2R* loads them (*SeuratObjects*) to R, filters the *SeuratObjects* based on cell metadata and genes (CELLxGENE), and merges *SeuratObjects* if applicable.

### Benchmark format conversion tools

To inspect the information kept after format conversion, we used the pbmc3k dataset analyzed by the standard *Seurat* and *Scanpy* workflow. The *SingleCellExperiment* used to inspect the information kept from *SingleCellExperiment* to *AnnData* was generated by *zellkonverter* (convert *AnnData* to *SingleCellExperiment*). The detailed code and output used to evaluate the retained information are provided in Supplementary Note.

To examine the running time and scalability of format conversion tools, we used *GEfetch2R* to select and download five processed objects (both *rds* and *h5ad* files) with more than one million cells from CELLxGENE (Supplementary Table S1), then subsampled them to varied cell numbers: 1.2 million (1.2M, two objects), 1 million (1M), 800,000 (800K), 600,000 (600K), 500,000 (500K), 300,000 (300K), 200,000 (200K), 100,000 (100K), 80,000 (80K), 60,000 (60K), 50,000 (50K), 30,000 (30K), 20,000 (20K), 10,000 (10K). Each format conversion tool was executed on all the five object sets (64 objects in total). For the same object set, we ensured consistency in the type of count matrices converted between different tools to avoid potential bias. All format conversion tools were executed on a server with one Intel(R) Xeon(R) Gold 5220R CPU, 512GB RAM, and CentOS 7.5.1804 operating system. The running time (elapsed time) was recorded using *system.time* command in R. To ensure the independence of each run, all commands are run in linear order on the Linux shell as R scripts.

### Format conversion between scRNA-seq objects

*GEfetch2R* uses *Seurat* to convert formats between *SeuratObject* and *SingleCellExperiment*, uses *Seurat* and *SeuratDisk* to convert formats between *SeuratObject* and *loom*, uses *Seurat* and *SeuratWrappers* to convert formats between *SeuratObject* and *CellDataSet*/*cell_data_set*, and uses *LoomExperiment* to convert formats between *SingleCellExperiment* and *loom*. For format conversion between *AnnData* and *SeuratObject*/*SingleCellExperiment*, all the tools used for benchmarking are available in *GEfetch2R*.

## Results

### Overview of *GEfetch2R*

*GEfetch2R* supports downloading all the three types of scRNA-seq data: raw data, count matrix, and processed object (Fig. 1). The databases corresponding to each data type, number of datasets/species/cells available, downloaded and returned data, and other key features are summarized in Table 2. For fetching raw data, *GEfetch2R* accepts GEO accessions as input, and downloads *sra*, *fastq*, and *bam* files from SRA and ENA. In particular, the parameters for downloading 10x-related *fastq* and *bam* files have been optimized, such as downloading and formatting the *fastq* files to *CellRanger* required format, and downloading *bam* files with original tags. With downloaded files, *GEfetch2R* processes them and loads the output to R (*SeuratObject* for 10x-generated data, *DESeqDataSet* for Smart-seq2 or bulk RNA-seq data). For fetching count matrices, with GEO accessions (GEO) and key words/filter criteria (UCSC Cell Browser and PanglaoDB) as input, *GEfetch2R* accesses count matrices and various annotations, loads them to R, and extracts subset based on cell metadata and genes (UCSC Cell Browser and PanglaoDB). For fetching processed objects, with DOI (Zenodo) and key words/filter criteria (CELLxGENE and HCA) as input, *GEfetch2R* downloads processed objects in given file extensions, loads *rds* files (SeuratObjects) to R, and extracts subset based on cell metadata and genes (CELLxGENE). The processed objects in other formats can be loaded to R/Python through format conversion.

**Figure 1:**
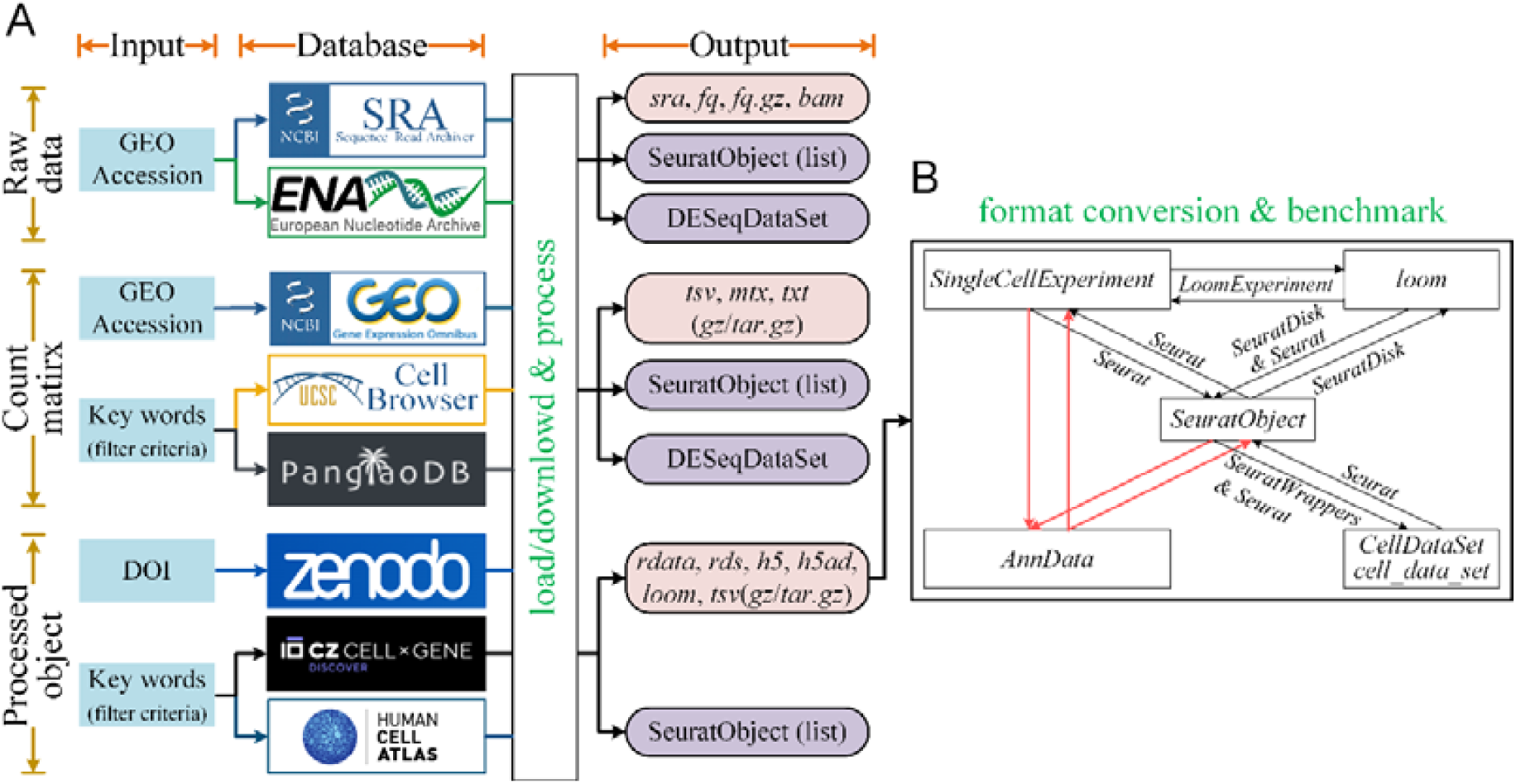
Overview of *GEfetch2R*. (A) Fetching and loading various data from public repositories to R. The download of raw data, count matrix, and processed object are indicated by ochre borders (horizontal). The input, supported databases, and final output are indicated by orange borders (vertical). The boxes in purple indicate R objects, in light pink indicate downloaded file formats. (B) Format conversions supported by *GEfetch2R*. The boxes indicate scRNA-seq objects, the directional arrows between boxes indicate conversion stream between objects, the texts above arrows represent the tool used for the conversion, the red directional arrows without texts indicate conversion streams that are benchmarked.

To enable the interoperability between scRNA-seq analysis tools and the integration of processed objects in diverse formats, *GEfetch2R* supports converting formats (i) between *SeuratObject* and *AnnData*, *SingleCellExperiment*, *CellDataSet*/*cell_data_set*, *loom*; (ii) between *SingleCellExperiment* and *loom*; (iii) between *SingleCellExperiment* and *AnnData*. Moreover, as several tools have been developed for the same conversion stream between *AnnData* and *SeuratObject/SingleCellExperiment*, we benchmarked their performance in terms of information kept, usability, running time, and scalability.

### Application of *GEfetch2R* in COVID-19 scRNA-seq altas exploration

Coronavirus disease 2019 (COVID-19) is a highly contagious disease caused by severe acute respiratory syndrome coronavirus 2, with symptoms ranging from mild to severe [30]. Depicting the dynamic immune responses across symptom severities at the single-cell level is an effective method to understand COVID-19 progression. To demonstrate the utility of *GEfetch2R*, we applied it to download and explore all T cells of a COVID-19 scRNA-seq altas [30] from UCSC Cell Browser. As shown in Fig. 2A, there are twelve T cell subtypes, including six subtypes of CD4^+^ T cells, three subtypes of CD8^+^ T cells, and three subtypes of natural killer T (NKT) cells. The cell type composition analysis across four conditions (healthy donor (HD), moderate, severe, and convalescent (conv)) revealed that compared with HDs, the proportions of four CD4^+^ T subtypes (CD4^+^ naive, CD4^+^ memory, CD4^+^ effector memory, and regulatory T (Treg)) and CD8^+^ naive subtype were significantly decreased in COVID-19 patients, and the proportion of CD4*^+^* naive subtype was still significantly reduced in COVID-19 conv samples (Fig. 2B-C). Meanwhile, the proportions of CD4^+^ effector-GNLY, CD8^+^ effector-GNLY, NKT CD56, and NKT CD160 subtypes were significantly increased in COVID-19 patients, and the CD4^+^ effector-GNLY subtype was nearly absent in HDs but highly enriched in moderate, severe and conv samples (Fig. 2B-C). The apoptosis scoring results showed that T cells in COVID-19 patients generally had significantly increased apoptosis scores (Fig. 2D).

**Figure 2:**
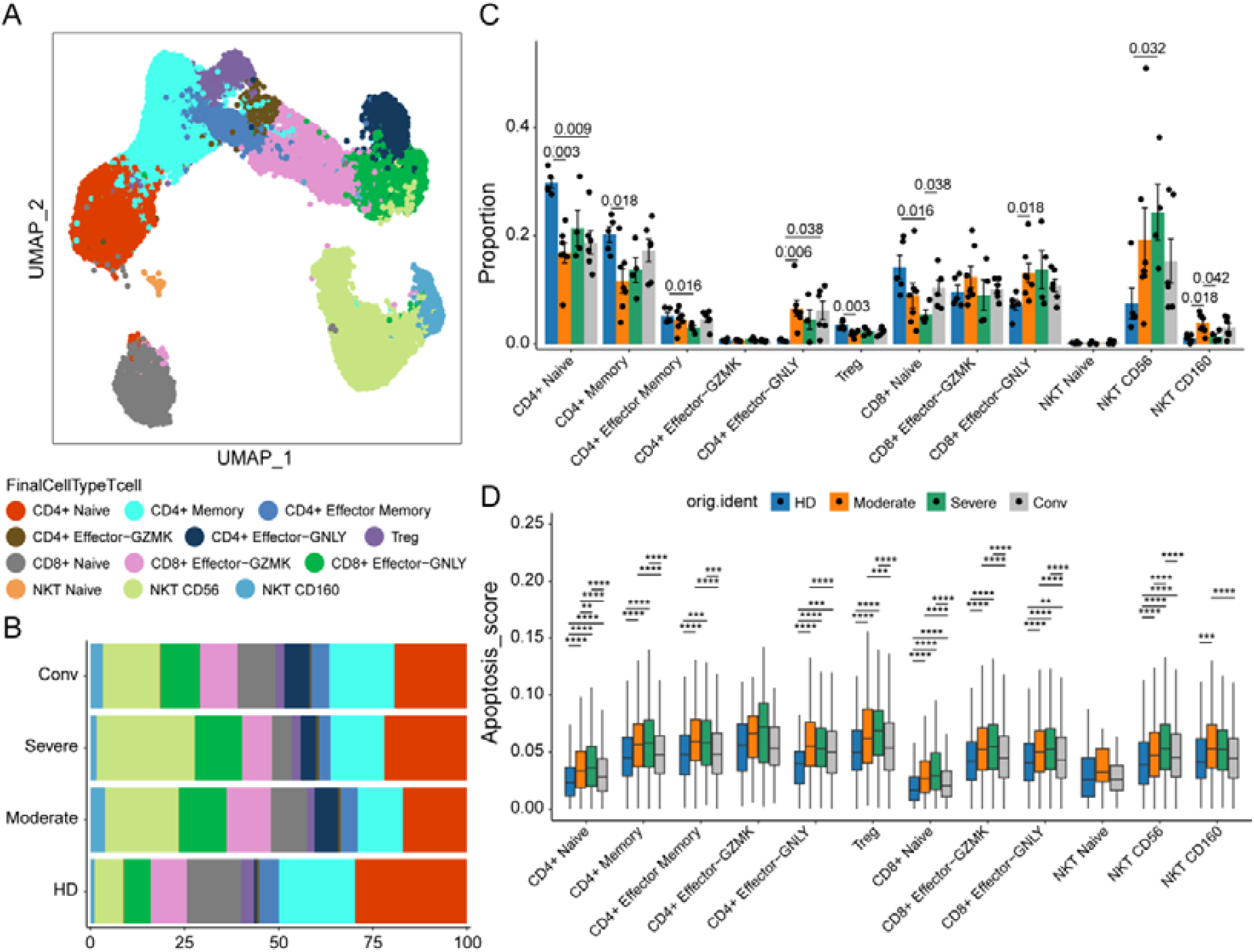
Application of *GEfetch2R* in COVID-19 scRNA-seq altas exploration. (A) UMAP embedding of all T cells. Clusters in different colors represent T cell subtypes. (B) The proportion of each cell type across HD, moderate, severe, and conv samples. (C) Differential cell type composition analysis. All differences with *P*LJ<LJ0.05 are labeled; two-sided unpaired Mann–Whitney *U*-test was used for analysis. (D) The apoptosis scoring results. All differences with Bonferroni adjusted *P*LJ<LJ0.01 are indicated. \*\**P*LJ<LJ0.01; \*\*\**P*LJ<LJ0.001; \*\*\*\**P*LJ<LJ0.0001; using two-sided unpaired Dunn’s test. NKT: natural killer T; Treg: regulatory T.

### Benchmark of format conversion tools Information kept

We evaluated and ranked the information kept of format conversion tools in terms of count matrix and annotation. Figure 3A and Supplementary Note show our rank reasoning and the overall information kept rank of each tool. In Seu2AD, *scDIOR* can preserve all the three count matrices and more annotations, thus ranking first. *SeuratDisk* can keep two of the three count matrices and more annotations, thus ranking second. While *sceasy* keeps only one count matrix and the least annotations, thus ranking last. In SCE2AD, *zellkonverter* and *scDIOR* both keep all the three count matrices, while *zellkonverter* preserves the most comprehensive annotations, thus *zellkonverter* ranks first. *sceasy* keeps only one count matrix and less comprehensive annotations, resulting in the lowest information kept rank. In AD2Seu, *scDIOR*, *sceasy*, and *SeuratDisk* can keep two of the three count matrices, which is more than *schard*. And, *scDIOR* preserves the most comprehensive annotations, thus ranking first. Both *schard* and *sceasy* retain the least annotations, thus *schard* ranks last. In AD2SCE, *zellkonverter* can preserve two of the three count matrices and the most comprehensive annotations, thus ranking first. While both *schard* and *scDIOR* keep only one count matrix, and *schard* retains the least annotations, thus *schard* ranks last.

**Figure 3:**
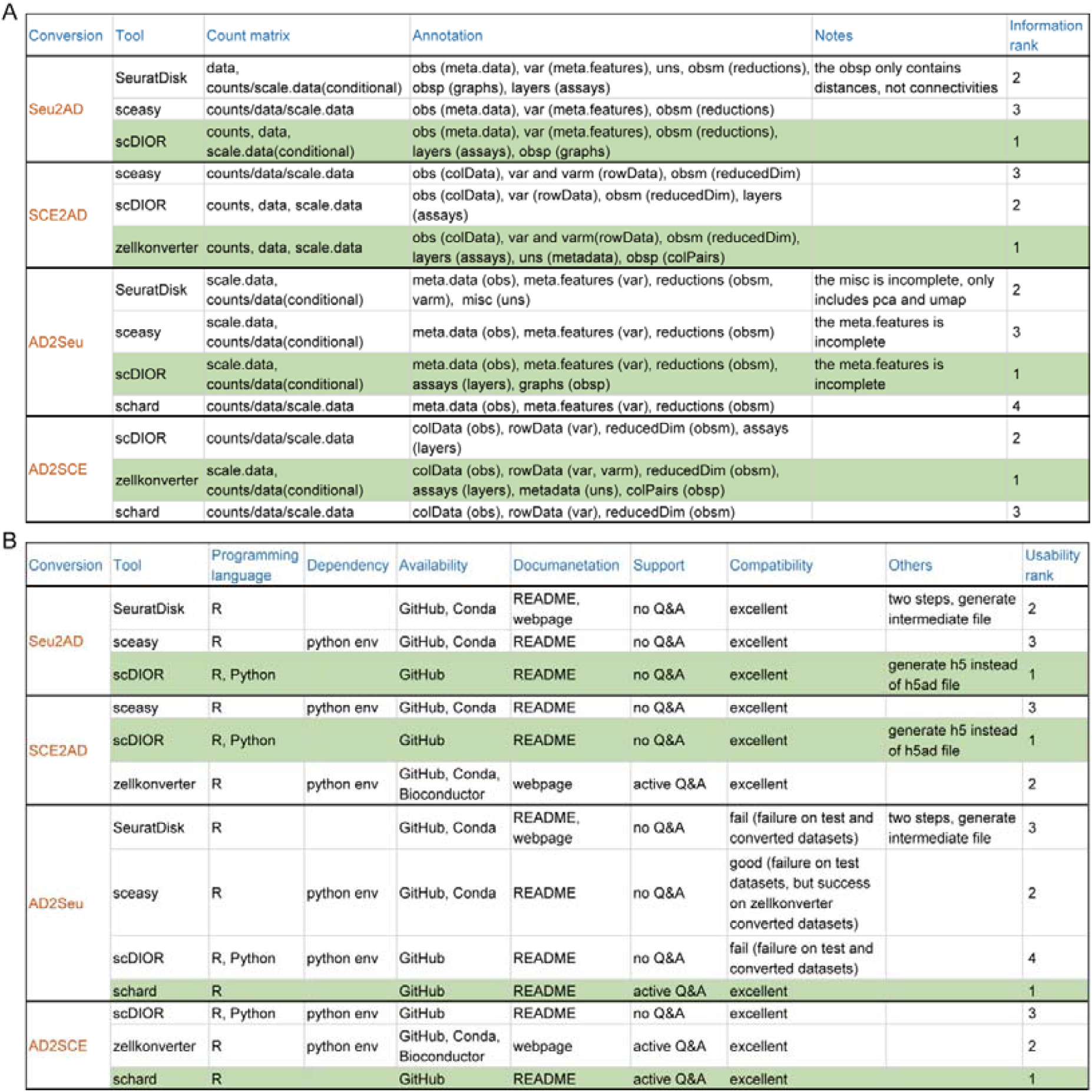
The information kept and usability of format conversion tools. (A) The information kept of format conversion tools in terms of count matrix and annotation. In ‘Count maatrix’ column, ‘counts’ represents raw count matrix, ‘data’ represents normalized data matrix, ‘scale.data’ represents scaled data matrix. In ‘Annotation’ column, the text outside/inside parentheses represents the data slot of the source/target object. (B) The usability of format conversion tools in terms of programming language, dependency, availability, documentation, support, and compatibility. Q&A: question and answer. The tool with the highest rank is marked in green.

### Usability

We evaluated and ranked the usability of format conversion tools in terms of programming language, dependency, availability, documentation, support, and compatibility. Fig. 3B shows our rank reasoning and the overall usability rank of each tool. In Seu2AD, *sceasy* requires an additional Python environment, resulting in the lowest usability rank. *SeuratDisk* and *scDIOR* have comparable usability, but *scDIOR* can be used in both R and Python platforms, thus having a higher usability rank. In SCE2AD, *scDIOR* is the only tool that doesn’t require an additional Python environment and can be used in both R and Python platforms, resulting in the highest usability rank. *zellkonverter* is hosted on Bioconductor and has active questions and answers (Q&As), thus ranking second. In AD2Seu, *schard* doesn’t require an additional Python environment, has active Q&As, and is the only tool that runs successfully on all test data. Among the remaining three tools, *sceasy* runs successfully on *zellkonverter* converted *AnnData*, while the other tools still fail. Based on the above results, *schard* ranks first and *sceasy* ranks second in usability, and the subsequent running time is recorded on *zellkonverter* converted *AnnData*. In AD2SCE, *schard* has active Q&As and is the only tool that doesn’t require an additional Python environment, resulting in the highest usability rank. *zellkonverter* and *scDIOR* have comparable usability, but the former is slightly better. In detail, *zellkonverter* is hosted on Bioconductor and has active Q&As, while *scDIOR* can be used in both R and Python platforms.

### Running time and scalability

In Supplementary Table S2, we summarized the running time of the format conversion tools for converting between *AnnData* and *SeuratObject*/*SingleCellExperiment* on five object sets (64 objects in total). Figure 4 shows that in AD2SCE, *schard* is always the fastest tool across all the five object sets, while *zellkonverter* is the slowest tool in two of the five object sets. In AD2Seu, when the cell number is smaller than 200K, *schard* is faster than *sceasy* or has comparable speed, but *sceasy* is significantly faster than *schard* when the cell number exceeds 200K. This alternation means that *sceasy* has better scalability than *schard*. In SCE2AD, *zellkonverter* is always the fastest tool across all the five object sets, while *sceasy* and *scDIOR* are the slowest tools in two of the five object sets. In Seu2AD, across all the five object sets, *sceasy* is always the fastest tool, while *SeuratDisk* is always the slowest tool.

**Figure 4:**
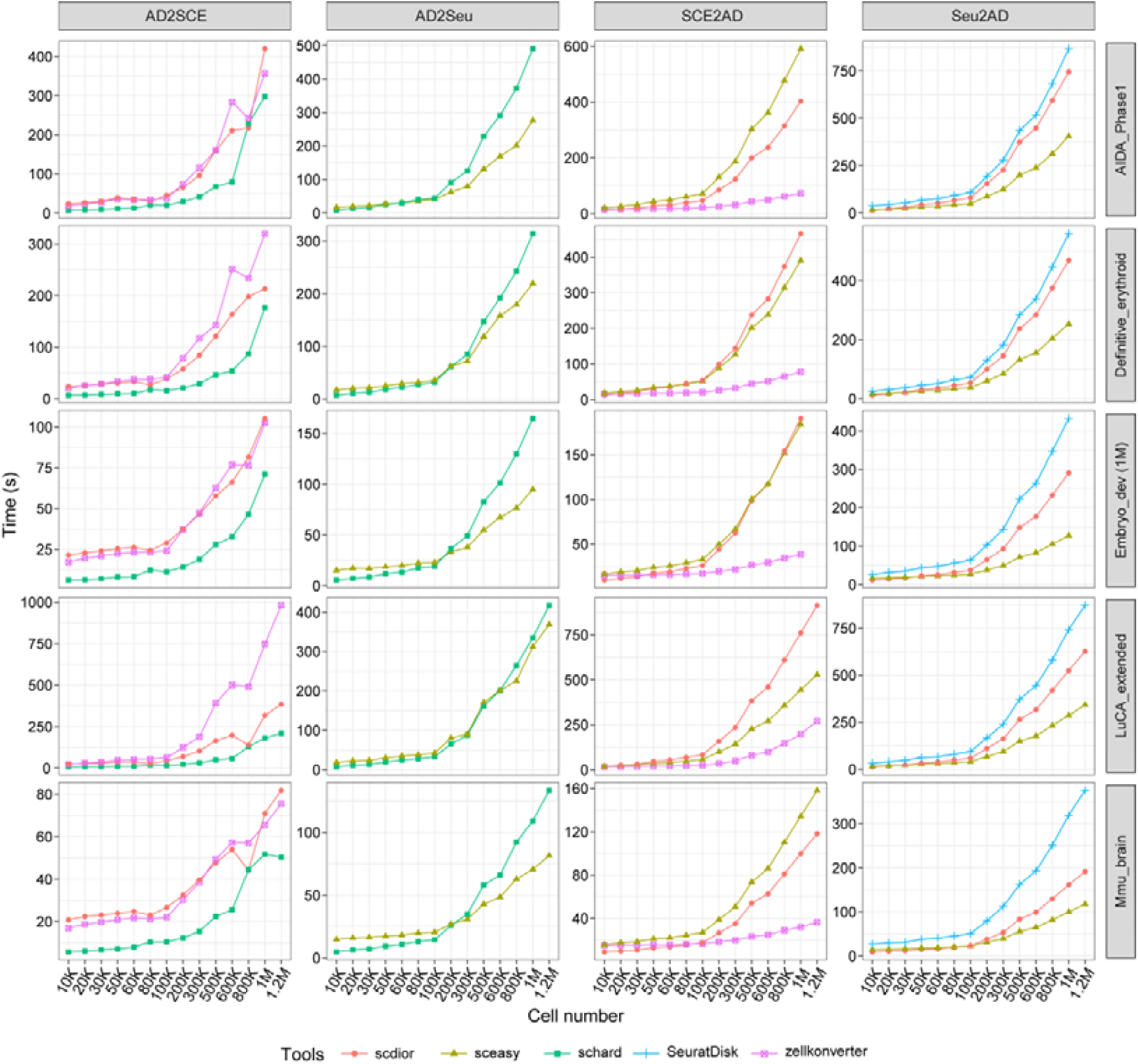
The running time of format conversion tools. There are twenty subplots, distributed in four rows and five columns. Each row represents the format conversion tools are executed on the same object set. Each column represents the same conversion stream. x-axis means the cell number (K: thousand, M: million), *y*-axis means the running time.

### Comparison of *GEfetch2R* with other similar tools

Table 1 shows a feature comparison of *GEfetch2R* with other tools that are capable of downloading data from public repositories. *GEfetch2R* distinguishes itself in the following aspects. Firstly, *GEfetch2R* supports downloading the most diverse data types (raw data, count matrix, and processed objects), and each data type can be downloaded from multiple public repositories. This benefits users in collecting data more comprehensively and choosing appropriate download data types according to different needs. For quick data exploration, downloading processed objects is a good choice, whereas for fine data integration to draw conclusions, downloading raw data or count matrices can help reduce the bias. Secondly, *GEfetch2R* adopts multiple methods to speed up downloading, including Aspera support, parallel downloading *fastq* files, and parallel splitting *sra* files into *fastq* files. Thirdly, *GEfetch2R* supports downloading data of both scRNA-seq and bulk RNA-seq, and considers the characteristics of data generated by different protocols. For 10x-generated data, *GEfetch2R* downloads and formats the *fastq* files to *CellRanger* required format, downloads *bam* files with original tags, converts *bam* files to *fastq* files using *bamtofastq*, and aligns *fastq* files to the reference genome using *CellRanger*. For Smart-seq2 and bulk RNA-seq data, *GEfetch2R* converts *bam* files to *fastq* files using *samtools*, and aligns *fastq* files to the reference genome using *STAR*. This expands the scope of use of *GEfetch2R* and makes the data it downloads easier to use. Fourthly, besides data download, *GEfetch2R* can process the downloaded data, load output/downloaded count matrices and annotations to R (*SeuratObject*/*DESeqDataSet*), extract a subset of the *SeuratObject* based on cell metadata and genes, and merge multiple *SeuratObjects* if applicable. This greatly distinguishes *GEfetch2R* from other tools, which are pure data downloaders. Lastly, *GEfetch2R* provides the most comprehensive object format conversions, and benchmarks the format conversion tools for converting between *AnnData* and *SeuratObject*/*SingleCellExperiment*. This enables users to integrate multiple objects in diverse formats, bridges the widely used scRNA-seq analysis tools, and guides the selection of format conversion tools.

## Conclusions and Discussions

*GEfetch2R* is an R package dedicated to facilitating researchers to access and explore the existing gene expression data from various public repositories. As a data downloader, *GEfetch2R* supports downloading the most diverse scRNA-seq data types, including raw data (SRA and ENA), count matrices (GEO, UCSC Cell Browser, and PanglaoDB), and processed objects (Zenodo, CELLxGENE, and HCA). Besides the data download ability, *GEfetch2R* can process the downloaded data, load output/downloaded count matrices and annotations to R (*SeuratObject*/*DESeqDataSet*), extract a subset of the *SeuratObject* based on cell metadata and genes, and merge multiple *SeuratObjects* if applicable. Furthermore, to enable the integration of scRNA-seq data and the interoperability between different analysis tools, *GEfetch2R* provides the most comprehensive format conversions between different scRNA-seq objects, including *SeuratObject*, *AnnData*, *SingleCellExperiment*, *CellDataSet*/*cell_data_set*, and *loom*. In particular, *GEfetch2R* benchmarks the format conversion tools for converting between *SeuratObject*/*SingleCellExperiment* and *AnnData*. In Seu2AD, *scDIOR* has the best performance in terms of information kept and usability, and *sceasy* is always the fastest tool. In SCE2AD, *zellkonverter* has the highest information kept rank, *scDIOR* is the best in usability, and *zellkonverter* is consistently the fastest tool. In AD2Seu, *scDIOR* has the best performance in terms of information kept, *schard* is the best in usability, and *sceasy* has better scalability than *schard*. In AD2SCE, *zellkonverter* has the highest information kept rank, *schard* is the best in terms of usability and speed.

In addition to fetching scRNA-seq data, *GEfetch2R* can also be used to download raw data (SRA and ENA) and count matrices (GEO) of bulk RNA-seq, process the downloaded data, and load output/downloaded count matrices to R (*DESeqDataSet*). Currently, there are still several aspects of *GEfetch2R* that can be improved. Firstly, *GEfetch2R* only supports processing the Smart-seq2 and 10x-generated raw data, users can download the raw data generated by other scRNA-seq protocols using *GEfetch2R*, but the subsequent process is not available. Secondly, many widely used scRNA-seq databases are not supported by *GEfetch2R*, such as Single Cell Portal [31]. We will actively develop to support more scRNA-seq protocols and databases. Besides, since the APIs of supported databases may change, we will update *GEfetch2R* on time to ensure usage.

## Availability and Requirements

- Project name: *GEfetch2R*
- Project homepage: https://github.com/showteeth/GEfetch2R
- Software documentation: https://showteeth.github.io/GEfetch2R
- Docker image: https://hub.docker.com/r/soyabean/gefetch2r
- Operating system(s): Platform independent
- Programming language: R
- Other requirements: R 2.10 or higher, Python 3.7.3 or higher (format conversion)
- License: GPL-3.0
- RRID: SCR_026714

## Additional Files

Supplementary Note. Step-by-step code and output showing information kept of format conversion tools.

Supplementary Fig. S1. The detailed workflow of *GEfetch2R*.

Supplementary Table S1. The datasets used for benchmarking format conversion tools.

Supplementary Table S2. The running time of the format conversion tools on five object sets.

## Abbreviations

scRNA-seq: single-cell RNA sequencing
SRA: Sequence Read Archive
ENA: European Nucleotide Archive
GEO: Gene Expression Omnibus
HCA: Human Cell Atlas
10x: 10x Genomics
Seu2AD: from SeuratObject to AnnData
AD2Seu: from AnnData to SeuratObject
SCE2AD: from SingleCellExperiment to AnnData
AD2SCE: from AnnData to SingleCellExperiment
K: thousand
M: million
COVID-19: Coronavirus disease 2019
NKT: natural killer T
HD: healthy donor
conv: convalescent
Treg: regulatory T
Q&As: questions and answers

## Acknowledgements

We thank SRA, ENA, GEO, PanglanDB, UCSC Cell Browser, Zenodo, CELLxGENE, and HCA for generously hosting the single-cell/bulk RNA-seq data and providing websites/APIs for users to access these data, and gratefully acknowledge all the data contributors. These are the cornerstones for the development of *GEfetch2R*.

## Author Contributions

Y.S., J.W., and J.G. contributed to the conception and design of *GEfetch2R*. Y.S. implemented and benchmarked *GEfetch2R*. J.W. and J.G. provided supervision and secured funding. All authors prepared and approved the manuscript.

## Data Availability

The datasets used in this study are freely available from SRA, CELLxGENE, and *SeuratData* [32]. All scripts used for the COVID-19 scRNA-seq altas exploration and benchmark of format conversion tools, including downloading, processing,

subsampling, running, and visualizing, are available in the GitHub repository [33]. Snapshots of the code and data are available in Software Heritage [34].

---

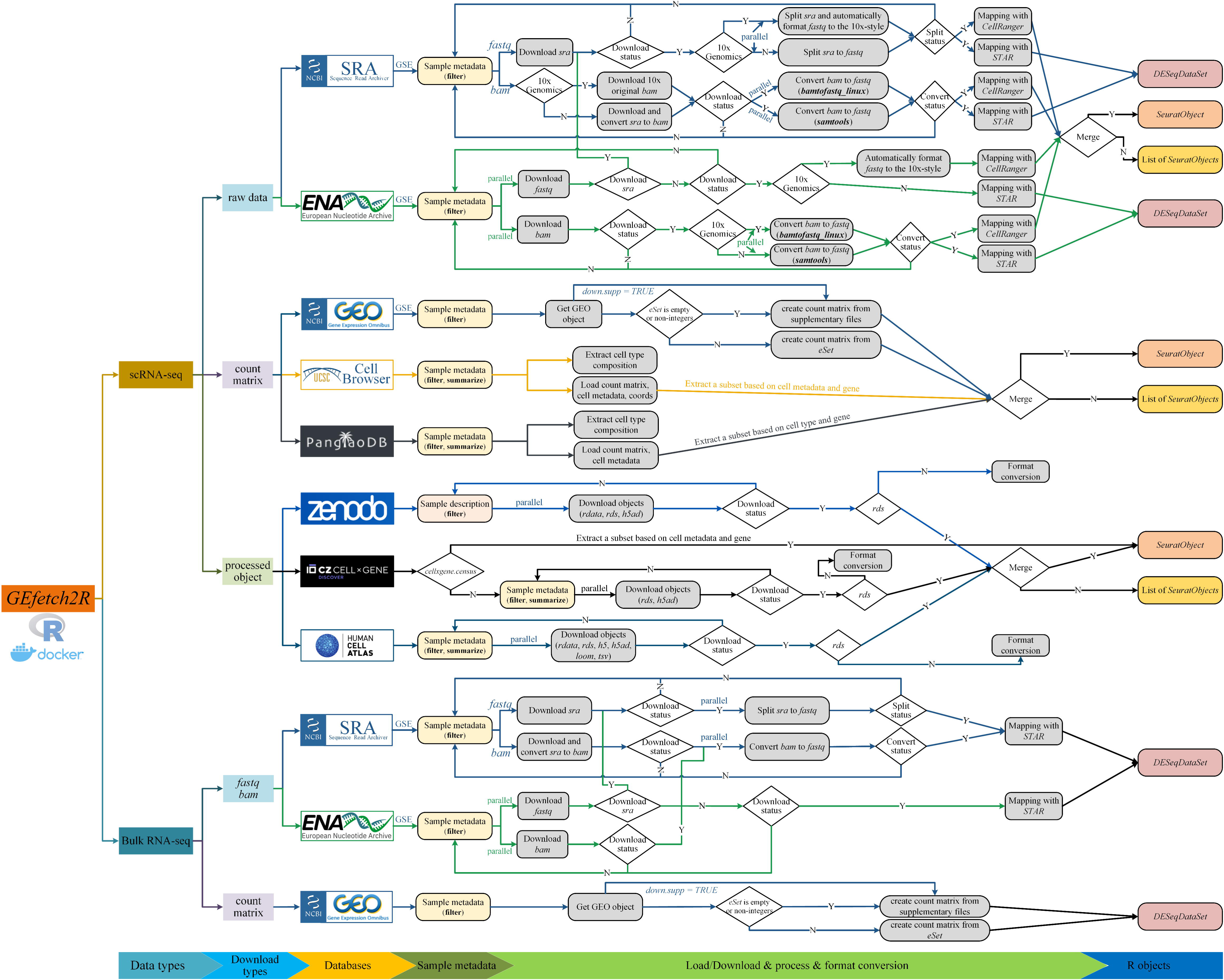

