## Supplementary Note for "*GEfetch2R*: fetching single-cell/bulk RNA-seq data from public repositories to R and benchmarking the subsequent format conversion tools": Supplementary Note Step-by-step code and output showing information kept of format conversion tools.html

GEfetch2R\_benchmark


### Library and configuration¶

In [2]:

```
# read .h5ad files
import scanpy as sc

# setup R environment
import os
os.environ["R_HOME"] = r"/Library/Frameworks/R.framework/Resources" 
# !pip install rpy2==3.5.1
# enables the %%R magic
%load_ext rpy2.ipython

# for display
from IPython.core.interactiveshell import InteractiveShell
InteractiveShell.ast_node_interactivity = "all"
from IPython.display import display, HTML
display(HTML("<style>.container { width:83%; align: left; }</style>"))
display(HTML("<style>#toc-wrapper{ position: relative; width: 20%; top: 130px; left: 0px; }</style>"))
```

```
The rpy2.ipython extension is already loaded. To reload it, use:
  %reload_ext rpy2.ipython
```

### SeuratObject to AnnData¶

#### SeuratObject¶

In [3]:

```
%%R
library(Seurat)
library(tidyverse)
library(SeuratData) # load pbmc3k.final
data('pbmc3k.final')
```

```
    WARNING: The R package "reticulate" does not
    consider that it could be called from a Python process. This
    results in a quasi-obligatory segfault when rpy2 is evaluating
    R code using it. On the hand, rpy2 is accounting for the
    fact that it might already be running embedded in a Python
    process. This is why:
    - Python -> rpy2 -> R -> reticulate: crashes
    - R -> reticulate -> Python -> rpy2: works

    The issue with reticulate is tracked here:
    https://github.com/rstudio/reticulate/issues/208
```

```
R[write to console]: Attaching SeuratObject
```

```
── Attaching core tidyverse packages ───────────────────────────────────────────────────────────── tidyverse 2.0.0 ──
✔ dplyr     1.1.2     ✔ readr     2.1.4
✔ forcats   1.0.0     ✔ stringr   1.5.0
✔ ggplot2   3.4.2     ✔ tibble    3.2.1
✔ lubridate 1.9.2     ✔ tidyr     1.3.0
✔ purrr     1.0.1     
── Conflicts ─────────────────────────────────────────────────────────────────────────────── tidyverse_conflicts() ──
✖ dplyr::filter() masks stats::filter()
✖ dplyr::lag()    masks stats::lag()
ℹ Use the ]8;;http://conflicted.r-lib.org/conflicted package]8;; to force all conflicts to become errors
```

```
R[write to console]: Using cached data manifest, last updated at 2024-06-06 17:43:46

R[write to console]: ── Installed datasets ────────────────────────────────────────────────────────────────────────── SeuratData v0.2.2 ──

R[write to console]: ✔ ifnb   3.1.0                                            ✔ pbmc3k 3.1.4


R[write to console]: ──────────────────────────────────────────────────────── Key ────────────────────────────────────────────────────────

R[write to console]: ✔ Dataset loaded successfully
❯ Dataset built with a newer version of Seurat than installed
❓ Unknown version of Seurat installed
```

In [4]:

```
%%R
# read data
pbmc3k.final.counts = Seurat::GetAssayData(object =  pbmc3k.final[['RNA']], slot = 'counts')
pbmc3k.final[["rawcounts"]] <- Seurat::CreateAssayObject(counts = pbmc3k.final.counts )
pbmc3k.final
```

```
An object of class Seurat 
27428 features across 2638 samples within 2 assays 
Active assay: RNA (13714 features, 2000 variable features)
 1 other assay present: rawcounts
 2 dimensional reductions calculated: pca, umap
```

Detailed information about The Seurat Class

##### Count matrix (`assays` and `slot`)¶

In [5]:

```
# for display
from IPython.core.interactiveshell import InteractiveShell
InteractiveShell.ast_node_interactivity = "all"
```

In [6]:

```
%%R
# raw count matrix
pbmc3k.final@assays$RNA@counts[1:10,1:15]
```

```
10 x 15 sparse Matrix of class "dgCMatrix"
```

```
R[write to console]:   [[ suppressing 15 column names ‘AAACATACAACCAC’, ‘AAACATTGAGCTAC’, ‘AAACATTGATCAGC’ ... ]]
```

```
AL627309.1    . . . . . . . . . . . . . . .
AP006222.2    . . . . . . . . . . . . . . .
RP11-206L10.2 . . . . . . . . . . . . . . .
RP11-206L10.9 . . . . . . . . . . . . . . .
LINC00115     . . . . . . . . . . . . . . .
NOC2L         . . . . . . . . . . . 1 . . .
KLHL17        . . . . . . . . . . . . . . .
PLEKHN1       . . . . . . . . . . . . . . .
RP11-54O7.17  . . . . . . . . . . . . . . .
HES4          . . . . . . . . . . . . . . .
```

In [7]:

```
%%R
# shape
dim(pbmc3k.final@assays$RNA@counts)
```

```
[1] 13714  2638
```

In [8]:

```
%%R
# log-normalized count matrix
pbmc3k.final@assays$RNA@data[1:10,1:15]
```

```
10 x 15 sparse Matrix of class "dgCMatrix"
```

```
R[write to console]:   [[ suppressing 15 column names ‘AAACATACAACCAC’, ‘AAACATTGAGCTAC’, ‘AAACATTGATCAGC’ ... ]]
```

```
AL627309.1    . . . . . . . . . . . .        . . .
AP006222.2    . . . . . . . . . . . .        . . .
RP11-206L10.2 . . . . . . . . . . . .        . . .
RP11-206L10.9 . . . . . . . . . . . .        . . .
LINC00115     . . . . . . . . . . . .        . . .
NOC2L         . . . . . . . . . . . 1.646272 . . .
KLHL17        . . . . . . . . . . . .        . . .
PLEKHN1       . . . . . . . . . . . .        . . .
RP11-54O7.17  . . . . . . . . . . . .        . . .
HES4          . . . . . . . . . . . .        . . .
```

In [9]:

```
%%R
# shape
dim(pbmc3k.final@assays$RNA@data)
```

```
[1] 13714  2638
```

In [10]:

```
%%R
# scaled count matrix
pbmc3k.final@assays$RNA@scale.data[1:10,1:4]
```

```
              AAACATACAACCAC AAACATTGAGCTAC AAACATTGATCAGC AAACCGTGCTTCCG
AL627309.1       -0.05812316    -0.05812316    -0.05812316    -0.05812316
AP006222.2       -0.03357571    -0.03357571    -0.03357571    -0.03357571
RP11-206L10.2    -0.04166819    -0.04166819    -0.04166819    -0.04166819
RP11-206L10.9    -0.03364562    -0.03364562    -0.03364562    -0.03364562
LINC00115        -0.08223981    -0.08223981    -0.08223981    -0.08223981
NOC2L            -0.31717081    -0.31717081    -0.31717081    -0.31717081
KLHL17           -0.05344722    -0.05344722    -0.05344722    -0.05344722
PLEKHN1          -0.05082183    -0.05082183    -0.05082183    -0.05082183
RP11-54O7.17     -0.03308805    -0.03308805    -0.03308805    -0.03308805
HES4             -0.23376818    -0.23376818    -0.23376818    -0.23376818
```

In [11]:

```
%%R
# shape
dim(pbmc3k.final@assays$RNA@scale.data)
```

```
[1] 13714  2638
```

In [12]:

```
%%R
# raw count matrix in rawcounts assay
pbmc3k.final@assays$rawcounts@counts[1:10,1:15]
```

```
10 x 15 sparse Matrix of class "dgCMatrix"
```

```
R[write to console]:   [[ suppressing 15 column names ‘AAACATACAACCAC’, ‘AAACATTGAGCTAC’, ‘AAACATTGATCAGC’ ... ]]
```

```
AL627309.1    . . . . . . . . . . . . . . .
AP006222.2    . . . . . . . . . . . . . . .
RP11-206L10.2 . . . . . . . . . . . . . . .
RP11-206L10.9 . . . . . . . . . . . . . . .
LINC00115     . . . . . . . . . . . . . . .
NOC2L         . . . . . . . . . . . 1 . . .
KLHL17        . . . . . . . . . . . . . . .
PLEKHN1       . . . . . . . . . . . . . . .
RP11-54O7.17  . . . . . . . . . . . . . . .
HES4          . . . . . . . . . . . . . . .
```

In [13]:

```
%%R
# shape
dim(pbmc3k.final@assays$rawcounts@counts)
```

```
[1] 13714  2638
```

##### `meta.data` - cells' meta-information¶

In [14]:

```
%%R
# cell's meta-information
pbmc3k.final@meta.data %>% head()
```

```
               orig.ident nCount_RNA nFeature_RNA seurat_annotations percent.mt
AAACATACAACCAC     pbmc3k       2419          779       Memory CD4 T  3.0177759
AAACATTGAGCTAC     pbmc3k       4903         1352                  B  3.7935958
AAACATTGATCAGC     pbmc3k       3147         1129       Memory CD4 T  0.8897363
AAACCGTGCTTCCG     pbmc3k       2639          960         CD14+ Mono  1.7430845
AAACCGTGTATGCG     pbmc3k        980          521                 NK  1.2244898
AAACGCACTGGTAC     pbmc3k       2163          781       Memory CD4 T  1.6643551
               RNA_snn_res.0.5 seurat_clusters nCount_rawcounts
AAACATACAACCAC               1               1             2419
AAACATTGAGCTAC               3               3             4903
AAACATTGATCAGC               1               1             3147
AAACCGTGCTTCCG               2               2             2639
AAACCGTGTATGCG               6               6              980
AAACGCACTGGTAC               1               1             2163
               nFeature_rawcounts
AAACATACAACCAC                779
AAACATTGAGCTAC               1352
AAACATTGATCAGC               1129
AAACCGTGCTTCCG                960
AAACCGTGTATGCG                521
AAACGCACTGGTAC                781
```

##### `meta.features` - annotation of features¶

In [15]:

```
%%R
# annotation of features
pbmc3k.final@assays$RNA@meta.features %>% head()
```

```
                 vst.mean vst.variance vst.variance.expected
AL627309.1    0.003411676  0.003401325           0.003645407
AP006222.2    0.001137225  0.001136363           0.001144957
RP11-206L10.2 0.001895375  0.001892500           0.001965766
RP11-206L10.9 0.001137225  0.001136363           0.001144957
LINC00115     0.006823351  0.006779363           0.007480978
NOC2L         0.107278241  0.159514698           0.203221328
              vst.variance.standardized vst.variable
AL627309.1                    0.9330441        FALSE
AP006222.2                    0.9924937        FALSE
RP11-206L10.2                 0.9627290        FALSE
RP11-206L10.9                 0.9924937        FALSE
LINC00115                     0.9062135        FALSE
NOC2L                         0.7849309        FALSE
```

##### `reductions` - dimensional reduction results and feature loadings¶

In [16]:

```
%%R
# dimensional reduction results of pca
pbmc3k.final@reductions$pca@cell.embeddings[1:5, 1:5]
```

```
                     PC_1       PC_2       PC_3       PC_4        PC_5
AAACATACAACCAC -4.7296855 -0.5184265 -0.7623220 -2.3156790 -0.07160006
AAACATTGAGCTAC -0.5174029  4.5918957  5.9091921  6.9118856 -1.96243034
AAACATTGATCAGC -3.1891063 -3.4695154 -0.8313710 -2.0019985 -5.10442765
AAACCGTGCTTCCG 12.7933021  0.1007166  0.6310221 -0.3687338  0.21838204
AAACCGTGTATGCG -3.1288078 -6.3481412  1.2507776  3.0191026  7.84739502
```

In [17]:

```
%%R
# pca feature loadings
pbmc3k.final@reductions$pca@feature.loadings[1:5, 1:5]
```

```
               PC_1        PC_2        PC_3        PC_4        PC_5
PPBP    0.010990202  0.01148426 -0.15176092  0.10403737 0.003299077
LYZ     0.116231706  0.01472515 -0.01280613 -0.04414540 0.049906881
S100A9  0.115414362  0.01895146 -0.02368853 -0.05787777 0.085382309
IGLL5  -0.007987473  0.05454239  0.04901533  0.06694722 0.004603231
GNLY   -0.015238762 -0.13375626  0.04101340  0.06912322 0.104558611
```

In [18]:

```
%%R
# dimensional reduction results of umap
pbmc3k.final@reductions$umap@cell.embeddings[1:5, 1:2]
```

```
                  UMAP_1    UMAP_2
AAACATACAACCAC -4.232792 -4.152139
AAACATTGAGCTAC -4.892886 10.985685
AAACATTGATCAGC -5.508639 -7.211088
AAACCGTGCTTCCG 11.332233  3.161727
AAACCGTGTATGCG -7.450703  1.092022
```

##### `graphs` - relationship of cells, graphs¶

In [19]:

```
%%R
# RNA_nn
pbmc3k.final@graphs$RNA_nn[1:10,1:10]
```

```
10 x 10 sparse Matrix of class "dgCMatrix"
```

```
R[write to console]:   [[ suppressing 10 column names ‘AAACATACAACCAC’, ‘AAACATTGAGCTAC’, ‘AAACATTGATCAGC’ ... ]]
```

```
AAACATACAACCAC 1 . . . . . . . . .
AAACATTGAGCTAC . 1 . . . . . . . .
AAACATTGATCAGC . . 1 . . . . . . .
AAACCGTGCTTCCG . . . 1 . . . . . .
AAACCGTGTATGCG . . . . 1 . . . . .
AAACGCACTGGTAC . . . . . 1 . . . .
AAACGCTGACCAGT 1 . . . . . 1 . . .
AAACGCTGGTTCTT . . . . . . . 1 . .
AAACGCTGTAGCCA . . . . . . . . 1 .
AAACGCTGTTTCTG . . . . . . . . . 1
```

In [20]:

```
%%R
# RNA_snn
pbmc3k.final@graphs$RNA_snn[1:10,1:10]
```

```
10 x 10 sparse Matrix of class "dgCMatrix"
```

```
R[write to console]:   [[ suppressing 10 column names ‘AAACATACAACCAC’, ‘AAACATTGAGCTAC’, ‘AAACATTGATCAGC’ ... ]]
```

```
AAACATACAACCAC 1.0000000 . . . . . 0.11111111 .          . .
AAACATTGAGCTAC .         1 . . . . .          .          . .
AAACATTGATCAGC .         . 1 . . . .          .          . .
AAACCGTGCTTCCG .         . . 1 . . .          .          . .
AAACCGTGTATGCG .         . . . 1 . .          .          . .
AAACGCACTGGTAC .         . . . . 1 .          .          . .
AAACGCTGACCAGT 0.1111111 . . . . . 1.00000000 0.08108108 . .
AAACGCTGGTTCTT .         . . . . . 0.08108108 1.00000000 . .
AAACGCTGTAGCCA .         . . . . . .          .          1 .
AAACGCTGTTTCTG .         . . . . . .          .          . 1
```

##### `misc` - miscellaneous information¶

In [21]:

```
%%R
# miscellaneous information
pbmc3k.final@misc
```

```
list()
```

##### `commands` - logged commands¶

In [22]:

```
%%R
# logged commands run on this Seurat object
names(pbmc3k.final@commands)
```

```
[1] "NormalizeData.RNA"        "FindVariableFeatures.RNA"
[3] "ScaleData.RNA"            "RunPCA.RNA"              
[5] "JackStraw.RNA.pca"        "ScoreJackStraw"          
[7] "FindNeighbors.RNA.pca"    "FindClusters"            
[9] "RunUMAP.RNA.pca"
```

In [23]:

```
%%R
# parameters for SNN Graph Construction
pbmc3k.final@commands$FindNeighbors.RNA.pca
```

```
Command: FindNeighbors(pbmc3k.final, dims = 1:10)
Time: 2020-04-30 12:54:51
reduction : pca 
dims : 1 2 3 4 5 6 7 8 9 10 
assay : RNA 
k.param : 20 
compute.SNN : TRUE 
prune.SNN : 0.06666667 
nn.method : rann 
annoy.metric : euclidean 
nn.eps : 0 
verbose : TRUE 
force.recalc : FALSE 
do.plot : FALSE 
graph.name : RNA_nn RNA_snn
```

In [24]:

```
%%R
# convert SeuratObject to AnnData (use current conda environment)
# now integrated into GEfetch2R
Seu2AD = function(seu.obj, method = c("SeuratDisk", "sceasy", "scDIOR"), out.folder = NULL,
                  out.filename = NULL, assay="RNA", slot = "counts", save.scale = FALSE){
  # check parameters
  method <- match.arg(arg = method)
  # check folder
  if(is.null(out.folder)){
    out.folder = getwd()
  }
  if(! dir.exists(out.folder)){
    message(out.folder, " does not exist, create automatically!")
    dir.create(path = out.folder, showWarnings = FALSE)
  }
  # out name
  out.name = deparse(substitute(seu.obj))
  # conversion
  if(method == "SeuratDisk"){
    if(is.null(out.filename)){
      seu.out.name = file.path(out.folder, paste0(out.name, "_SeuratDisk.h5Seurat"))
    }else{
      seu.out.name = file.path(out.folder, out.filename)
    }
    if(save.scale){
      seu.scale = Seurat::GetAssayData(object = seu.obj, slot = "scale.data", assay = assay)
      if(nrow(seu.scale) == 0){
        message("There is no scale.data in seu.obj!")
      }
    }else{
      seu.obj[["RNA"]]@scale.data = matrix(numeric(0),0,0)
    }
    seu.log = tryCatch(
      {
        SeuratDisk::SaveH5Seurat(seu.obj, filename = seu.out.name, overwrite = TRUE)
        SeuratDisk::Convert(seu.out.name, dest = "h5ad", assay = assay, overwrite = TRUE)
      },
      error = function(cond) {
        message("There is an error when using SeuratDisk: ", cond)
      }
    )
    return(seu.log)
  }else if(method == "sceasy"){
    if(is.null(out.filename)){
      sceasy.out.name = file.path(out.folder, paste0(out.name, "_sceasy.h5ad"))
    }else{
      sceasy.out.name = file.path(out.folder, out.filename)
    }
    sceasy.log = tryCatch(
      {
        # reticulate::use_condaenv("/Applications/anaconda3", required = TRUE)
        # or set RETICULATE_PYTHON = "/Applications/anaconda3/bin/python" in Renvion
        sceasy::convertFormat(seu.obj, from="seurat", to="anndata", drop_single_values = FALSE,
                              outFile=sceasy.out.name, main_layer = slot, assay=assay)
      },
      error = function(cond) {
        message("There is an error when using sceasy: ", cond)
      }
    )
    return(sceasy.log)
  }else if(method == "scDIOR"){
    if(is.null(out.filename)){
      scdior.out.name = file.path(out.folder, paste0(out.name, "_scDIOR.h5"))
    }else{
      scdior.out.name = file.path(out.folder, out.filename)
    }
    scdior.log = tryCatch(
      {
        dior::write_h5(data = seu.obj, object.type = "seurat", file = scdior.out.name,
                       assay.name = assay, save.scale = save.scale)
        # adata = diopy.input.read_h5(file = 'pbmc3k.h5') # require diopy to load h5 to AnnData
      },
      error = function(cond) {
        message("There is an error when using scDIOR: ", cond)
      }
    )
    return(scdior.log)
  }
}
```

#### SeuratDisk¶

**Retained information** (`SeuratObject -> AnnData`):

- count matrix: log-normalized data (`data -> raw.X/X`), scaled data (`scale.data -> X`)/raw count matrix (`counts -> raw.X`) (`save.scale = TRUE/FALSE`)
- cells' meta-information (`meta.data -> obs`)
- annotation of features (`meta.features -> var`)
- dimensional reduction results (`reductions -> obsm`)
- relationship of cells, graphs (only contains `RNA_snn` graph) (`graphs -> obsp`)
- alternative assay (`assay -> layers`)

In [25]:

```
%%R
# set save.scale=TRUE, scaled data is stored in X，log-normalized data is in raw.X
# set save.scale=FALSE, log-normalized data is stored in X，raw count matrix is in raw.X
Seu2AD(seu.obj = pbmc3k.final, method = "SeuratDisk", out.folder = "./",
       assay="RNA", save.scale = TRUE)
```

```
R[write to console]: Registered S3 method overwritten by 'SeuratDisk':
  method            from  
  as.sparse.H5Group Seurat

R[write to console]: Creating h5Seurat file for version 3.1.5.9900

R[write to console]: Adding counts for RNA

R[write to console]: Adding data for RNA

R[write to console]: Adding scale.data for RNA

R[write to console]: Adding variable features for RNA

R[write to console]: Adding feature-level metadata for RNA

R[write to console]: Adding counts for rawcounts

R[write to console]: Adding data for rawcounts

R[write to console]: No variable features found for rawcounts

R[write to console]: No feature-level metadata found for rawcounts

R[write to console]: Adding cell embeddings for pca

R[write to console]: Adding loadings for pca

R[write to console]: No projected loadings for pca

R[write to console]: Adding standard deviations for pca

R[write to console]: Adding JackStraw information for pca

R[write to console]: Adding cell embeddings for umap

R[write to console]: No loadings for umap

R[write to console]: No projected loadings for umap

R[write to console]: No standard deviations for umap

R[write to console]: No JackStraw data for umap

R[write to console]: Validating h5Seurat file

R[write to console]: Adding scale.data from RNA as X

R[write to console]: Transfering meta.features to var

R[write to console]: Adding data from RNA as raw

R[write to console]: Transfering meta.features to raw/var

R[write to console]: Transfering meta.data to obs

R[write to console]: Adding dimensional reduction information for pca

R[write to console]: Adding feature loadings for pca

R[write to console]: Adding dimensional reduction information for umap

R[write to console]: Adding RNA_snn as neighbors

R[write to console]: Adding data from rawcounts as a layer
```

```
[1] "/Users/soyabean/Desktop/tmp/scdown/benchmark/pbmc3k.final_SeuratDisk.h5ad"
```

In [26]:

```
seudisk_ann = sc.read("./pbmc3k.final_SeuratDisk.h5ad")
seudisk_ann
```

```
/Applications/anaconda3/lib/python3.7/site-packages/anndata/compat/__init__.py:235: FutureWarning: Moving element from .uns['neighbors']['distances'] to .obsp['distances'].

This is where adjacency matrices should go now.
  FutureWarning,
```

Out[26]:

```
AnnData object with n_obs × n_vars = 2638 × 13714
    obs: 'orig.ident', 'nCount_RNA', 'nFeature_RNA', 'seurat_annotations', 'percent.mt', 'RNA_snn_res.0.5', 'seurat_clusters', 'nCount_rawcounts', 'nFeature_rawcounts'
    var: 'vst.mean', 'vst.variance', 'vst.variance.expected', 'vst.variance.standardized', 'vst.variable', 'rawcounts_features'
    uns: 'neighbors'
    obsm: 'X_pca', 'X_umap'
    varm: 'PCs'
    layers: 'rawcounts'
    obsp: 'distances'
```

In [27]:

```
# scaled count matrix
seudisk_ann.X[0:10,0:4]
# shape
seudisk_ann.X.shape
```

Out[27]:

```
array([[-0.05812316, -0.03357571, -0.04166819, -0.03364562],
       [-0.05812316, -0.03357571, -0.04166819, -0.03364562],
       [-0.05812316, -0.03357571, -0.04166819, -0.03364562],
       [-0.05812316, -0.03357571, -0.04166819, -0.03364562],
       [-0.05812316, -0.03357571, -0.04166819, -0.03364562],
       [-0.05812316, -0.03357571, -0.04166819, -0.03364562],
       [-0.05812316, -0.03357571, -0.04166819, -0.03364562],
       [-0.05812316, -0.03357571, -0.04166819, -0.03364562],
       [-0.05812316, -0.03357571, -0.04166819, -0.03364562],
       [-0.05812316, -0.03357571, -0.04166819, -0.03364562]])
```

Out[27]:

```
(2638, 13714)
```

In [28]:

```
# log-normalized count matrix
seudisk_ann.raw.to_adata().X.toarray()[0:10,0:15]
# shape
seudisk_ann.raw.to_adata().X.shape
```

```
/Applications/anaconda3/lib/python3.7/site-packages/anndata/_core/raw.py:146: FutureWarning: X.dtype being converted to np.float32 from float64. In the next version of anndata (0.9) conversion will not be automatic. Pass dtype explicitly to avoid this warning. Pass `AnnData(X, dtype=X.dtype, ...)` to get the future behavour.
  uns=self._adata.uns.copy(),
```

Out[28]:

```
array([[0.       , 0.       , 0.       , 0.       , 0.       , 0.       ,
        0.       , 0.       , 0.       , 0.       , 0.       , 0.       ,
        0.       , 0.       , 0.       ],
       [0.       , 0.       , 0.       , 0.       , 0.       , 0.       ,
        0.       , 0.       , 0.       , 0.       , 0.       , 0.       ,
        0.       , 0.       , 1.625141 ],
       [0.       , 0.       , 0.       , 0.       , 0.       , 0.       ,
        0.       , 0.       , 0.       , 0.       , 0.       , 1.429744 ,
        0.       , 0.       , 0.       ],
       [0.       , 0.       , 0.       , 0.       , 0.       , 0.       ,
        0.       , 0.       , 0.       , 0.       , 0.       , 3.5583103,
        0.       , 0.       , 0.       ],
       [0.       , 0.       , 0.       , 0.       , 0.       , 0.       ,
        0.       , 0.       , 0.       , 0.       , 0.       , 0.       ,
        0.       , 0.       , 0.       ],
       [0.       , 0.       , 0.       , 0.       , 0.       , 0.       ,
        0.       , 0.       , 0.       , 0.       , 0.       , 1.7269024,
        0.       , 0.       , 0.       ],
       [0.       , 0.       , 0.       , 0.       , 0.       , 0.       ,
        0.       , 0.       , 0.       , 0.       , 0.       , 0.       ,
        0.       , 0.       , 0.       ],
       [0.       , 0.       , 0.       , 0.       , 0.       , 0.       ,
        0.       , 0.       , 0.       , 0.       , 0.       , 0.       ,
        0.       , 0.       , 0.       ],
       [0.       , 0.       , 0.       , 0.       , 0.       , 0.       ,
        0.       , 0.       , 0.       , 0.       , 0.       , 0.       ,
        0.       , 0.       , 0.       ],
       [0.       , 0.       , 0.       , 0.       , 0.       , 0.       ,
        0.       , 0.       , 0.       , 0.       , 0.       , 3.3392706,
        0.       , 0.       , 0.       ]], dtype=float32)
```

Out[28]:

```
(2638, 13714)
```

In [29]:

```
# cells' meta-information
seudisk_ann.obs.head()
```

Out[29]:

|  | orig.ident | nCount\_RNA | nFeature\_RNA | seurat\_annotations | percent.mt | RNA\_snn\_res.0.5 | seurat\_clusters | nCount\_rawcounts | nFeature\_rawcounts |
| --- | --- | --- | --- | --- | --- | --- | --- | --- | --- |
| AAACATACAACCAC | 0 | 2419.0 | 779 | 1 | 3.017776 | 1 | 1 | 2419.0 | 779 |
| AAACATTGAGCTAC | 0 | 4903.0 | 1352 | 3 | 3.793596 | 3 | 3 | 4903.0 | 1352 |
| AAACATTGATCAGC | 0 | 3147.0 | 1129 | 1 | 0.889736 | 1 | 1 | 3147.0 | 1129 |
| AAACCGTGCTTCCG | 0 | 2639.0 | 960 | 2 | 1.743085 | 2 | 2 | 2639.0 | 960 |
| AAACCGTGTATGCG | 0 | 980.0 | 521 | 6 | 1.224490 | 6 | 6 | 980.0 | 521 |

In [30]:

```
# annotation of features
seudisk_ann.var.head()
# shape
seudisk_ann.var.shape
```

Out[30]:

|  | vst.mean | vst.variance | vst.variance.expected | vst.variance.standardized | vst.variable | rawcounts\_features |
| --- | --- | --- | --- | --- | --- | --- |
| AL627309.1 | 0.003412 | 0.003401 | 0.003645 | 0.933044 | 0 | AL627309.1 |
| AP006222.2 | 0.001137 | 0.001136 | 0.001145 | 0.992494 | 0 | AP006222.2 |
| RP11-206L10.2 | 0.001895 | 0.001893 | 0.001966 | 0.962729 | 0 | RP11-206L10.2 |
| RP11-206L10.9 | 0.001137 | 0.001136 | 0.001145 | 0.992494 | 0 | RP11-206L10.9 |
| LINC00115 | 0.006823 | 0.006779 | 0.007481 | 0.906213 | 0 | LINC00115 |

Out[30]:

```
(13714, 6)
```

In [31]:

```
# unstructured annotation, converted from pbmc3k.final@commands$FindNeighbors.RNA.pca
seudisk_ann.uns['neighbors']['params']
```

Out[31]:

```
{'method': array(['snn'], dtype=object), 'n_neighbors': array([20.])}
```

In [32]:

```
# dimensional reduction results of pca
seudisk_ann.obsm['X_pca'][0:5, 0:5]
# dimensional reduction results of umap
seudisk_ann.obsm['X_umap'][0:5, 0:2]
```

Out[32]:

```
array([[-4.72968551, -0.51842651, -0.76232201, -2.31567898, -0.07160006],
       [-0.51740293,  4.59189566,  5.90919209,  6.91188558, -1.96243034],
       [-3.18910634, -3.46951536, -0.83137104, -2.00199849, -5.10442765],
       [12.79330206,  0.10071659,  0.63102207, -0.36873382,  0.21838204],
       [-3.12880778, -6.34814123,  1.25077756,  3.01910262,  7.84739502]])
```

Out[32]:

```
array([[-4.23279204, -4.1521394 ],
       [-4.89288606, 10.98568513],
       [-5.50863876, -7.2110884 ],
       [11.33223281,  3.16172697],
       [-7.45070281,  1.09202202]])
```

In [33]:

```
# feature loadings, wrong values
seudisk_ann.varm['PCs'][0:5, 0:5]
```

Out[33]:

```
array([[nan, nan, nan, nan, nan],
       [nan, nan, nan, nan, nan],
       [nan, nan, nan, nan, nan],
       [nan, nan, nan, nan, nan],
       [nan, nan, nan, nan, nan]])
```

In [34]:

```
# layers (raw count matrix)
seudisk_ann.layers['rawcounts'].toarray()
# shape
seudisk_ann.layers['rawcounts'].shape
```

Out[34]:

```
array([[0., 0., 0., ..., 0., 0., 0.],
       [0., 0., 0., ..., 0., 0., 0.],
       [0., 0., 0., ..., 0., 0., 0.],
       ...,
       [0., 0., 0., ..., 0., 0., 0.],
       [0., 0., 0., ..., 1., 0., 0.],
       [0., 0., 0., ..., 0., 0., 0.]])
```

Out[34]:

```
(2638, 13714)
```

In [35]:

```
# relationship of cells, graphs
seudisk_ann.obsp
# only contains RNA_snn graph
seudisk_ann.obsp['distances'].toarray()[0:10,0:10]
```

Out[35]:

```
PairwiseArrays with keys: distances
```

Out[35]:

```
array([[1.        , 0.        , 0.        , 0.        , 0.        ,
        0.        , 0.11111111, 0.        , 0.        , 0.        ],
       [0.        , 1.        , 0.        , 0.        , 0.        ,
        0.        , 0.        , 0.        , 0.        , 0.        ],
       [0.        , 0.        , 1.        , 0.        , 0.        ,
        0.        , 0.        , 0.        , 0.        , 0.        ],
       [0.        , 0.        , 0.        , 1.        , 0.        ,
        0.        , 0.        , 0.        , 0.        , 0.        ],
       [0.        , 0.        , 0.        , 0.        , 1.        ,
        0.        , 0.        , 0.        , 0.        , 0.        ],
       [0.        , 0.        , 0.        , 0.        , 0.        ,
        1.        , 0.        , 0.        , 0.        , 0.        ],
       [0.11111111, 0.        , 0.        , 0.        , 0.        ,
        0.        , 1.        , 0.08108108, 0.        , 0.        ],
       [0.        , 0.        , 0.        , 0.        , 0.        ,
        0.        , 0.08108108, 1.        , 0.        , 0.        ],
       [0.        , 0.        , 0.        , 0.        , 0.        ,
        0.        , 0.        , 0.        , 1.        , 0.        ],
       [0.        , 0.        , 0.        , 0.        , 0.        ,
        0.        , 0.        , 0.        , 0.        , 1.        ]])
```

#### sceasy¶

**Retained information** (`SeuratObject -> AnnData`):

- count matrix: log-normalized data/scaled data/raw count matrix (`data/scale.data/counts -> X`) (`slot` parameter)
- cells' meta-information (`meta.data -> obs`)
- annotation of features (`meta.features -> var`)
- dimensional reduction results (`reductions -> obsm`)

In [36]:

```
%%R
# slot = "counts": use raw count matrix 
Seu2AD(seu.obj = pbmc3k.final, method = "sceasy", out.folder = "./",
       assay="RNA", slot = "counts")
```

```
/Applications/anaconda3/lib/python3.7/site-packages/rpy2/rinterface.py:807: FutureWarning: X.dtype being converted to np.float32 from float64. In the next version of anndata (0.9) conversion will not be automatic. Pass dtype explicitly to avoid this warning. Pass `AnnData(X, dtype=X.dtype, ...)` to get the future behavour.
  error_occured)
```

```
AnnData object with n_obs × n_vars = 2638 × 13714
    obs: 'orig.ident', 'nCount_RNA', 'nFeature_RNA', 'seurat_annotations', 'percent.mt', 'RNA_snn_res.0.5', 'seurat_clusters', 'nCount_rawcounts', 'nFeature_rawcounts'
    var: 'vst.mean', 'vst.variance', 'vst.variance.expected', 'vst.variance.standardized', 'vst.variable'
    obsm: 'X_pca', 'X_umap'
```

In [37]:

```
seusceasy_ann = sc.read("./pbmc3k.final_sceasy.h5ad")
seusceasy_ann
```

Out[37]:

```
AnnData object with n_obs × n_vars = 2638 × 13714
    obs: 'orig.ident', 'nCount_RNA', 'nFeature_RNA', 'seurat_annotations', 'percent.mt', 'RNA_snn_res.0.5', 'seurat_clusters', 'nCount_rawcounts', 'nFeature_rawcounts'
    var: 'vst.mean', 'vst.variance', 'vst.variance.expected', 'vst.variance.standardized', 'vst.variable'
    obsm: 'X_pca', 'X_umap'
```

In [38]:

```
# raw count matrix
seusceasy_ann.X.toarray()
# shape
seusceasy_ann.X.shape
```

Out[38]:

```
array([[0., 0., 0., ..., 0., 0., 0.],
       [0., 0., 0., ..., 0., 0., 0.],
       [0., 0., 0., ..., 0., 0., 0.],
       ...,
       [0., 0., 0., ..., 0., 0., 0.],
       [0., 0., 0., ..., 1., 0., 0.],
       [0., 0., 0., ..., 0., 0., 0.]], dtype=float32)
```

Out[38]:

```
(2638, 13714)
```

In [39]:

```
# cells' meta-information
seusceasy_ann.obs.head()
```

Out[39]:

|  | orig.ident | nCount\_RNA | nFeature\_RNA | seurat\_annotations | percent.mt | RNA\_snn\_res.0.5 | seurat\_clusters | nCount\_rawcounts | nFeature\_rawcounts |
| --- | --- | --- | --- | --- | --- | --- | --- | --- | --- |
| AAACATACAACCAC | pbmc3k | 2419.0 | 779 | Memory CD4 T | 3.017776 | 1 | 1 | 2419.0 | 779 |
| AAACATTGAGCTAC | pbmc3k | 4903.0 | 1352 | B | 3.793596 | 3 | 3 | 4903.0 | 1352 |
| AAACATTGATCAGC | pbmc3k | 3147.0 | 1129 | Memory CD4 T | 0.889736 | 1 | 1 | 3147.0 | 1129 |
| AAACCGTGCTTCCG | pbmc3k | 2639.0 | 960 | CD14+ Mono | 1.743085 | 2 | 2 | 2639.0 | 960 |
| AAACCGTGTATGCG | pbmc3k | 980.0 | 521 | NK | 1.224490 | 6 | 6 | 980.0 | 521 |

In [40]:

```
# annotation of features
seusceasy_ann.var.head()
# shape
seusceasy_ann.var.shape
```

Out[40]:

|  | vst.mean | vst.variance | vst.variance.expected | vst.variance.standardized | vst.variable |
| --- | --- | --- | --- | --- | --- |
| AL627309.1 | 0.003412 | 0.003401 | 0.003645 | 0.933044 | False |
| AP006222.2 | 0.001137 | 0.001136 | 0.001145 | 0.992494 | False |
| RP11-206L10.2 | 0.001895 | 0.001893 | 0.001966 | 0.962729 | False |
| RP11-206L10.9 | 0.001137 | 0.001136 | 0.001145 | 0.992494 | False |
| LINC00115 | 0.006823 | 0.006779 | 0.007481 | 0.906213 | False |

Out[40]:

```
(13714, 5)
```

In [41]:

```
# dimensional reduction results of pca
seusceasy_ann.obsm['X_pca'][0:5, 0:5]
# dimensional reduction results of umap
seusceasy_ann.obsm['X_umap'][0:5, 0:2]
```

Out[41]:

```
array([[-4.72968551, -0.51842651, -0.76232201, -2.31567898, -0.07160006],
       [-0.51740293,  4.59189566,  5.90919209,  6.91188558, -1.96243034],
       [-3.18910634, -3.46951536, -0.83137104, -2.00199849, -5.10442765],
       [12.79330206,  0.10071659,  0.63102207, -0.36873382,  0.21838204],
       [-3.12880778, -6.34814123,  1.25077756,  3.01910262,  7.84739502]])
```

Out[41]:

```
array([[-4.23279204, -4.1521394 ],
       [-4.89288606, 10.98568513],
       [-5.50863876, -7.2110884 ],
       [11.33223281,  3.16172697],
       [-7.45070281,  1.09202202]])
```

#### scDIOR¶

**Retained information** (`SeuratObject -> AnnData`):

- count matrix:
  - `save.scale = TRUE`: log-normalized data (`data -> raw.X`), scaled data (`scale.data -> X`), raw count matrix (`counts -> layers`)
  - `save.scale = FALSE`: log-normalized data (`data -> X`), raw count matrix (`counts -> layers`)
- cells' meta-information (`meta.data -> obs`)
- annotation of features (`meta.features -> var`)
- dimensional reduction results (`reductions -> obsm`)
- relationship of cells, graphs (contains `RNA_snn` and `RNA_nn` graphs) (`graphs -> obsp`)
- alternative assay (`assay -> layers`)

scDIOR requires diopy to read `.h5` file.

In [42]:

```
%%R
# set save.scale=TRUE, scaled data is stored in X，log-normalized data is in raw.X, raw count matrix is in layers
# set save.scale=FALSE, log-normalized data is stored in X，raw count matrix is in layers
Seu2AD(seu.obj = pbmc3k.final, method = "scDIOR", out.folder = "./",
       assay="RNA", save.scale = TRUE)
```

```
NULL
```

In [43]:

```
# scDIOR require diopy to read h5
import diopy
```

In [44]:

```
seuscdior_ann = diopy.input.read_h5(file = "./pbmc3k.final_scDIOR.h5")
seuscdior_ann
```

Out[44]:

```
AnnData object with n_obs × n_vars = 2638 × 13714
    obs: 'orig.ident', 'nCount_RNA', 'nFeature_RNA', 'seurat_annotations', 'percent.mt', 'RNA_snn_res.0.5', 'seurat_clusters', 'nCount_rawcounts', 'nFeature_rawcounts'
    var: 'vst.mean', 'vst.variance', 'vst.variance.expected', 'vst.variance.standardized', 'vst.variable'
    obsm: 'X_pca', 'X_umap'
    layers: 'counts', 'rawcounts'
    obsp: 'distances', 'connectivities'
```

In [45]:

```
# scaled count matrix
seuscdior_ann.X[0:10,0:4]
# shape
seuscdior_ann.X.shape
```

Out[45]:

```
array([[-0.05812316, -0.03357571, -0.04166819, -0.03364562],
       [-0.05812316, -0.03357571, -0.04166819, -0.03364562],
       [-0.05812316, -0.03357571, -0.04166819, -0.03364562],
       [-0.05812316, -0.03357571, -0.04166819, -0.03364562],
       [-0.05812316, -0.03357571, -0.04166819, -0.03364562],
       [-0.05812316, -0.03357571, -0.04166819, -0.03364562],
       [-0.05812316, -0.03357571, -0.04166819, -0.03364562],
       [-0.05812316, -0.03357571, -0.04166819, -0.03364562],
       [-0.05812316, -0.03357571, -0.04166819, -0.03364562],
       [-0.05812316, -0.03357571, -0.04166819, -0.03364562]],
      dtype=float32)
```

Out[45]:

```
(2638, 13714)
```

In [46]:

```
# log-normalized count matrix
seuscdior_ann.raw.to_adata().X.toarray()[0:10,0:15]
# shape
seuscdior_ann.raw.to_adata().X.shape
```

Out[46]:

```
array([[0.       , 0.       , 0.       , 0.       , 0.       , 0.       ,
        0.       , 0.       , 0.       , 0.       , 0.       , 0.       ,
        0.       , 0.       , 0.       ],
       [0.       , 0.       , 0.       , 0.       , 0.       , 0.       ,
        0.       , 0.       , 0.       , 0.       , 0.       , 0.       ,
        0.       , 0.       , 1.625141 ],
       [0.       , 0.       , 0.       , 0.       , 0.       , 0.       ,
        0.       , 0.       , 0.       , 0.       , 0.       , 1.429744 ,
        0.       , 0.       , 0.       ],
       [0.       , 0.       , 0.       , 0.       , 0.       , 0.       ,
        0.       , 0.       , 0.       , 0.       , 0.       , 3.5583103,
        0.       , 0.       , 0.       ],
       [0.       , 0.       , 0.       , 0.       , 0.       , 0.       ,
        0.       , 0.       , 0.       , 0.       , 0.       , 0.       ,
        0.       , 0.       , 0.       ],
       [0.       , 0.       , 0.       , 0.       , 0.       , 0.       ,
        0.       , 0.       , 0.       , 0.       , 0.       , 1.7269024,
        0.       , 0.       , 0.       ],
       [0.       , 0.       , 0.       , 0.       , 0.       , 0.       ,
        0.       , 0.       , 0.       , 0.       , 0.       , 0.       ,
        0.       , 0.       , 0.       ],
       [0.       , 0.       , 0.       , 0.       , 0.       , 0.       ,
        0.       , 0.       , 0.       , 0.       , 0.       , 0.       ,
        0.       , 0.       , 0.       ],
       [0.       , 0.       , 0.       , 0.       , 0.       , 0.       ,
        0.       , 0.       , 0.       , 0.       , 0.       , 0.       ,
        0.       , 0.       , 0.       ],
       [0.       , 0.       , 0.       , 0.       , 0.       , 0.       ,
        0.       , 0.       , 0.       , 0.       , 0.       , 3.3392706,
        0.       , 0.       , 0.       ]], dtype=float32)
```

Out[46]:

```
(2638, 13714)
```

In [47]:

```
# raw count matrix
seuscdior_ann.layers['counts'].toarray()
```

Out[47]:

```
array([[0., 0., 0., ..., 0., 0., 0.],
       [0., 0., 0., ..., 0., 0., 0.],
       [0., 0., 0., ..., 0., 0., 0.],
       ...,
       [0., 0., 0., ..., 0., 0., 0.],
       [0., 0., 0., ..., 1., 0., 0.],
       [0., 0., 0., ..., 0., 0., 0.]], dtype=float32)
```

In [48]:

```
# cells' meta-information
seuscdior_ann.obs.head()
```

Out[48]:

|  | orig.ident | nCount\_RNA | nFeature\_RNA | seurat\_annotations | percent.mt | RNA\_snn\_res.0.5 | seurat\_clusters | nCount\_rawcounts | nFeature\_rawcounts |
| --- | --- | --- | --- | --- | --- | --- | --- | --- | --- |
| index |  |  |  |  |  |  |  |  |  |
| AAACATACAACCAC | pbmc3k | 2419.0 | 779 | Memory CD4 T | 3.017776 | 1 | 1 | 2419.0 | 779 |
| AAACATTGAGCTAC | pbmc3k | 4903.0 | 1352 | B | 3.793596 | 3 | 3 | 4903.0 | 1352 |
| AAACATTGATCAGC | pbmc3k | 3147.0 | 1129 | Memory CD4 T | 0.889736 | 1 | 1 | 3147.0 | 1129 |
| AAACCGTGCTTCCG | pbmc3k | 2639.0 | 960 | CD14+ Mono | 1.743085 | 2 | 2 | 2639.0 | 960 |
| AAACCGTGTATGCG | pbmc3k | 980.0 | 521 | NK | 1.224490 | 6 | 6 | 980.0 | 521 |

In [49]:

```
# annotation of features
seuscdior_ann.var.head()
# shape
seuscdior_ann.var.shape
```

Out[49]:

|  | vst.mean | vst.variance | vst.variance.expected | vst.variance.standardized | vst.variable |
| --- | --- | --- | --- | --- | --- |
| index |  |  |  |  |  |
| AL627309.1 | 0.003412 | 0.003401 | 0.003645 | 0.933044 | False |
| AP006222.2 | 0.001137 | 0.001136 | 0.001145 | 0.992494 | False |
| RP11-206L10.2 | 0.001895 | 0.001893 | 0.001966 | 0.962729 | False |
| RP11-206L10.9 | 0.001137 | 0.001136 | 0.001145 | 0.992494 | False |
| LINC00115 | 0.006823 | 0.006779 | 0.007481 | 0.906213 | False |

Out[49]:

```
(13714, 5)
```

In [50]:

```
# dimensional reduction results of pca
seuscdior_ann.obsm['X_pca'][0:5, 0:5]
# dimensional reduction results of umap
seuscdior_ann.obsm['X_umap'][0:5, 0:2]
```

Out[50]:

```
array([[-4.72968551, -0.51842651, -0.76232201, -2.31567898, -0.07160006],
       [-0.51740293,  4.59189566,  5.90919209,  6.91188558, -1.96243034],
       [-3.18910634, -3.46951536, -0.83137104, -2.00199849, -5.10442765],
       [12.79330206,  0.10071659,  0.63102207, -0.36873382,  0.21838204],
       [-3.12880778, -6.34814123,  1.25077756,  3.01910262,  7.84739502]])
```

Out[50]:

```
array([[-4.23279204, -4.1521394 ],
       [-4.89288606, 10.98568513],
       [-5.50863876, -7.2110884 ],
       [11.33223281,  3.16172697],
       [-7.45070281,  1.09202202]])
```

In [51]:

```
# layers (raw count matrix)
seuscdior_ann.layers['rawcounts'].toarray()
# shape
seuscdior_ann.layers['rawcounts'].shape
```

Out[51]:

```
array([[0., 0., 0., ..., 0., 0., 0.],
       [0., 0., 0., ..., 0., 0., 0.],
       [0., 0., 0., ..., 0., 0., 0.],
       ...,
       [0., 0., 0., ..., 0., 0., 0.],
       [0., 0., 0., ..., 1., 0., 0.],
       [0., 0., 0., ..., 0., 0., 0.]], dtype=float32)
```

Out[51]:

```
(2638, 13714)
```

In [52]:

```
# relationship of cells, graphs
seuscdior_ann.obsp
# RNA_nn graph
seuscdior_ann.obsp['distances'].toarray()[0:10,0:10]
# RNA_snn graph
seuscdior_ann.obsp['connectivities'].toarray()[0:10,0:10]
```

Out[52]:

```
PairwiseArrays with keys: distances, connectivities
```

Out[52]:

```
array([[1., 0., 0., 0., 0., 0., 1., 0., 0., 0.],
       [0., 1., 0., 0., 0., 0., 0., 0., 0., 0.],
       [0., 0., 1., 0., 0., 0., 0., 0., 0., 0.],
       [0., 0., 0., 1., 0., 0., 0., 0., 0., 0.],
       [0., 0., 0., 0., 1., 0., 0., 0., 0., 0.],
       [0., 0., 0., 0., 0., 1., 0., 0., 0., 0.],
       [0., 0., 0., 0., 0., 0., 1., 0., 0., 0.],
       [0., 0., 0., 0., 0., 0., 0., 1., 0., 0.],
       [0., 0., 0., 0., 0., 0., 0., 0., 1., 0.],
       [0., 0., 0., 0., 0., 0., 0., 0., 0., 1.]], dtype=float32)
```

Out[52]:

```
array([[1.        , 0.        , 0.        , 0.        , 0.        ,
        0.        , 0.11111111, 0.        , 0.        , 0.        ],
       [0.        , 1.        , 0.        , 0.        , 0.        ,
        0.        , 0.        , 0.        , 0.        , 0.        ],
       [0.        , 0.        , 1.        , 0.        , 0.        ,
        0.        , 0.        , 0.        , 0.        , 0.        ],
       [0.        , 0.        , 0.        , 1.        , 0.        ,
        0.        , 0.        , 0.        , 0.        , 0.        ],
       [0.        , 0.        , 0.        , 0.        , 1.        ,
        0.        , 0.        , 0.        , 0.        , 0.        ],
       [0.        , 0.        , 0.        , 0.        , 0.        ,
        1.        , 0.        , 0.        , 0.        , 0.        ],
       [0.11111111, 0.        , 0.        , 0.        , 0.        ,
        0.        , 1.        , 0.08108108, 0.        , 0.        ],
       [0.        , 0.        , 0.        , 0.        , 0.        ,
        0.        , 0.08108108, 1.        , 0.        , 0.        ],
       [0.        , 0.        , 0.        , 0.        , 0.        ,
        0.        , 0.        , 0.        , 1.        , 0.        ],
       [0.        , 0.        , 0.        , 0.        , 0.        ,
        0.        , 0.        , 0.        , 0.        , 1.        ]],
      dtype=float32)
```

### AnnData to SeuratObject¶

#### AnnData¶

**data source and preprocessing**: Preprocessing and clustering 3k PBMCs (legacy workflow)

In [71]:

```
pbmc3k_ann = sc.read("./write/pbmc3k.h5ad")
pbmc3k_ann
```

Out[71]:

```
AnnData object with n_obs × n_vars = 2638 × 1838
    obs: 'n_genes', 'n_genes_by_counts', 'total_counts', 'total_counts_mt', 'pct_counts_mt', 'leiden'
    var: 'gene_ids', 'n_cells', 'mt', 'n_cells_by_counts', 'mean_counts', 'pct_dropout_by_counts', 'total_counts', 'highly_variable', 'means', 'dispersions', 'dispersions_norm', 'mean', 'std'
    uns: 'hvg', 'leiden', 'log1p', 'neighbors', 'pca', 'rank_genes_groups', 'umap'
    obsm: 'X_pca', 'X_umap'
    varm: 'PCs'
    layers: 'logcounts', 'rawcounts'
    obsp: 'connectivities', 'distances'
```

##### Count matrix (`X` and `layers`)¶

In [72]:

```
# raw count matrix
pbmc3k_ann.raw.X.toarray()
# shape
pbmc3k_ann.raw.X.shape
```

Out[72]:

```
array([[0., 0., 0., ..., 0., 0., 0.],
       [0., 0., 0., ..., 0., 0., 0.],
       [0., 0., 0., ..., 0., 0., 0.],
       ...,
       [0., 0., 0., ..., 0., 0., 0.],
       [0., 0., 0., ..., 1., 0., 0.],
       [0., 0., 0., ..., 0., 0., 0.]], dtype=float32)
```

Out[72]:

```
(2638, 13714)
```

In [73]:

```
# raw count matrix in layers
pbmc3k_ann.layers['rawcounts'].toarray()
# shape
pbmc3k_ann.layers['rawcounts'].shape
```

Out[73]:

```
array([[0., 0., 0., ..., 0., 0., 0.],
       [0., 0., 0., ..., 0., 0., 0.],
       [0., 0., 0., ..., 0., 0., 1.],
       ...,
       [0., 0., 0., ..., 0., 0., 1.],
       [0., 0., 0., ..., 0., 0., 0.],
       [0., 0., 0., ..., 0., 0., 0.]], dtype=float32)
```

Out[73]:

```
(2638, 1838)
```

In [74]:

```
# log-normalized count matrix
pbmc3k_ann.layers['logcounts'].toarray()
# shape
pbmc3k_ann.layers['logcounts'].shape
```

Out[74]:

```
array([[0.       , 0.       , 0.       , ..., 0.       , 0.       ,
        0.       ],
       [0.       , 0.       , 0.       , ..., 0.       , 0.       ,
        0.       ],
       [0.       , 0.       , 0.       , ..., 0.       , 0.       ,
        1.429744 ],
       ...,
       [0.       , 0.       , 0.       , ..., 0.       , 0.       ,
        1.9370484],
       [0.       , 0.       , 0.       , ..., 0.       , 0.       ,
        0.       ],
       [0.       , 0.       , 0.       , ..., 0.       , 0.       ,
        0.       ]], dtype=float32)
```

Out[74]:

```
(2638, 1838)
```

In [75]:

```
# scaled count matrix
pbmc3k_ann.X
# shape
pbmc3k_ann.X.shape
```

Out[75]:

```
array([[-0.17146961, -0.2808123 , -0.04667677, ..., -0.09826882,
        -0.20909512, -0.5312033 ],
       [-0.21458235, -0.37265328, -0.05480441, ..., -0.266844  ,
        -0.31314582, -0.5966543 ],
       [-0.3768877 , -0.29508454, -0.05752748, ..., -0.15865591,
        -0.17087644,  1.3789997 ],
       ...,
       [-0.20708963, -0.2504642 , -0.04639699, ..., -0.05114426,
        -0.16106427,  2.041497  ],
       [-0.1903285 , -0.2263338 , -0.04399936, ..., -0.00591774,
        -0.13521305, -0.48211104],
       [-0.33378935, -0.25358772, -0.05271561, ..., -0.07842438,
        -0.13032718, -0.47133783]], dtype=float32)
```

Out[75]:

```
(2638, 1838)
```

##### `obs` - cells' meta-information¶

In [76]:

```
pbmc3k_ann.obs.head()
# shape
pbmc3k_ann.var.shape
```

Out[76]:

|  | n\_genes | n\_genes\_by\_counts | total\_counts | total\_counts\_mt | pct\_counts\_mt | leiden |
| --- | --- | --- | --- | --- | --- | --- |
| AAACATACAACCAC-1 | 781 | 779 | 2419.0 | 73.0 | 3.017776 | 0 |
| AAACATTGAGCTAC-1 | 1352 | 1352 | 4903.0 | 186.0 | 3.793596 | 2 |
| AAACATTGATCAGC-1 | 1131 | 1129 | 3147.0 | 28.0 | 0.889736 | 0 |
| AAACCGTGCTTCCG-1 | 960 | 960 | 2639.0 | 46.0 | 1.743085 | 4 |
| AAACCGTGTATGCG-1 | 522 | 521 | 980.0 | 12.0 | 1.224490 | 5 |

Out[76]:

```
(1838, 13)
```

##### `var` - annotation of features¶

In [77]:

```
pbmc3k_ann.var.head()
# shape
pbmc3k_ann.var.shape
```

Out[77]:

|  | gene\_ids | n\_cells | mt | n\_cells\_by\_counts | mean\_counts | pct\_dropout\_by\_counts | total\_counts | highly\_variable | means | dispersions | dispersions\_norm | mean | std |
| --- | --- | --- | --- | --- | --- | --- | --- | --- | --- | --- | --- | --- | --- |
| TNFRSF4 | ENSG00000186827 | 155 | False | 155 | 0.077407 | 94.259259 | 209.0 | True | 0.277410 | 2.086050 | 0.665406 | -3.672069e-10 | 0.424481 |
| CPSF3L | ENSG00000127054 | 202 | False | 202 | 0.094815 | 92.518519 | 256.0 | True | 0.385194 | 4.506987 | 2.955005 | -2.372437e-10 | 0.460416 |
| ATAD3C | ENSG00000215915 | 9 | False | 9 | 0.009259 | 99.666667 | 25.0 | True | 0.038252 | 3.953486 | 4.352607 | 8.472988e-12 | 0.119465 |
| C1orf86 | ENSG00000162585 | 501 | False | 501 | 0.227778 | 81.444444 | 615.0 | True | 0.678283 | 2.713522 | 0.543183 | 3.389195e-10 | 0.685145 |
| RER1 | ENSG00000157916 | 608 | False | 608 | 0.298148 | 77.481481 | 805.0 | True | 0.814813 | 3.447533 | 1.582528 | 7.696297e-11 | 0.736050 |

Out[77]:

```
(1838, 13)
```

##### `uns` - unstructured annotation¶

In [78]:

```
pbmc3k_ann.uns.keys()
```

Out[78]:

```
dict_keys(['hvg', 'leiden', 'log1p', 'neighbors', 'pca', 'rank_genes_groups', 'umap'])
```

In [79]:

```
# pca variance
pbmc3k_ann.uns['pca']['variance']
```

Out[79]:

```
array([32.110455 , 18.718655 , 15.607329 , 13.235289 ,  4.802269 ,
        3.9859324,  3.5262327,  3.2334454,  3.1212087,  3.075261 ,
        2.9980748,  2.959521 ,  2.9517848,  2.9442477,  2.913872 ,
        2.8990302,  2.880682 ,  2.864685 ,  2.8430636,  2.8357508,
        2.8314214,  2.8182364,  2.8035524,  2.7999873,  2.788954 ,
        2.778101 ,  2.7705767,  2.7602205,  2.7538602,  2.7459552,
        2.7371864,  2.7341268,  2.722202 ,  2.7123108,  2.7024777,
        2.7000473,  2.6838503,  2.6790507,  2.6769078,  2.6739945,
        2.6648538,  2.6573114,  2.6511767,  2.6417756,  2.6329703,
        2.6295197,  2.6245294,  2.618376 ,  2.6180034,  2.6018658],
      dtype=float32)
```

In [80]:

```
# pca parameters
pbmc3k_ann.uns['pca']['params']
```

Out[80]:

```
{'use_highly_variable': True, 'zero_center': True}
```

##### `obsm` - dimensional reduction results¶

In [81]:

```
pbmc3k_ann.obsm
```

Out[81]:

```
AxisArrays with keys: X_pca, X_umap
```

In [82]:

```
# dimensional reduction results of pca
pbmc3k_ann.obsm['X_pca']
```

Out[82]:

```
array([[-5.556221  , -0.25772715,  0.18679433, ..., -0.34272835,
         1.4820554 ,  1.8977244 ],
       [-7.209527  , -7.4820013 , -0.16271746, ..., -1.9744129 ,
        -1.5622702 , -1.49611   ],
       [-2.6944373 ,  1.5836617 ,  0.6631235 , ...,  0.544482  ,
        -0.5436244 , -4.3394427 ],
       ...,
       [-0.7853934 , -6.718591  , -1.5988475 , ..., -0.5608387 ,
        -0.10692333,  0.5838822 ],
       [ 0.28127232, -5.9218583 , -1.1628891 , ..., -1.3899633 ,
         3.5770402 ,  1.2988257 ],
       [-0.09076758, -0.6635025 , -0.13485482, ...,  0.37157103,
         0.75083363, -0.6659949 ]], dtype=float32)
```

In [83]:

```
# dimensional reduction results of umap
pbmc3k_ann.obsm['X_umap']
```

Out[83]:

```
array([[ 7.906657 ,  3.556091 ],
       [ 9.248348 , 12.544332 ],
       [ 7.629986 ,  3.8347855],
       ...,
       [ 7.2876377, 13.075106 ],
       [ 8.105372 , 14.307881 ],
       [ 8.511535 ,  3.3921196]], dtype=float32)
```

##### `varm` - feature loadings¶

In [84]:

```
# feature loadings
pbmc3k_ann.varm['PCs']
```

Out[84]:

```
array([[-2.60148179e-02,  3.25416843e-03,  1.89788977e-03, ...,
        -5.18770702e-03,  1.44968908e-02, -6.67473301e-04],
       [-8.27822462e-03,  9.08316299e-03, -7.81411130e-04, ...,
         3.08727100e-02, -8.86981003e-03, -2.88053416e-03],
       [-3.31518659e-03,  3.20968428e-03,  2.79858650e-04, ...,
         1.01477914e-02, -5.30328136e-04,  1.50829612e-03],
       ...,
       [ 8.34176037e-03, -1.24651939e-03, -4.12195362e-03, ...,
        -1.01806019e-02,  9.22558550e-03,  2.79657058e-02],
       [-1.64065659e-02,  4.41013835e-02, -2.13347375e-05, ...,
         9.99553967e-03, -4.50964272e-03, -1.36533342e-02],
       [-1.51882619e-02,  4.00086790e-02,  5.41223399e-03, ...,
        -3.72782419e-03,  2.11074371e-02,  3.59644145e-02]])
```

##### `obsp` - relationship of cells, graphs¶

In [85]:

```
# relationship of cells, graphs
pbmc3k_ann.obsp
```

Out[85]:

```
PairwiseArrays with keys: connectivities, distances
```

In [86]:

```
pbmc3k_ann.obsp['distances'].toarray()
```

Out[86]:

```
array([[0., 0., 0., ..., 0., 0., 0.],
       [0., 0., 0., ..., 0., 0., 0.],
       [0., 0., 0., ..., 0., 0., 0.],
       ...,
       [0., 0., 0., ..., 0., 0., 0.],
       [0., 0., 0., ..., 0., 0., 0.],
       [0., 0., 0., ..., 0., 0., 0.]])
```

In [87]:

```
pbmc3k_ann.obsp['connectivities'].toarray()
```

Out[87]:

```
array([[0., 0., 0., ..., 0., 0., 0.],
       [0., 0., 0., ..., 0., 0., 0.],
       [0., 0., 0., ..., 0., 0., 0.],
       ...,
       [0., 0., 0., ..., 0., 0., 0.],
       [0., 0., 0., ..., 0., 0., 0.],
       [0., 0., 0., ..., 0., 0., 0.]], dtype=float32)
```

In [88]:

```
%%R
# convert AnnData to SeuratObject (use current conda environment)
# now integrated into GEfetch2R
AD2Seu = function(anndata.file, method = c("SeuratDisk", "sceasy",	"scDIOR", "schard", "SeuratDisk+scDIOR"), assay = "RNA",
                  load.assays = "RNA", slot = "counts", use.raw = TRUE){
  # check parameters
  method <- match.arg(arg = method)

  # check file
  if(!file.exists(anndata.file)){
    stop(anndata.file, " does not exist, please check!")
  }
  # conversion
  if(grepl(pattern = "SeuratDisk", x = method)){
    # SeuratDisk
    seu = tryCatch(
      {
        SeuratDisk::Convert(anndata.file, dest = "h5seurat", overwrite = TRUE, assay = assay)
        h5seurat.file = gsub(pattern = "h5ad$", replacement = "h5seurat", x = anndata.file)
        # https://github.com/mojaveazure/seurat-disk/issues/109
        f <- hdf5r::H5File$new(h5seurat.file, "r+")
        groups <- f$ls(recursive = TRUE)
        for (name in groups$name[grepl("categories", groups$name)]) {
          names <- strsplit(name, "/")[[1]]
          names <- c(names[1:length(names) - 1], "levels")
          new_name <- paste(names, collapse = "/")
          f[[new_name]] <- f[[name]]
        }
        for (name in groups$name[grepl("codes", groups$name)]) {
          names <- strsplit(name, "/")[[1]]
          names <- c(names[1:length(names) - 1], "values")
          new_name <- paste(names, collapse = "/")
          f[[new_name]] <- f[[name]]
          grp <- f[[new_name]]
          grp$write(args = list(1:grp$dims), value = grp$read() + 1)
        }
        f$close_all()
        SeuratDisk::LoadH5Seurat(h5seurat.file, assays = load.assays)
      },
      error = function(cond) {
        message("There is an error when using SeuratDisk: ", cond)
      }
    )
    if(grepl(pattern = "scDIOR", x = method)){
      # scDIOR
      seu.scdior = tryCatch(
        {
          dior::read_h5ad(file = anndata.file, assay_name=assay, target.object = "seurat")
        },
        error = function(cond) {
          message("There is an error when using scDIOR: ", cond)
        }
      )
      # add additional assays
      all.assays = Seurat::Assays(seu.scdior)
      unused.assays = setdiff(all.assays, assay)
      if(length(unused.assays) > 0){
        for (ay in unused.assays){
          # https://github.com/JiekaiLab/dior/blob/2b1ea47b6661c8a10d9455f3baeeccb8f12be2f0/R/seuratIO.R#L65
          # https://github.com/satijalab/seurat-object/blob/58bf437fe058dd78913d9ef7b48008a3e24a306a/R/assay.R#L157
          assay.data <- Seurat::GetAssayData(object =  seu.scdior[[ay]], slot = 'counts')
          seu[[ay]] <- Seurat::CreateAssayObject(counts = assay.data )
        }
      }
      # add graphs
      seu@graphs = seu.scdior@graphs
    }
  }else if(method == "sceasy"){
    seu = tryCatch(
      {
        sceasy::convertFormat(anndata.file, from="anndata", to="seurat",
                              main_layer = slot, assay = assay)
      },
      error = function(cond) {
        message("There is an error when using sceasy: ", cond)
      }
    )
  }else if(method == "scDIOR"){
    seu = tryCatch(
      {
        dior::read_h5ad(file = anndata.file, assay_name=assay, target.object = "seurat")
      },
      error = function(cond) {
        message("There is an error when using scDIOR: ", cond)
      }
    )
  }else if(method == "schard"){
    seu = tryCatch(
      {
        schard::h5ad2seurat(file = anndata.file, use.raw = use.raw, assay = assay)
      },
      error = function(cond) {
        message("There is an error when using schard: ", cond)
      }
    )
  }
  return(seu)
}
```

#### SeuratDisk¶

**Retained information** (`AnnData -> SeuratObject`):

- count matrix: scaled data (`X -> scale.data`), log-normalized data/raw count matrix (`raw.X -> data/counts`)
- cells' meta-information (`obs -> meta.data`)
- annotation of features (`var -> meta.features`)
- dimensional reduction results (`obsm -> reductions`)
- feature loadings (`varm -> reductions`)
- unstructured annotation (`uns -> misc`)

In [89]:

```
%%R
# when raw count matrix stored in adata.raw, the counts and data will be raw count matrix
ann.seu = AD2Seu(anndata.file = "./write/pbmc3k.h5ad", 
                 method = "SeuratDisk", assay="RNA", load.assays = c("RNA"))
ann.seu
```

```
R[write to console]: Warning:
R[write to console]:  Unknown file type: h5ad

R[write to console]: Creating h5Seurat file for version 3.1.5.9900

R[write to console]: Adding X as scale.data

R[write to console]: Adding raw/X as data

R[write to console]: Adding raw/X as counts

R[write to console]: Adding meta.features from raw/var

R[write to console]: Adding dispersions from scaled feature-level metadata

R[write to console]: Adding dispersions_norm from scaled feature-level metadata

R[write to console]: Merging gene_ids from scaled feature-level metadata

R[write to console]: Adding highly_variable from scaled feature-level metadata

R[write to console]: Adding mean from scaled feature-level metadata

R[write to console]: Merging mean_counts from scaled feature-level metadata

R[write to console]: Adding means from scaled feature-level metadata

R[write to console]: Merging mt from scaled feature-level metadata

R[write to console]: Merging n_cells from scaled feature-level metadata

R[write to console]: Merging n_cells_by_counts from scaled feature-level metadata

R[write to console]: Merging pct_dropout_by_counts from scaled feature-level metadata

R[write to console]: Adding std from scaled feature-level metadata

R[write to console]: Merging total_counts from scaled feature-level metadata

R[write to console]: Adding X_pca as cell embeddings for pca

R[write to console]: Adding X_umap as cell embeddings for umap

R[write to console]: Adding PCs as feature loadings fpr pca

R[write to console]: Adding miscellaneous information for pca

R[write to console]: Adding standard deviations for pca

R[write to console]: Adding miscellaneous information for umap

R[write to console]: Adding hvg to miscellaneous data

R[write to console]: Adding leiden to miscellaneous data

R[write to console]: Adding log1p to miscellaneous data

R[write to console]: Adding rank_genes_groups to miscellaneous data

R[write to console]: Adding layer logcounts as data in assay logcounts

R[write to console]: Adding layer rawcounts as data in assay rawcounts

R[write to console]: Validating h5Seurat file

R[write to console]: Warning:
R[write to console]:  Feature names cannot have underscores ('_'), replacing with dashes ('-')

R[write to console]: Initializing RNA with data

R[write to console]: Adding counts for RNA

R[write to console]: Adding scale.data for RNA

R[write to console]: Adding feature-level metadata for RNA

R[write to console]: Adding reduction pca

R[write to console]: Adding cell embeddings for pca

R[write to console]: Adding feature loadings for pca

R[write to console]: Adding miscellaneous information for pca

R[write to console]: Adding reduction umap

R[write to console]: Adding cell embeddings for umap

R[write to console]: Adding miscellaneous information for umap

R[write to console]: Adding command information

R[write to console]: Adding cell-level metadata
```

```
An object of class Seurat 
13714 features across 2638 samples within 1 assay 
Active assay: RNA (13714 features, 0 variable features)
 2 dimensional reductions calculated: pca, umap
```

In [90]:

```
%%R
# raw count matrix
ann.seu@assays$RNA@counts[1:10,1:15]
```

```
10 x 15 sparse Matrix of class "dgCMatrix"
```

```
R[write to console]:   [[ suppressing 15 column names ‘AAACATACAACCAC-1’, ‘AAACATTGAGCTAC-1’, ‘AAACATTGATCAGC-1’ ... ]]
```

```
AL627309.1    . . . . . . . . . . . . . . .
AP006222.2    . . . . . . . . . . . . . . .
RP11-206L10.2 . . . . . . . . . . . . . . .
RP11-206L10.9 . . . . . . . . . . . . . . .
LINC00115     . . . . . . . . . . . . . . .
NOC2L         . . . . . . . . . . . 1 . . .
KLHL17        . . . . . . . . . . . . . . .
PLEKHN1       . . . . . . . . . . . . . . .
RP11-54O7.17  . . . . . . . . . . . . . . .
HES4          . . . . . . . . . . . . . . .
```

In [91]:

```
%%R
# shape
dim(ann.seu@assays$RNA@counts)
```

```
[1] 13714  2638
```

In [92]:

```
%%R
# the data slot contains raw count matrix
ann.seu@assays$RNA@data[1:10,1:15]
```

```
10 x 15 sparse Matrix of class "dgCMatrix"
```

```
R[write to console]:   [[ suppressing 15 column names ‘AAACATACAACCAC-1’, ‘AAACATTGAGCTAC-1’, ‘AAACATTGATCAGC-1’ ... ]]
```

```
AL627309.1    . . . . . . . . . . . . . . .
AP006222.2    . . . . . . . . . . . . . . .
RP11-206L10.2 . . . . . . . . . . . . . . .
RP11-206L10.9 . . . . . . . . . . . . . . .
LINC00115     . . . . . . . . . . . . . . .
NOC2L         . . . . . . . . . . . 1 . . .
KLHL17        . . . . . . . . . . . . . . .
PLEKHN1       . . . . . . . . . . . . . . .
RP11-54O7.17  . . . . . . . . . . . . . . .
HES4          . . . . . . . . . . . . . . .
```

In [93]:

```
%%R
# shape
dim(ann.seu@assays$RNA@data)
```

```
[1] 13714  2638
```

In [94]:

```
%%R
# scaled count matrix
ann.seu@assays$RNA@scale.data[1:10,1:4]
```

```
         AAACATACAACCAC-1 AAACATTGAGCTAC-1 AAACATTGATCAGC-1 AAACCGTGCTTCCG-1
TNFRSF4       -0.17146961      -0.21458235      -0.37688771      -0.28524107
CPSF3L        -0.28081229      -0.37265328      -0.29508454      -0.28173482
ATAD3C        -0.04667677      -0.05480441      -0.05752748      -0.05222671
C1orf86       -0.47516865      -0.68339121      -0.52097195      -0.48492861
RER1          -0.54402399       0.63395083       1.33264792       1.57267952
TNFRSF25       4.92849684      -0.33483663      -0.30936241      -0.27182469
TNFRSF9       -0.03802770      -0.04558870      -0.10310833      -0.07455204
CTNNBIP1      -0.28057277      -0.49826378      -0.27252606      -0.25887546
SRM           -0.34178808      -0.54191375      -0.50079864      -0.41675180
UBIAD1        -0.19536127      -0.20901665      -0.22022836      -0.20847099
```

In [95]:

```
%%R
# shape
dim(ann.seu@assays$RNA@scale.data)
```

```
[1] 1838 2638
```

In [96]:

```
%%R
suppressMessages(library(tidyverse))
# cell's meta-information
ann.seu@meta.data %>% head()
```

```
                 n_genes n_genes_by_counts total_counts total_counts_mt
AAACATACAACCAC-1     781               779         2419              73
AAACATTGAGCTAC-1    1352              1352         4903             186
AAACATTGATCAGC-1    1131              1129         3147              28
AAACCGTGCTTCCG-1     960               960         2639              46
AAACCGTGTATGCG-1     522               521          980              12
AAACGCACTGGTAC-1     782               781         2163              36
                 pct_counts_mt leiden
AAACATACAACCAC-1     3.0177760      0
AAACATTGAGCTAC-1     3.7935958      2
AAACATTGATCAGC-1     0.8897362      0
AAACCGTGCTTCCG-1     1.7430845      4
AAACCGTGTATGCG-1     1.2244898      5
AAACGCACTGGTAC-1     1.6643550      0
```

In [97]:

```
%%R
# annotation of features
ann.seu@assays$RNA@meta.features %>% head()
```

```
                     gene_ids n_cells    mt n_cells_by_counts mean_counts
AL627309.1    ENSG00000237683       9 FALSE                 9 0.003333333
AP006222.2    ENSG00000228463       3 FALSE                 3 0.001111111
RP11-206L10.2 ENSG00000228327       5 FALSE                 5 0.001851852
RP11-206L10.9 ENSG00000237491       3 FALSE                 3 0.001111111
LINC00115     ENSG00000225880      18 FALSE                18 0.006666667
NOC2L         ENSG00000188976     258 FALSE               258 0.106666669
              pct_dropout_by_counts total_counts dispersions dispersions_norm
AL627309.1                 99.66667            9           0                0
AP006222.2                 99.88889            3           0                0
RP11-206L10.2              99.81481            5           0                0
RP11-206L10.9              99.88889            3           0                0
LINC00115                  99.33333           18           0                0
NOC2L                      90.44444          288           0                0
              highly_variable mean means std
AL627309.1              FALSE    0     0   0
AP006222.2              FALSE    0     0   0
RP11-206L10.2           FALSE    0     0   0
RP11-206L10.9           FALSE    0     0   0
LINC00115               FALSE    0     0   0
NOC2L                   FALSE    0     0   0
```

In [98]:

```
%%R
# dimensional reduction results of pca
ann.seu@reductions$pca@cell.embeddings[1:5, 1:5]
```

```
                      PC_1       PC_2       PC_3       PC_4        PC_5
AAACATACAACCAC-1 -5.556221 -0.2577271  0.1867943 -2.8000970  0.05072495
AAACATTGAGCTAC-1 -7.209527 -7.4820013 -0.1627175  8.0185165 -3.00661612
AAACATTGATCAGC-1 -2.694437  1.5836617  0.6631235 -2.2056429  1.78901792
AAACCGTGCTTCCG-1 10.143297  1.3685347 -1.2098237  0.7000697  2.90616465
AAACCGTGTATGCG-1  1.112813  8.1527987 -1.3323525  4.2524910 -1.96318078
```

In [99]:

```
%%R
# pca feature loadings
ann.seu@reductions$pca@feature.loadings[1:5, 1:5]
```

```
                PC_1         PC_2          PC_3         PC_4          PC_5
TNFRSF4 -0.026014818  0.003254168  0.0018978898 -0.036262553  0.0167816579
CPSF3L  -0.008278225  0.009083163 -0.0007814111  0.008882667 -0.0063649304
ATAD3C  -0.003315187  0.003209684  0.0002798587 -0.001740866 -0.0003630426
C1orf86  0.010650732 -0.000268140 -0.0070081116  0.002366116 -0.0038789101
RER1     0.013711839  0.027387908 -0.0107853832  0.006192871  0.0182563197
```

In [100]:

```
%%R
# dimensional reduction results of umap
ann.seu@reductions$umap@cell.embeddings[1:5, 1:2]
```

```
                    umap_1     umap_2
AAACATACAACCAC-1  7.906657  3.5560911
AAACATTGAGCTAC-1  9.248348 12.5443316
AAACATTGATCAGC-1  7.629986  3.8347855
AAACCGTGCTTCCG-1  0.131578  5.5397391
AAACCGTGTATGCG-1 10.055341 -0.6474292
```

In [101]:

```
%%R
# miscellaneous information
ann.seu@reductions$pca@misc
```

```
$params
  use_highly_variable zero_center
1                TRUE        TRUE

$variance
 [1] 32.110455 18.718655 15.607329 13.235289  4.802269  3.985932  3.526233
 [8]  3.233445  3.121209  3.075261  2.998075  2.959521  2.951785  2.944248
[15]  2.913872  2.899030  2.880682  2.864685  2.843064  2.835751  2.831421
[22]  2.818236  2.803552  2.799987  2.788954  2.778101  2.770577  2.760221
[29]  2.753860  2.745955  2.737186  2.734127  2.722202  2.712311  2.702478
[36]  2.700047  2.683850  2.679051  2.676908  2.673995  2.664854  2.657311
[43]  2.651177  2.641776  2.632970  2.629520  2.624529  2.618376  2.618003
[50]  2.601866

$variance_ratio
 [1] 0.020128191 0.011733645 0.009783334 0.008296438 0.003010265 0.002498551
 [7] 0.002210392 0.002026860 0.001956505 0.001927704 0.001879320 0.001855153
[13] 0.001850304 0.001845579 0.001826538 0.001817235 0.001805733 0.001795706
[19] 0.001782152 0.001777568 0.001774855 0.001766590 0.001757385 0.001755151
[25] 0.001748234 0.001741431 0.001736715 0.001730223 0.001726236 0.001721281
[31] 0.001715784 0.001713866 0.001706391 0.001700191 0.001694027 0.001692504
[37] 0.001682351 0.001679342 0.001677999 0.001676173 0.001670443 0.001665715
[43] 0.001661870 0.001655977 0.001650457 0.001648294 0.001645166 0.001641309
[49] 0.001641075 0.001630959
```

In [102]:

```
%%R
# miscellaneous information
ann.seu@reductions$umap@misc
```

```
$params
        a        b
1 0.58303 1.334167
```

#### sceasy¶

**Retained information** (`AnnData -> SeuratObject`):

- count matrix:
  - `slot = "scale.data"`: scaled data (`X -> scale.data`, **1838 x 2638**), raw count matrix (`raw.X -> data`, **13714 x 2638**), `NULL -> counts`
  - `slot = "data"`: scaled count matrix (`X -> data`, 1838 x 2638), raw count matrix (`raw.X -> counts`, **1838 x 2638**), `NULL -> scale.data`
  - `slot = "counts"`: scaled count matrix (`X -> counts`, 1838 x 2638), scaled count matrix (`X -> data`, 1838 x 2638), `NULL -> scale.data`
- cells' meta-information (`obs -> meta.data`)
- annotation of features (`var -> meta.features`), missing *dispersions, dispersions\_norm, highly\_variable, mean, means, std*
- dimensional reduction results (`obsm -> reductions`)

In [103]:

```
%%R
# adata.raw is raw count matrix
# slot = "scale.data": X -> scale.data (1838 x 2638), raw.X -> data (raw count matrix, 13714 x 2638), counts is empty
# slot = "data": X -> data (scaled count matrix, 1838 x 2638), raw.X -> counts (raw count matrix, 1838 x 2638), scale.data is empty
# slot = "counts": X -> counts (scaled count matrix, 1838 x 2638), data is scaled count matrix (1838 x 2638), scale.data is empty
ann.sceasy = AD2Seu(anndata.file = "./write/pbmc3k.h5ad",
                    method = "sceasy", assay="RNA", slot = "scale.data")
ann.sceasy
```

```
R[write to console]: Warning:
R[write to console]:  Feature names cannot have underscores ('_'), replacing with dashes ('-')

R[write to console]: X -> scale.data; raw.X -> data
```

```
An object of class Seurat 
13714 features across 2638 samples within 1 assay 
Active assay: RNA (13714 features, 0 variable features)
 2 dimensional reductions calculated: pca, umap
```

In [104]:

```
%%R
# counts is empty
ann.sceasy@assays$RNA@counts
```

```
<0 x 0 matrix>
```

In [105]:

```
%%R
# the data slot contains raw count matrix
ann.sceasy@assays$RNA@data[1:10,1:15]
```

```
10 x 15 sparse Matrix of class "dgCMatrix"
```

```
R[write to console]:   [[ suppressing 15 column names ‘AAACATACAACCAC-1’, ‘AAACATTGAGCTAC-1’, ‘AAACATTGATCAGC-1’ ... ]]
```

```
AL627309.1    . . . . . . . . . . . . . . .
AP006222.2    . . . . . . . . . . . . . . .
RP11-206L10.2 . . . . . . . . . . . . . . .
RP11-206L10.9 . . . . . . . . . . . . . . .
LINC00115     . . . . . . . . . . . . . . .
NOC2L         . . . . . . . . . . . 1 . . .
KLHL17        . . . . . . . . . . . . . . .
PLEKHN1       . . . . . . . . . . . . . . .
RP11-54O7.17  . . . . . . . . . . . . . . .
HES4          . . . . . . . . . . . . . . .
```

In [106]:

```
%%R
# shape
dim(ann.sceasy@assays$RNA@data)
```

```
[1] 13714  2638
```

In [107]:

```
%%R
# scaled count matrix
ann.sceasy@assays$RNA@scale.data[1:10,1:4]
```

```
         AAACATACAACCAC-1 AAACATTGAGCTAC-1 AAACATTGATCAGC-1 AAACCGTGCTTCCG-1
TNFRSF4       -0.17146961      -0.21458235      -0.37688771      -0.28524107
CPSF3L        -0.28081229      -0.37265328      -0.29508454      -0.28173482
ATAD3C        -0.04667677      -0.05480441      -0.05752748      -0.05222671
C1orf86       -0.47516865      -0.68339121      -0.52097195      -0.48492861
RER1          -0.54402399       0.63395083       1.33264792       1.57267952
TNFRSF25       4.92849684      -0.33483663      -0.30936241      -0.27182469
TNFRSF9       -0.03802770      -0.04558870      -0.10310833      -0.07455204
CTNNBIP1      -0.28057277      -0.49826378      -0.27252606      -0.25887546
SRM           -0.34178808      -0.54191375      -0.50079864      -0.41675180
UBIAD1        -0.19536127      -0.20901665      -0.22022836      -0.20847099
```

In [108]:

```
%%R
# shape
dim(ann.sceasy@assays$RNA@scale.data)
```

```
[1] 1838 2638
```

In [109]:

```
%%R
suppressMessages(library(tidyverse))
# cell's meta-information
ann.sceasy@meta.data %>% head()
```

```
                 nFeaturess_RNA nFeaturess_RNA_by_counts total_counts
AAACATACAACCAC-1            781                      779         2419
AAACATTGAGCTAC-1           1352                     1352         4903
AAACATTGATCAGC-1           1131                     1129         3147
AAACCGTGCTTCCG-1            960                      960         2639
AAACCGTGTATGCG-1            522                      521          980
AAACGCACTGGTAC-1            782                      781         2163
                 total_counts_mt pct_counts_mt leiden
AAACATACAACCAC-1              73     3.0177760      0
AAACATTGAGCTAC-1             186     3.7935958      2
AAACATTGATCAGC-1              28     0.8897362      0
AAACCGTGCTTCCG-1              46     1.7430845      4
AAACCGTGTATGCG-1              12     1.2244898      5
AAACGCACTGGTAC-1              36     1.6643550      0
```

In [110]:

```
%%R
# annotation of features, missing dispersions, dispersions_norm, highly_variable, mean, means, std
ann.sceasy@assays$RNA@meta.features %>% head()
```

```
                     gene_ids n_cells    mt n_cells_by_counts mean_counts
AL627309.1    ENSG00000237683       9 FALSE                 9 0.003333333
AP006222.2    ENSG00000228463       3 FALSE                 3 0.001111111
RP11-206L10.2 ENSG00000228327       5 FALSE                 5 0.001851852
RP11-206L10.9 ENSG00000237491       3 FALSE                 3 0.001111111
LINC00115     ENSG00000225880      18 FALSE                18 0.006666667
NOC2L         ENSG00000188976     258 FALSE               258 0.106666669
              pct_dropout_by_counts total_counts
AL627309.1                 99.66667            9
AP006222.2                 99.88889            3
RP11-206L10.2              99.81481            5
RP11-206L10.9              99.88889            3
LINC00115                  99.33333           18
NOC2L                      90.44444          288
```

In [111]:

```
%%R
# dimensional reduction results of pca
ann.sceasy@reductions$pca@cell.embeddings[1:5, 1:5]
```

```
                      PC_1       PC_2       PC_3       PC_4        PC_5
AAACATACAACCAC-1 -5.556221 -0.2577271  0.1867943 -2.8000970  0.05072495
AAACATTGAGCTAC-1 -7.209527 -7.4820013 -0.1627175  8.0185165 -3.00661612
AAACATTGATCAGC-1 -2.694437  1.5836617  0.6631235 -2.2056429  1.78901792
AAACCGTGCTTCCG-1 10.143297  1.3685347 -1.2098237  0.7000697  2.90616465
AAACCGTGTATGCG-1  1.112813  8.1527987 -1.3323525  4.2524910 -1.96318078
```

In [112]:

```
%%R
# dimensional reduction results of umap
ann.sceasy@reductions$umap@cell.embeddings[1:5, 1:2]
```

```
                    UMAP_1     UMAP_2
AAACATACAACCAC-1  7.906657  3.5560911
AAACATTGAGCTAC-1  9.248348 12.5443316
AAACATTGATCAGC-1  7.629986  3.8347855
AAACCGTGCTTCCG-1  0.131578  5.5397391
AAACCGTGTATGCG-1 10.055341 -0.6474292
```

#### scDIOR¶

**Retained information** (`AnnData -> SeuratObject`):

- count matrix: scaled data (`X -> scale.data`), log-normalized data/raw count matrix (`raw.X -> data and counts`)
- cells' meta-information (`obs -> meta.data`)
- annotation of features (`var -> meta.features`), missing *dispersions, dispersions\_norm, highly\_variable, mean, means, std*
- dimensional reduction results (`obsm -> reductions`)
- relationship of cells, graphs (`obsp -> graphs`)
- additional assays (`layers -> assays`)

In [113]:

```
%%R
ann.scdior = AD2Seu(anndata.file = "./write/pbmc3k.h5ad",
                    method = "scDIOR", assay="RNA")
ann.scdior
```

```
R[write to console]: Warning:
R[write to console]:  No columnames present in cell embeddings, setting to 'PCA_1:50'

R[write to console]: Warning:
R[write to console]:  No columnames present in cell embeddings, setting to 'UMAP_1:2'

R[write to console]: Warning:
R[write to console]:  Feature names cannot have underscores ('_'), replacing with dashes ('-')
```

```
An object of class Seurat 
17390 features across 2638 samples within 3 assays 
Active assay: RNA (13714 features, 0 variable features)
 2 other assays present: logcounts, rawcounts
 2 dimensional reductions calculated: pca, umap
```

In [114]:

```
%%R
# assays
ann.scdior@assays
```

```
$RNA
Assay data with 13714 features for 2638 cells
First 10 features:
 AL627309.1, AP006222.2, RP11-206L10.2, RP11-206L10.9, LINC00115, NOC2L,
KLHL17, PLEKHN1, RP11-54O7.17, HES4 

$logcounts
Assay data with 1838 features for 2638 cells
First 10 features:
 TNFRSF4, CPSF3L, ATAD3C, C1orf86, RER1, TNFRSF25, TNFRSF9, CTNNBIP1,
SRM, UBIAD1 

$rawcounts
Assay data with 1838 features for 2638 cells
First 10 features:
 TNFRSF4, CPSF3L, ATAD3C, C1orf86, RER1, TNFRSF25, TNFRSF9, CTNNBIP1,
SRM, UBIAD1
```

In [115]:

```
%%R
# raw count matrix
ann.scdior@assays$RNA@counts[1:10,1:15]
```

```
10 x 15 sparse Matrix of class "dgCMatrix"
```

```
R[write to console]:   [[ suppressing 15 column names ‘AAACATACAACCAC-1’, ‘AAACATTGAGCTAC-1’, ‘AAACATTGATCAGC-1’ ... ]]
```

```
AL627309.1    . . . . . . . . . . . . . . .
AP006222.2    . . . . . . . . . . . . . . .
RP11-206L10.2 . . . . . . . . . . . . . . .
RP11-206L10.9 . . . . . . . . . . . . . . .
LINC00115     . . . . . . . . . . . . . . .
NOC2L         . . . . . . . . . . . 1 . . .
KLHL17        . . . . . . . . . . . . . . .
PLEKHN1       . . . . . . . . . . . . . . .
RP11-54O7.17  . . . . . . . . . . . . . . .
HES4          . . . . . . . . . . . . . . .
```

In [116]:

```
%%R
# shape
dim(ann.scdior@assays$RNA@counts)
```

```
[1] 13714  2638
```

In [117]:

```
%%R
# the data slot contains raw count matrix
ann.scdior@assays$RNA@data[1:10,1:15]
```

```
10 x 15 sparse Matrix of class "dgCMatrix"
```

```
R[write to console]:   [[ suppressing 15 column names ‘AAACATACAACCAC-1’, ‘AAACATTGAGCTAC-1’, ‘AAACATTGATCAGC-1’ ... ]]
```

```
AL627309.1    . . . . . . . . . . . . . . .
AP006222.2    . . . . . . . . . . . . . . .
RP11-206L10.2 . . . . . . . . . . . . . . .
RP11-206L10.9 . . . . . . . . . . . . . . .
LINC00115     . . . . . . . . . . . . . . .
NOC2L         . . . . . . . . . . . 1 . . .
KLHL17        . . . . . . . . . . . . . . .
PLEKHN1       . . . . . . . . . . . . . . .
RP11-54O7.17  . . . . . . . . . . . . . . .
HES4          . . . . . . . . . . . . . . .
```

In [118]:

```
%%R
# shape
dim(ann.scdior@assays$RNA@data)
```

```
[1] 13714  2638
```

In [119]:

```
%%R
# scaled count matrix
ann.scdior@assays$RNA@scale.data[1:10,1:4]
```

```
         AAACATACAACCAC-1 AAACATTGAGCTAC-1 AAACATTGATCAGC-1 AAACCGTGCTTCCG-1
TNFRSF4       -0.17146961      -0.21458235      -0.37688771      -0.28524107
CPSF3L        -0.28081229      -0.37265328      -0.29508454      -0.28173482
ATAD3C        -0.04667677      -0.05480441      -0.05752748      -0.05222671
C1orf86       -0.47516865      -0.68339121      -0.52097195      -0.48492861
RER1          -0.54402399       0.63395083       1.33264792       1.57267952
TNFRSF25       4.92849684      -0.33483663      -0.30936241      -0.27182469
TNFRSF9       -0.03802770      -0.04558870      -0.10310833      -0.07455204
CTNNBIP1      -0.28057277      -0.49826378      -0.27252606      -0.25887546
SRM           -0.34178808      -0.54191375      -0.50079864      -0.41675180
UBIAD1        -0.19536127      -0.20901665      -0.22022836      -0.20847099
```

In [120]:

```
%%R
# shape
dim(ann.scdior@assays$RNA@scale.data)
```

```
[1] 1838 2638
```

In [121]:

```
%%R
suppressMessages(library(tidyverse))
# cell's meta-information
ann.scdior@meta.data %>% head()
```

```
                 n_genes n_genes_by_counts total_counts total_counts_mt
AAACATACAACCAC-1     781               779         2419              73
AAACATTGAGCTAC-1    1352              1352         4903             186
AAACATTGATCAGC-1    1131              1129         3147              28
AAACCGTGCTTCCG-1     960               960         2639              46
AAACCGTGTATGCG-1     522               521          980              12
AAACGCACTGGTAC-1     782               781         2163              36
                 pct_counts_mt leiden
AAACATACAACCAC-1     3.0177760      0
AAACATTGAGCTAC-1     3.7935958      2
AAACATTGATCAGC-1     0.8897362      0
AAACCGTGCTTCCG-1     1.7430845      4
AAACCGTGTATGCG-1     1.2244898      5
AAACGCACTGGTAC-1     1.6643550      0
```

In [122]:

```
%%R
# annotation of features, missing dispersions, dispersions_norm, highly_variable, mean, means, std
ann.scdior@assays$RNA@meta.features %>% head()
```

```
                     gene_ids n_cells    mt n_cells_by_counts mean_counts
AL627309.1    ENSG00000237683       9 FALSE                 9 0.003333333
AP006222.2    ENSG00000228463       3 FALSE                 3 0.001111111
RP11-206L10.2 ENSG00000228327       5 FALSE                 5 0.001851852
RP11-206L10.9 ENSG00000237491       3 FALSE                 3 0.001111111
LINC00115     ENSG00000225880      18 FALSE                18 0.006666667
NOC2L         ENSG00000188976     258 FALSE               258 0.106666669
              pct_dropout_by_counts total_counts
AL627309.1                 99.66667            9
AP006222.2                 99.88889            3
RP11-206L10.2              99.81481            5
RP11-206L10.9              99.88889            3
LINC00115                  99.33333           18
NOC2L                      90.44444          288
```

In [123]:

```
%%R
# dimensional reduction results of pca
ann.scdior@reductions$pca@cell.embeddings[1:5, 1:5]
```

```
                     PCA_1      PCA_2      PCA_3      PCA_4       PCA_5
AAACATACAACCAC-1 -5.556221 -0.2577271  0.1867943 -2.8000970  0.05072495
AAACATTGAGCTAC-1 -7.209527 -7.4820013 -0.1627175  8.0185165 -3.00661612
AAACATTGATCAGC-1 -2.694437  1.5836617  0.6631235 -2.2056429  1.78901792
AAACCGTGCTTCCG-1 10.143297  1.3685347 -1.2098237  0.7000697  2.90616465
AAACCGTGTATGCG-1  1.112813  8.1527987 -1.3323525  4.2524910 -1.96318078
```

In [124]:

```
%%R
# dimensional reduction results of umap
ann.scdior@reductions$umap@cell.embeddings[1:5, 1:2]
```

```
                    UMAP_1     UMAP_2
AAACATACAACCAC-1  7.906657  3.5560911
AAACATTGAGCTAC-1  9.248348 12.5443316
AAACATTGATCAGC-1  7.629986  3.8347855
AAACCGTGCTTCCG-1  0.131578  5.5397391
AAACCGTGTATGCG-1 10.055341 -0.6474292
```

In [125]:

```
%%R
# relationship of cells, graphs
# RNA_nn
ann.scdior@graphs$RNA_nn[1:10,1:10]
```

```
10 x 10 sparse Matrix of class "dgCMatrix"
```

```
R[write to console]:   [[ suppressing 10 column names ‘AAACATACAACCAC-1’, ‘AAACATTGAGCTAC-1’, ‘AAACATTGATCAGC-1’ ... ]]
```

```
AAACATACAACCAC-1 . . . . . . . . . .
AAACATTGAGCTAC-1 . . . . . . . . . .
AAACATTGATCAGC-1 . . . . . . . . . .
AAACCGTGCTTCCG-1 . . . . . . . . . .
AAACCGTGTATGCG-1 . . . . . . . . . .
AAACGCACTGGTAC-1 . . . . . . . . . .
AAACGCTGACCAGT-1 . . . . . . . . . .
AAACGCTGGTTCTT-1 . . . . . . . . . .
AAACGCTGTAGCCA-1 . . . . . . . . . .
AAACGCTGTTTCTG-1 . . . . . . . . . .
```

In [126]:

```
%%R
# relationship of cells, graphs
# RNA_snn
ann.scdior@graphs$RNA_snn[1:10,1:10]
```

```
10 x 10 sparse Matrix of class "dgCMatrix"
```

```
R[write to console]:   [[ suppressing 10 column names ‘AAACATACAACCAC-1’, ‘AAACATTGAGCTAC-1’, ‘AAACATTGATCAGC-1’ ... ]]
```

```
AAACATACAACCAC-1 . . . . . . . . . .
AAACATTGAGCTAC-1 . . . . . . . . . .
AAACATTGATCAGC-1 . . . . . . . . . .
AAACCGTGCTTCCG-1 . . . . . . . . . .
AAACCGTGTATGCG-1 . . . . . . . . . .
AAACGCACTGGTAC-1 . . . . . . . . . .
AAACGCTGACCAGT-1 . . . . . . . . . .
AAACGCTGGTTCTT-1 . . . . . . . . . .
AAACGCTGTAGCCA-1 . . . . . . . . . .
AAACGCTGTTTCTG-1 . . . . . . . . . .
```

In [127]:

```
%%R
dim(ann.scdior@graphs$RNA_snn)
```

```
[1] 2638 2638
```

#### schard¶

**Retained information** (`AnnData -> SeuratObject`):

- count matrix:
  - `use.raw = TRUE`: raw count matrix (`raw.X -> counts/data`, **13714 x 2638**), `NULL -> scale.data`
  - `use.raw = FALSE`: scaled count matrix (`X -> data`, 1838 x 2638), `NULL -> scale.data and counts`
- cells' meta-information (`obs -> meta.data`)
- annotation of features (`var -> meta.features`), missing *dispersions, dispersions\_norm, highly\_variable, mean, means, std* when `use.raw = TRUE`
- dimensional reduction results (`obsm -> reductions`)

In [128]:

```
%%R
# use.raw = TRUE: meta.features missing dispersions, dispersions_norm, highly_variable, mean, means, std
ann.schard = AD2Seu(anndata.file = "./write/pbmc3k.h5ad",
                    method = "schard", assay="RNA", use.raw = TRUE)
ann.schard
```

```
R[write to console]: Warning:
R[write to console]:  Keys should be one or more alphanumeric characters followed by an underscore, setting key from rna to rna_

R[write to console]: Warning:
R[write to console]:  Invalid name supplied, making object name syntactically valid. New object name is X_indexn_genesn_genes_by_countstotal_countstotal_counts_mtpct_counts_mtleiden; see ?make.names for more details on syntax validity
```

```
An object of class Seurat 
13714 features across 2638 samples within 1 assay 
Active assay: RNA (13714 features, 0 variable features)
 2 dimensional reductions calculated: Xpca_, Xumap_
```

In [129]:

```
%%R
# raw count matrix
ann.schard@assays$RNA@counts[1:10,1:15]
```

```
10 x 15 sparse Matrix of class "dgCMatrix"
```

```
R[write to console]:   [[ suppressing 15 column names ‘AAACATACAACCAC-1’, ‘AAACATTGAGCTAC-1’, ‘AAACATTGATCAGC-1’ ... ]]
```

```
AL627309.1    . . . . . . . . . . . . . . .
AP006222.2    . . . . . . . . . . . . . . .
RP11-206L10.2 . . . . . . . . . . . . . . .
RP11-206L10.9 . . . . . . . . . . . . . . .
LINC00115     . . . . . . . . . . . . . . .
NOC2L         . . . . . . . . . . . 1 . . .
KLHL17        . . . . . . . . . . . . . . .
PLEKHN1       . . . . . . . . . . . . . . .
RP11-54O7.17  . . . . . . . . . . . . . . .
HES4          . . . . . . . . . . . . . . .
```

In [130]:

```
%%R
# shape
dim(ann.schard@assays$RNA@counts)
```

```
[1] 13714  2638
```

In [131]:

```
%%R
# the data slot contains raw count matrix
ann.schard@assays$RNA@data[1:10,1:15]
```

```
10 x 15 sparse Matrix of class "dgCMatrix"
```

```
R[write to console]:   [[ suppressing 15 column names ‘AAACATACAACCAC-1’, ‘AAACATTGAGCTAC-1’, ‘AAACATTGATCAGC-1’ ... ]]
```

```
AL627309.1    . . . . . . . . . . . . . . .
AP006222.2    . . . . . . . . . . . . . . .
RP11-206L10.2 . . . . . . . . . . . . . . .
RP11-206L10.9 . . . . . . . . . . . . . . .
LINC00115     . . . . . . . . . . . . . . .
NOC2L         . . . . . . . . . . . 1 . . .
KLHL17        . . . . . . . . . . . . . . .
PLEKHN1       . . . . . . . . . . . . . . .
RP11-54O7.17  . . . . . . . . . . . . . . .
HES4          . . . . . . . . . . . . . . .
```

In [132]:

```
%%R
# shape
dim(ann.schard@assays$RNA@data)
```

```
[1] 13714  2638
```

In [133]:

```
%%R
# scaled count matrix is empty
ann.schard@assays$RNA@scale.data
```

```
<0 x 0 matrix>
```

In [134]:

```
%%R
suppressMessages(library(tidyverse))
# cell's meta-information
ann.schard@meta.data %>% head()
```

```
                    orig.ident nCount_RNA nFeature_RNA          X_index n_genes
AAACATACAACCAC-1 SeuratProject       2419          779 AAACATACAACCAC-1     781
AAACATTGAGCTAC-1 SeuratProject       4903         1352 AAACATTGAGCTAC-1    1352
AAACATTGATCAGC-1 SeuratProject       3147         1129 AAACATTGATCAGC-1    1131
AAACCGTGCTTCCG-1 SeuratProject       2639          960 AAACCGTGCTTCCG-1     960
AAACCGTGTATGCG-1 SeuratProject        980          521 AAACCGTGTATGCG-1     522
AAACGCACTGGTAC-1 SeuratProject       2163          781 AAACGCACTGGTAC-1     782
                 n_genes_by_counts total_counts total_counts_mt pct_counts_mt
AAACATACAACCAC-1               779         2419              73     3.0177760
AAACATTGAGCTAC-1              1352         4903             186     3.7935958
AAACATTGATCAGC-1              1129         3147              28     0.8897362
AAACCGTGCTTCCG-1               960         2639              46     1.7430845
AAACCGTGTATGCG-1               521          980              12     1.2244898
AAACGCACTGGTAC-1               781         2163              36     1.6643550
                 leiden
AAACATACAACCAC-1      0
AAACATTGAGCTAC-1      2
AAACATTGATCAGC-1      0
AAACCGTGCTTCCG-1      4
AAACCGTGTATGCG-1      5
AAACGCACTGGTAC-1      0
```

In [135]:

```
%%R
# annotation of features, missing dispersions, dispersions_norm, highly_variable, mean, means, std (use.raw = TRUE)
ann.schard@assays$RNA@meta.features %>% head()
```

```
                     _index        gene_ids n_cells    mt n_cells_by_counts
AL627309.1       AL627309.1 ENSG00000237683       9 FALSE                 9
AP006222.2       AP006222.2 ENSG00000228463       3 FALSE                 3
RP11-206L10.2 RP11-206L10.2 ENSG00000228327       5 FALSE                 5
RP11-206L10.9 RP11-206L10.9 ENSG00000237491       3 FALSE                 3
LINC00115         LINC00115 ENSG00000225880      18 FALSE                18
NOC2L                 NOC2L ENSG00000188976     258 FALSE               258
              mean_counts pct_dropout_by_counts total_counts
AL627309.1    0.003333333              99.66667            9
AP006222.2    0.001111111              99.88889            3
RP11-206L10.2 0.001851852              99.81481            5
RP11-206L10.9 0.001111111              99.88889            3
LINC00115     0.006666667              99.33333           18
NOC2L         0.106666669              90.44444          288
```

In [136]:

```
%%R
# dimensional reduction results of pca
ann.schard@reductions$Xpca_@cell.embeddings[1:5, 1:5]
```

```
                    Xpca_1     Xpca_2     Xpca_3     Xpca_4      Xpca_5
AAACATACAACCAC-1 -5.556221 -0.2577271  0.1867943 -2.8000970  0.05072495
AAACATTGAGCTAC-1 -7.209527 -7.4820013 -0.1627175  8.0185165 -3.00661612
AAACATTGATCAGC-1 -2.694437  1.5836617  0.6631235 -2.2056429  1.78901792
AAACCGTGCTTCCG-1 10.143297  1.3685347 -1.2098237  0.7000697  2.90616465
AAACCGTGTATGCG-1  1.112813  8.1527987 -1.3323525  4.2524910 -1.96318078
```

In [137]:

```
%%R
# dimensional reduction results of umap
ann.schard@reductions$Xumap_@cell.embeddings[1:5, 1:2]
```

```
                   Xumap_1    Xumap_2
AAACATACAACCAC-1  7.906657  3.5560911
AAACATTGAGCTAC-1  9.248348 12.5443316
AAACATTGATCAGC-1  7.629986  3.8347855
AAACCGTGCTTCCG-1  0.131578  5.5397391
AAACCGTGTATGCG-1 10.055341 -0.6474292
```

#### SeuratDisk + scDIOR¶

**Retained information** (`AnnData -> SeuratObject`):

- count matrix: scaled data (`X -> scale.data`), log-normalized data/raw count matrix (`raw.X -> data/counts`)
- cells' meta-information (`obs -> meta.data`)
- annotation of features (`var -> meta.features`)
- dimensional reduction results (`obsm -> reductions`)
- feature loadings (`varm -> reductions`)
- unstructured annotation (`uns -> misc`)
- relationship of cells, graphs (`obsp -> graphs`) (`scDIOR`)
- additional assays (`layers -> assays`) (`scDIOR`)

In [138]:

```
%%R
# add graphs and additional assays from scDIOR to SeuratDisk
ann.seuscdior = AD2Seu(anndata.file = "./write/pbmc3k.h5ad",
                       method = "SeuratDisk+scDIOR", assay="RNA", load.assays = c("RNA"))
ann.seuscdior
```

```
R[write to console]: Warning:
R[write to console]:  Unknown file type: h5ad

R[write to console]: Creating h5Seurat file for version 3.1.5.9900

R[write to console]: Adding X as scale.data

R[write to console]: Adding raw/X as data

R[write to console]: Adding raw/X as counts

R[write to console]: Adding meta.features from raw/var

R[write to console]: Adding dispersions from scaled feature-level metadata

R[write to console]: Adding dispersions_norm from scaled feature-level metadata

R[write to console]: Merging gene_ids from scaled feature-level metadata

R[write to console]: Adding highly_variable from scaled feature-level metadata

R[write to console]: Adding mean from scaled feature-level metadata

R[write to console]: Merging mean_counts from scaled feature-level metadata

R[write to console]: Adding means from scaled feature-level metadata

R[write to console]: Merging mt from scaled feature-level metadata

R[write to console]: Merging n_cells from scaled feature-level metadata

R[write to console]: Merging n_cells_by_counts from scaled feature-level metadata

R[write to console]: Merging pct_dropout_by_counts from scaled feature-level metadata

R[write to console]: Adding std from scaled feature-level metadata

R[write to console]: Merging total_counts from scaled feature-level metadata

R[write to console]: Adding X_pca as cell embeddings for pca

R[write to console]: Adding X_umap as cell embeddings for umap

R[write to console]: Adding PCs as feature loadings fpr pca

R[write to console]: Adding miscellaneous information for pca

R[write to console]: Adding standard deviations for pca

R[write to console]: Adding miscellaneous information for umap

R[write to console]: Adding hvg to miscellaneous data

R[write to console]: Adding leiden to miscellaneous data

R[write to console]: Adding log1p to miscellaneous data

R[write to console]: Adding rank_genes_groups to miscellaneous data

R[write to console]: Adding layer logcounts as data in assay logcounts

R[write to console]: Adding layer rawcounts as data in assay rawcounts

R[write to console]: Validating h5Seurat file

R[write to console]: Warning:
R[write to console]:  Feature names cannot have underscores ('_'), replacing with dashes ('-')

R[write to console]: Initializing RNA with data

R[write to console]: Adding counts for RNA

R[write to console]: Adding scale.data for RNA

R[write to console]: Adding feature-level metadata for RNA

R[write to console]: Adding reduction pca

R[write to console]: Adding cell embeddings for pca

R[write to console]: Adding feature loadings for pca

R[write to console]: Adding miscellaneous information for pca

R[write to console]: Adding reduction umap

R[write to console]: Adding cell embeddings for umap

R[write to console]: Adding miscellaneous information for umap

R[write to console]: Adding command information

R[write to console]: Adding cell-level metadata

R[write to console]: Warning:
R[write to console]:  No columnames present in cell embeddings, setting to 'PCA_1:50'

R[write to console]: Warning:
R[write to console]:  No columnames present in cell embeddings, setting to 'UMAP_1:2'

R[write to console]: Warning:
R[write to console]:  Feature names cannot have underscores ('_'), replacing with dashes ('-')
```

```
An object of class Seurat 
17390 features across 2638 samples within 3 assays 
Active assay: RNA (13714 features, 0 variable features)
 2 other assays present: logcounts, rawcounts
 2 dimensional reductions calculated: pca, umap
```

In [139]:

```
%%R
# assays
ann.seuscdior@assays
```

```
$RNA
Assay data with 13714 features for 2638 cells
First 10 features:
 AL627309.1, AP006222.2, RP11-206L10.2, RP11-206L10.9, LINC00115, NOC2L,
KLHL17, PLEKHN1, RP11-54O7.17, HES4 

$logcounts
Assay data with 1838 features for 2638 cells
First 10 features:
 TNFRSF4, CPSF3L, ATAD3C, C1orf86, RER1, TNFRSF25, TNFRSF9, CTNNBIP1,
SRM, UBIAD1 

$rawcounts
Assay data with 1838 features for 2638 cells
First 10 features:
 TNFRSF4, CPSF3L, ATAD3C, C1orf86, RER1, TNFRSF25, TNFRSF9, CTNNBIP1,
SRM, UBIAD1
```

In [140]:

```
%%R
# raw count matrix
ann.seuscdior@assays$RNA@counts[1:10,1:15]
```

```
10 x 15 sparse Matrix of class "dgCMatrix"
```

```
R[write to console]:   [[ suppressing 15 column names ‘AAACATACAACCAC-1’, ‘AAACATTGAGCTAC-1’, ‘AAACATTGATCAGC-1’ ... ]]
```

```
AL627309.1    . . . . . . . . . . . . . . .
AP006222.2    . . . . . . . . . . . . . . .
RP11-206L10.2 . . . . . . . . . . . . . . .
RP11-206L10.9 . . . . . . . . . . . . . . .
LINC00115     . . . . . . . . . . . . . . .
NOC2L         . . . . . . . . . . . 1 . . .
KLHL17        . . . . . . . . . . . . . . .
PLEKHN1       . . . . . . . . . . . . . . .
RP11-54O7.17  . . . . . . . . . . . . . . .
HES4          . . . . . . . . . . . . . . .
```

In [141]:

```
%%R
# shape
dim(ann.seuscdior@assays$RNA@counts)
```

```
[1] 13714  2638
```

In [142]:

```
%%R
# the data slot contains raw count matrix
ann.seuscdior@assays$RNA@data[1:10,1:15]
```

```
10 x 15 sparse Matrix of class "dgCMatrix"
```

```
R[write to console]:   [[ suppressing 15 column names ‘AAACATACAACCAC-1’, ‘AAACATTGAGCTAC-1’, ‘AAACATTGATCAGC-1’ ... ]]
```

```
AL627309.1    . . . . . . . . . . . . . . .
AP006222.2    . . . . . . . . . . . . . . .
RP11-206L10.2 . . . . . . . . . . . . . . .
RP11-206L10.9 . . . . . . . . . . . . . . .
LINC00115     . . . . . . . . . . . . . . .
NOC2L         . . . . . . . . . . . 1 . . .
KLHL17        . . . . . . . . . . . . . . .
PLEKHN1       . . . . . . . . . . . . . . .
RP11-54O7.17  . . . . . . . . . . . . . . .
HES4          . . . . . . . . . . . . . . .
```

In [143]:

```
%%R
# shape
dim(ann.seuscdior@assays$RNA@data)
```

```
[1] 13714  2638
```

In [144]:

```
%%R
# scaled count matrix
ann.seuscdior@assays$RNA@scale.data[1:10,1:4]
```

```
         AAACATACAACCAC-1 AAACATTGAGCTAC-1 AAACATTGATCAGC-1 AAACCGTGCTTCCG-1
TNFRSF4       -0.17146961      -0.21458235      -0.37688771      -0.28524107
CPSF3L        -0.28081229      -0.37265328      -0.29508454      -0.28173482
ATAD3C        -0.04667677      -0.05480441      -0.05752748      -0.05222671
C1orf86       -0.47516865      -0.68339121      -0.52097195      -0.48492861
RER1          -0.54402399       0.63395083       1.33264792       1.57267952
TNFRSF25       4.92849684      -0.33483663      -0.30936241      -0.27182469
TNFRSF9       -0.03802770      -0.04558870      -0.10310833      -0.07455204
CTNNBIP1      -0.28057277      -0.49826378      -0.27252606      -0.25887546
SRM           -0.34178808      -0.54191375      -0.50079864      -0.41675180
UBIAD1        -0.19536127      -0.20901665      -0.22022836      -0.20847099
```

In [145]:

```
%%R
# shape
dim(ann.seuscdior@assays$RNA@scale.data)
```

```
[1] 1838 2638
```

In [146]:

```
%%R
suppressMessages(library(tidyverse))
# cell's meta-information
ann.seuscdior@meta.data %>% head()
```

```
                 n_genes n_genes_by_counts total_counts total_counts_mt
AAACATACAACCAC-1     781               779         2419              73
AAACATTGAGCTAC-1    1352              1352         4903             186
AAACATTGATCAGC-1    1131              1129         3147              28
AAACCGTGCTTCCG-1     960               960         2639              46
AAACCGTGTATGCG-1     522               521          980              12
AAACGCACTGGTAC-1     782               781         2163              36
                 pct_counts_mt leiden nCount_logcounts nFeature_logcounts
AAACATACAACCAC-1     3.0177760      0         255.1598                141
AAACATTGAGCTAC-1     3.7935958      2         350.1350                249
AAACATTGATCAGC-1     0.8897362      0         324.0653                200
AAACCGTGCTTCCG-1     1.7430845      4         361.0839                187
AAACCGTGTATGCG-1     1.2244898      5         246.8949                 91
AAACGCACTGGTAC-1     1.6643550      0         281.6889                140
                 nCount_rawcounts nFeature_rawcounts
AAACATACAACCAC-1              216                141
AAACATTGAGCTAC-1              503                249
AAACATTGATCAGC-1              289                200
AAACCGTGCTTCCG-1              403                187
AAACCGTGTATGCG-1              202                 91
AAACGCACTGGTAC-1              273                140
```

In [147]:

```
%%R
# annotation of features
ann.seuscdior@assays$RNA@meta.features %>% head()
```

```
                     gene_ids n_cells    mt n_cells_by_counts mean_counts
AL627309.1    ENSG00000237683       9 FALSE                 9 0.003333333
AP006222.2    ENSG00000228463       3 FALSE                 3 0.001111111
RP11-206L10.2 ENSG00000228327       5 FALSE                 5 0.001851852
RP11-206L10.9 ENSG00000237491       3 FALSE                 3 0.001111111
LINC00115     ENSG00000225880      18 FALSE                18 0.006666667
NOC2L         ENSG00000188976     258 FALSE               258 0.106666669
              pct_dropout_by_counts total_counts dispersions dispersions_norm
AL627309.1                 99.66667            9           0                0
AP006222.2                 99.88889            3           0                0
RP11-206L10.2              99.81481            5           0                0
RP11-206L10.9              99.88889            3           0                0
LINC00115                  99.33333           18           0                0
NOC2L                      90.44444          288           0                0
              highly_variable mean means std
AL627309.1              FALSE    0     0   0
AP006222.2              FALSE    0     0   0
RP11-206L10.2           FALSE    0     0   0
RP11-206L10.9           FALSE    0     0   0
LINC00115               FALSE    0     0   0
NOC2L                   FALSE    0     0   0
```

In [148]:

```
%%R
# dimensional reduction results of pca
ann.seuscdior@reductions$pca@cell.embeddings[1:5, 1:5]
```

```
                      PC_1       PC_2       PC_3       PC_4        PC_5
AAACATACAACCAC-1 -5.556221 -0.2577271  0.1867943 -2.8000970  0.05072495
AAACATTGAGCTAC-1 -7.209527 -7.4820013 -0.1627175  8.0185165 -3.00661612
AAACATTGATCAGC-1 -2.694437  1.5836617  0.6631235 -2.2056429  1.78901792
AAACCGTGCTTCCG-1 10.143297  1.3685347 -1.2098237  0.7000697  2.90616465
AAACCGTGTATGCG-1  1.112813  8.1527987 -1.3323525  4.2524910 -1.96318078
```

In [149]:

```
%%R
# pca feature loadings
ann.seuscdior@reductions$pca@feature.loadings[1:5, 1:5]
```

```
                PC_1         PC_2          PC_3         PC_4          PC_5
TNFRSF4 -0.026014818  0.003254168  0.0018978898 -0.036262553  0.0167816579
CPSF3L  -0.008278225  0.009083163 -0.0007814111  0.008882667 -0.0063649304
ATAD3C  -0.003315187  0.003209684  0.0002798587 -0.001740866 -0.0003630426
C1orf86  0.010650732 -0.000268140 -0.0070081116  0.002366116 -0.0038789101
RER1     0.013711839  0.027387908 -0.0107853832  0.006192871  0.0182563197
```

In [150]:

```
%%R
# dimensional reduction results of umap
ann.seuscdior@reductions$umap@cell.embeddings[1:5, 1:2]
```

```
                    umap_1     umap_2
AAACATACAACCAC-1  7.906657  3.5560911
AAACATTGAGCTAC-1  9.248348 12.5443316
AAACATTGATCAGC-1  7.629986  3.8347855
AAACCGTGCTTCCG-1  0.131578  5.5397391
AAACCGTGTATGCG-1 10.055341 -0.6474292
```

In [151]:

```
%%R
# miscellaneous information
ann.seuscdior@reductions$pca@misc
```

```
$params
  use_highly_variable zero_center
1                TRUE        TRUE

$variance
 [1] 32.110455 18.718655 15.607329 13.235289  4.802269  3.985932  3.526233
 [8]  3.233445  3.121209  3.075261  2.998075  2.959521  2.951785  2.944248
[15]  2.913872  2.899030  2.880682  2.864685  2.843064  2.835751  2.831421
[22]  2.818236  2.803552  2.799987  2.788954  2.778101  2.770577  2.760221
[29]  2.753860  2.745955  2.737186  2.734127  2.722202  2.712311  2.702478
[36]  2.700047  2.683850  2.679051  2.676908  2.673995  2.664854  2.657311
[43]  2.651177  2.641776  2.632970  2.629520  2.624529  2.618376  2.618003
[50]  2.601866

$variance_ratio
 [1] 0.020128191 0.011733645 0.009783334 0.008296438 0.003010265 0.002498551
 [7] 0.002210392 0.002026860 0.001956505 0.001927704 0.001879320 0.001855153
[13] 0.001850304 0.001845579 0.001826538 0.001817235 0.001805733 0.001795706
[19] 0.001782152 0.001777568 0.001774855 0.001766590 0.001757385 0.001755151
[25] 0.001748234 0.001741431 0.001736715 0.001730223 0.001726236 0.001721281
[31] 0.001715784 0.001713866 0.001706391 0.001700191 0.001694027 0.001692504
[37] 0.001682351 0.001679342 0.001677999 0.001676173 0.001670443 0.001665715
[43] 0.001661870 0.001655977 0.001650457 0.001648294 0.001645166 0.001641309
[49] 0.001641075 0.001630959
```

In [152]:

```
%%R
# miscellaneous information
ann.seuscdior@reductions$umap@misc
```

```
$params
        a        b
1 0.58303 1.334167
```

In [153]:

```
%%R
# relationship of cells, graphs
# RNA_nn
ann.seuscdior@graphs$RNA_nn[1:10,1:10]
```

```
10 x 10 sparse Matrix of class "dgCMatrix"
```

```
R[write to console]:   [[ suppressing 10 column names ‘AAACATACAACCAC-1’, ‘AAACATTGAGCTAC-1’, ‘AAACATTGATCAGC-1’ ... ]]
```

```
AAACATACAACCAC-1 . . . . . . . . . .
AAACATTGAGCTAC-1 . . . . . . . . . .
AAACATTGATCAGC-1 . . . . . . . . . .
AAACCGTGCTTCCG-1 . . . . . . . . . .
AAACCGTGTATGCG-1 . . . . . . . . . .
AAACGCACTGGTAC-1 . . . . . . . . . .
AAACGCTGACCAGT-1 . . . . . . . . . .
AAACGCTGGTTCTT-1 . . . . . . . . . .
AAACGCTGTAGCCA-1 . . . . . . . . . .
AAACGCTGTTTCTG-1 . . . . . . . . . .
```

In [154]:

```
%%R
# relationship of cells, graphs
# RNA_snn
ann.seuscdior@graphs$RNA_snn[1:10,1:10]
```

```
10 x 10 sparse Matrix of class "dgCMatrix"
```

```
R[write to console]:   [[ suppressing 10 column names ‘AAACATACAACCAC-1’, ‘AAACATTGAGCTAC-1’, ‘AAACATTGATCAGC-1’ ... ]]
```

```
AAACATACAACCAC-1 . . . . . . . . . .
AAACATTGAGCTAC-1 . . . . . . . . . .
AAACATTGATCAGC-1 . . . . . . . . . .
AAACCGTGCTTCCG-1 . . . . . . . . . .
AAACCGTGTATGCG-1 . . . . . . . . . .
AAACGCACTGGTAC-1 . . . . . . . . . .
AAACGCTGACCAGT-1 . . . . . . . . . .
AAACGCTGGTTCTT-1 . . . . . . . . . .
AAACGCTGTAGCCA-1 . . . . . . . . . .
AAACGCTGTTTCTG-1 . . . . . . . . . .
```

In [155]:

```
%%R
dim(ann.seuscdior@graphs$RNA_snn)
```

```
[1] 2638 2638
```

### AnnData to SingleCellExperiemnt¶

In [156]:

```
%%R
# convert AnnData to SingleCellExperiemnt (use current conda environment)
# now integrated into GEfetch2R
AD2SCE = function(anndata.file, method = c("scDIOR", "zellkonverter", "schard"), assay = "RNA",
                  slot = "counts", use.raw = TRUE){
  # check parameters
  method <- match.arg(arg = method)

  # check file
  if(!file.exists(anndata.file)){
    stop(anndata.file, " does not exist, please check!")
  }
  # conversion
  if(method == "scDIOR"){
    sce = tryCatch(
      {
        if(use.raw){
          anndata <- reticulate::import("anndata")
          adata = anndata$read_h5ad(anndata.file)
          diopy = reticulate::import("diopy")
          h5.file = gsub(pattern = ".h5ad$", replacement = "_tmp.h5", x = anndata.file)
          diopy$output$write_h5(adata = adata$raw$to_adata(), file=h5.file, assay_name=assay, save_X = TRUE)
          dior::read_h5(file = h5.file, target.object = "singlecellexperiment")
        }else{
          dior::read_h5ad(file = anndata.file, assay_name = assay, target.object = "singlecellexperiment")
        }
      },
      error = function(cond) {
        message("There is an error when using scDIOR: ", cond)
      }
    )
  }else if(method == "zellkonverter"){
    sce = tryCatch(
      {
        anndata <- reticulate::import("anndata")
        adata <- anndata$read_h5ad(anndata.file)
        zellkonverter::AnnData2SCE(adata, X_name = slot, raw = use.raw)
      },
      error = function(cond) {
        message("There is an error when using zellkonverter: ", cond)
      }
    )
  }else if(method == "schard"){
    sce = tryCatch(
      {
        schard::h5ad2sce(anndata.file, use.raw = use.raw)
      },
      error = function(cond) {
        message("There is an error when using schard: ", cond)
      }
    )
  }
}
```

#### scDIOR¶

**Retained information** (`AnnData -> SingleCellExperiemnt`):

- count matrix:
  - `use.raw = TRUE`: raw count matrix (`raw.X -> assays`)
  - `use.raw = FALSE`: scaled count matrix (`X/layers -> assays`)
- cells' meta-information (`obs -> colData`)
- annotation of features (`var -> rowData`), missing dispersions, dispersions\_norm, highly\_variable, mean, means, std when `use.raw = TRUE`
- dimensional reduction results (`obsm -> reducedDim`)

In [157]:

```
%%R
# use.raw = TRUE: no layers, rowData missing columns, raw count matrix
# use.raw = FALSE, layers, full rowData columns, scaled count matrix
sce.scdior = AD2SCE(anndata.file = "./write/pbmc3k.h5ad",
                    method = "scDIOR", assay = "RNA", use.raw = TRUE)
sce.scdior
```

```
class: SingleCellExperiment 
dim: 13714 2638 
metadata(0):
assays(1): X
rownames(13714): AL627309.1 AP006222.2 ... PNRC2-1 SRSF10-1
rowData names(7): gene_ids n_cells ... pct_dropout_by_counts
  total_counts
colnames(2638): AAACATACAACCAC-1 AAACATTGAGCTAC-1 ... TTTGCATGAGAGGC-1
  TTTGCATGCCTCAC-1
colData names(6): n_genes n_genes_by_counts ... pct_counts_mt leiden
reducedDimNames(2): pca umap
altExpNames(0):
```

In [158]:

```
%%R
suppressMessages(library(SingleCellExperiment))
assay(sce.scdior, "X")[1:10, 1:15]
```

```
10 x 15 sparse Matrix of class "dgCMatrix"
```

```
R[write to console]:   [[ suppressing 15 column names ‘AAACATACAACCAC-1’, ‘AAACATTGAGCTAC-1’, ‘AAACATTGATCAGC-1’ ... ]]
```

```
AL627309.1    . . . . . . . . . . . . . . .
AP006222.2    . . . . . . . . . . . . . . .
RP11-206L10.2 . . . . . . . . . . . . . . .
RP11-206L10.9 . . . . . . . . . . . . . . .
LINC00115     . . . . . . . . . . . . . . .
NOC2L         . . . . . . . . . . . 1 . . .
KLHL17        . . . . . . . . . . . . . . .
PLEKHN1       . . . . . . . . . . . . . . .
RP11-54O7.17  . . . . . . . . . . . . . . .
HES4          . . . . . . . . . . . . . . .
```

In [159]:

```
%%R
# shape
dim(assay(sce.scdior, "X"))
```

```
[1] 13714  2638
```

In [160]:

```
%%R
suppressMessages(library(tidyverse))
# cells' meta-information
colData(sce.scdior) %>% head()
```

```
DataFrame with 6 rows and 6 columns
                   n_genes n_genes_by_counts total_counts total_counts_mt
                 <integer>         <integer>    <numeric>       <numeric>
AAACATACAACCAC-1       781               779         2419              73
AAACATTGAGCTAC-1      1352              1352         4903             186
AAACATTGATCAGC-1      1131              1129         3147              28
AAACCGTGCTTCCG-1       960               960         2639              46
AAACCGTGTATGCG-1       522               521          980              12
AAACGCACTGGTAC-1       782               781         2163              36
                 pct_counts_mt   leiden
                     <numeric> <factor>
AAACATACAACCAC-1      3.017776        0
AAACATTGAGCTAC-1      3.793596        2
AAACATTGATCAGC-1      0.889736        0
AAACCGTGCTTCCG-1      1.743085        4
AAACCGTGTATGCG-1      1.224490        5
AAACGCACTGGTAC-1      1.664355        0
```

In [161]:

```
%%R
# annotation of features, missing dispersions, dispersions_norm, highly_variable, mean, means, std when use.raw = TRUE
rowData(sce.scdior) %>% head()
```

```
DataFrame with 6 rows and 7 columns
                     gene_ids   n_cells        mt n_cells_by_counts mean_counts
                  <character> <integer> <logical>         <integer>   <numeric>
AL627309.1    ENSG00000237683         9     FALSE                 9  0.00333333
AP006222.2    ENSG00000228463         3     FALSE                 3  0.00111111
RP11-206L10.2 ENSG00000228327         5     FALSE                 5  0.00185185
RP11-206L10.9 ENSG00000237491         3     FALSE                 3  0.00111111
LINC00115     ENSG00000225880        18     FALSE                18  0.00666667
NOC2L         ENSG00000188976       258     FALSE               258  0.10666667
              pct_dropout_by_counts total_counts
                          <numeric>    <numeric>
AL627309.1                  99.6667            9
AP006222.2                  99.8889            3
RP11-206L10.2               99.8148            5
RP11-206L10.9               99.8889            3
LINC00115                   99.3333           18
NOC2L                       90.4444          288
```

In [162]:

```
%%R
# dimensional reduction results of pca
reducedDim(sce.scdior, "pca")[1:5, 1:5]
```

```
                     PCA_1      PCA_2      PCA_3      PCA_4       PCA_5
AAACATACAACCAC-1 -5.556221 -0.2577271  0.1867943 -2.8000970  0.05072495
AAACATTGAGCTAC-1 -7.209527 -7.4820013 -0.1627175  8.0185165 -3.00661612
AAACATTGATCAGC-1 -2.694437  1.5836617  0.6631235 -2.2056429  1.78901792
AAACCGTGCTTCCG-1 10.143297  1.3685347 -1.2098237  0.7000697  2.90616465
AAACCGTGTATGCG-1  1.112813  8.1527987 -1.3323525  4.2524910 -1.96318078
```

In [163]:

```
%%R
# dimensional reduction results of umap
reducedDim(sce.scdior, "umap")[1:5, 1:2]
```

```
                    UMAP_1     UMAP_2
AAACATACAACCAC-1  7.906657  3.5560911
AAACATTGAGCTAC-1  9.248348 12.5443316
AAACATTGATCAGC-1  7.629986  3.8347855
AAACCGTGCTTCCG-1  0.131578  5.5397391
AAACCGTGTATGCG-1 10.055341 -0.6474292
```

#### zellkonverter¶

**Retained information** (`AnnData -> SingleCellExperiemnt`):

- count matrix:
  - `use.raw = TRUE`: full raw count matrix (`raw.X -> altExp`, **13714 x 2638**); raw count matrix, log-normalized count matrix, scaled count matrix (`X and layers -> assays`, **1838 x 2638**)
  - `use.raw = FALSE`: raw count matrix, log-normalized count matrix, scaled count matrix (`X and layers -> assays`, **1838 x 2638**)
- cells' meta-information (`obs -> colData`)
- annotation of features (`var -> rowData`)
- dimensional reduction results (`obsm -> reducedDim`)
- feature loadings (`varm -> rowData`)
- unstructured annotation (`uns -> metadata`)
- relationship of cells, graphs (`obsp -> colPairs`)

In [164]:

```
%%R
sce.zell = AD2SCE(anndata.file = "./write/pbmc3k.h5ad",
                  method = "zellkonverter", slot = "scale.data", use.raw = TRUE)
sce.zell
```

```
R[write to console]: Registered S3 method overwritten by 'zellkonverter':
  method                                             from      
  py_to_r.pandas.core.arrays.categorical.Categorical reticulate
```

```
class: SingleCellExperiment 
dim: 1838 2638 
metadata(7): hvg leiden ... rank_genes_groups umap
assays(3): scale.data logcounts rawcounts
rownames(1838): TNFRSF4 CPSF3L ... S100B PRMT2
rowData names(14): gene_ids n_cells ... std varm
colnames(2638): AAACATACAACCAC-1 AAACATTGAGCTAC-1 ... TTTGCATGAGAGGC-1
  TTTGCATGCCTCAC-1
colData names(6): n_genes n_genes_by_counts ... pct_counts_mt leiden
reducedDimNames(2): X_pca X_umap
altExpNames(1): raw
```

In [165]:

```
%%R
suppressMessages(library(SingleCellExperiment))
# raw count matrix (layers: rawcounts)
assay(sce.zell, "rawcounts")[1:10, 1:15]
```

```
10 x 15 sparse Matrix of class "dgCMatrix"
```

```
R[write to console]:   [[ suppressing 15 column names ‘AAACATACAACCAC-1’, ‘AAACATTGAGCTAC-1’, ‘AAACATTGATCAGC-1’ ... ]]
```

```
TNFRSF4  . . . . . . . . 1 . . . . . .
CPSF3L   . . . . . . . . . . 1 . . . .
ATAD3C   . . . . . . . . . . . . . . .
C1orf86  . . . . . . . . 1 . . . . . .
RER1     . 1 1 1 . . . . . . . 1 . 1 .
TNFRSF25 2 . . . . . . 1 . . . . . . .
TNFRSF9  . . . . . . . . . . . . . . .
CTNNBIP1 . . . . 1 . . . . . . . 1 . .
SRM      . . . . . . 1 . . . . . . . .
UBIAD1   . . . . . . . . . . . . . . .
```

In [166]:

```
%%R
# shape
dim(assay(sce.zell, "rawcounts"))
```

```
[1] 1838 2638
```

In [167]:

```
%%R
# log-normalized count matrix (layers: logcounts)
assay(sce.zell, "logcounts")[1:10, 1:15]
```

```
10 x 15 sparse Matrix of class "dgCMatrix"
```

```
R[write to console]:   [[ suppressing 15 column names ‘AAACATACAACCAC-1’, ‘AAACATTGAGCTAC-1’, ‘AAACATTGATCAGC-1’ ... ]]
```

```
TNFRSF4  .        .        .        .        .        . .        .       
CPSF3L   .        .        .        .        .        . .        .       
ATAD3C   .        .        .        .        .        . .        .       
C1orf86  .        .        .        .        .        . .        .       
RER1     .        1.111715 1.429744 1.566387 .        . .        .       
TNFRSF25 2.226555 .        .        .        .        . .        1.690977
TNFRSF9  .        .        .        .        .        . .        .       
CTNNBIP1 .        .        .        .        2.416278 . .        .       
SRM      .        .        .        .        .        . 1.722356 .       
UBIAD1   .        .        .        .        .        . .        .       
                                                         
TNFRSF4  2.179642 . .        .        .        .        .
CPSF3L   .        . 1.268336 .        .        .        .
ATAD3C   .        . .        .        .        .        .
C1orf86  2.179642 . .        .        .        .        .
RER1     .        . .        1.646272 .        1.457932 .
TNFRSF25 .        . .        .        .        .        .
TNFRSF9  .        . .        .        .        .        .
CTNNBIP1 .        . .        .        1.638876 .        .
SRM      .        . .        .        .        .        .
UBIAD1   .        . .        .        .        .        .
```

In [168]:

```
%%R
# shape
dim(assay(sce.zell, "logcounts"))
```

```
[1] 1838 2638
```

In [169]:

```
%%R
# scaled count matrix
assay(sce.zell, "scale.data")[1:10, 1:4]
```

```
         AAACATACAACCAC-1 AAACATTGAGCTAC-1 AAACATTGATCAGC-1 AAACCGTGCTTCCG-1
TNFRSF4       -0.17146961      -0.21458235      -0.37688771      -0.28524107
CPSF3L        -0.28081229      -0.37265328      -0.29508454      -0.28173482
ATAD3C        -0.04667677      -0.05480441      -0.05752748      -0.05222671
C1orf86       -0.47516865      -0.68339121      -0.52097195      -0.48492861
RER1          -0.54402399       0.63395083       1.33264792       1.57267952
TNFRSF25       4.92849684      -0.33483663      -0.30936241      -0.27182469
TNFRSF9       -0.03802770      -0.04558870      -0.10310833      -0.07455204
CTNNBIP1      -0.28057277      -0.49826378      -0.27252606      -0.25887546
SRM           -0.34178808      -0.54191375      -0.50079864      -0.41675180
UBIAD1        -0.19536127      -0.20901665      -0.22022836      -0.20847099
```

In [170]:

```
%%R
# shape
dim(assay(sce.zell, "scale.data"))
```

```
[1] 1838 2638
```

In [171]:

```
%%R
# full raw count matrix
altExp(sce.zell)
```

```
class: SummarizedExperiment 
dim: 13714 2638 
metadata(0):
assays(1): X
rownames: NULL
rowData names(7): gene_ids n_cells ... pct_dropout_by_counts
  total_counts
colnames(2638): AAACATACAACCAC-1 AAACATTGAGCTAC-1 ... TTTGCATGAGAGGC-1
  TTTGCATGCCTCAC-1
colData names(0):
```

In [172]:

```
%%R
suppressMessages(library(tidyverse))
# cells' meta-information
colData(sce.zell) %>% head()
```

```
DataFrame with 6 rows and 6 columns
                   n_genes n_genes_by_counts total_counts total_counts_mt
                 <numeric>         <integer>    <numeric>       <numeric>
AAACATACAACCAC-1       781               779         2419              73
AAACATTGAGCTAC-1      1352              1352         4903             186
AAACATTGATCAGC-1      1131              1129         3147              28
AAACCGTGCTTCCG-1       960               960         2639              46
AAACCGTGTATGCG-1       522               521          980              12
AAACGCACTGGTAC-1       782               781         2163              36
                 pct_counts_mt   leiden
                     <numeric> <factor>
AAACATACAACCAC-1      3.017776        0
AAACATTGAGCTAC-1      3.793596        2
AAACATTGATCAGC-1      0.889736        0
AAACCGTGCTTCCG-1      1.743085        4
AAACCGTGTATGCG-1      1.224490        5
AAACGCACTGGTAC-1      1.664355        0
```

In [173]:

```
%%R
# annotation of features and feature loadings (varm)
rowData(sce.zell) %>% head()
```

```
DataFrame with 6 rows and 14 columns
                gene_ids   n_cells        mt n_cells_by_counts mean_counts
             <character> <numeric> <logical>         <numeric>   <numeric>
TNFRSF4  ENSG00000186827       155     FALSE               155  0.07740740
CPSF3L   ENSG00000127054       202     FALSE               202  0.09481481
ATAD3C   ENSG00000215915         9     FALSE                 9  0.00925926
C1orf86  ENSG00000162585       501     FALSE               501  0.22777778
RER1     ENSG00000157916       608     FALSE               608  0.29814816
TNFRSF25 ENSG00000215788       170     FALSE               170  0.08851852
         pct_dropout_by_counts total_counts highly_variable     means
                     <numeric>    <numeric>       <logical> <numeric>
TNFRSF4                94.2593          209            TRUE 0.2774103
CPSF3L                 92.5185          256            TRUE 0.3851941
ATAD3C                 99.6667           25            TRUE 0.0382519
C1orf86                81.4444          615            TRUE 0.6782825
RER1                   77.4815          805            TRUE 0.8148132
TNFRSF25               93.7037          239            TRUE 0.3026148
         dispersions dispersions_norm         mean       std
           <numeric>        <numeric>    <numeric> <numeric>
TNFRSF4      2.08605         0.665406 -3.67207e-10  0.424481
CPSF3L       4.50699         2.955005 -2.37244e-10  0.460416
ATAD3C       3.95349         4.352607  8.47299e-12  0.119465
C1orf86      2.71352         0.543183  3.38920e-10  0.685145
RER1         3.44753         1.582528  7.69630e-11  0.736050
TNFRSF25     3.19553         1.274278  3.13501e-10  0.430840
                                             varm
                                      <DataFrame>
TNFRSF4  -0.02601482: 0.00325417: 0.001897890:...
CPSF3L   -0.00827822: 0.00908316:-0.000781411:...
ATAD3C   -0.00331519: 0.00320968: 0.000279859:...
C1orf86   0.01065073:-0.00026814:-0.007008112:...
RER1      0.01371184: 0.02738791:-0.010785383:...
TNFRSF25 -0.02663716: 0.01085256: 0.002487649:...
```

In [174]:

```
%%R
# dimensional reduction results of pca
reducedDim(sce.zell, "X_pca")[1:5, 1:5]
```

```
                      [,1]       [,2]       [,3]       [,4]        [,5]
AAACATACAACCAC-1 -5.556221 -0.2577271  0.1867943 -2.8000970  0.05072495
AAACATTGAGCTAC-1 -7.209527 -7.4820013 -0.1627175  8.0185165 -3.00661612
AAACATTGATCAGC-1 -2.694437  1.5836617  0.6631235 -2.2056429  1.78901792
AAACCGTGCTTCCG-1 10.143297  1.3685347 -1.2098237  0.7000697  2.90616465
AAACCGTGTATGCG-1  1.112813  8.1527987 -1.3323525  4.2524910 -1.96318078
```

In [175]:

```
%%R
# dimensional reduction results of umap
reducedDim(sce.zell, "X_umap")[1:5, 1:2]
```

```
                      [,1]       [,2]
AAACATACAACCAC-1  7.906657  3.5560911
AAACATTGAGCTAC-1  9.248348 12.5443316
AAACATTGATCAGC-1  7.629986  3.8347855
AAACCGTGCTTCCG-1  0.131578  5.5397391
AAACCGTGTATGCG-1 10.055341 -0.6474292
```

In [176]:

```
%%R
# unstructured annotation
names(metadata(sce.zell))
```

```
[1] "hvg"               "leiden"            "log1p"            
[4] "neighbors"         "pca"               "rank_genes_groups"
[7] "umap"
```

In [177]:

```
%%R
# pca variance
metadata(sce.zell)$pca$variance
```

```
 [1] 32.110455 18.718655 15.607329 13.235289  4.802269  3.985932  3.526233
 [8]  3.233445  3.121209  3.075261  2.998075  2.959521  2.951785  2.944248
[15]  2.913872  2.899030  2.880682  2.864685  2.843064  2.835751  2.831421
[22]  2.818236  2.803552  2.799987  2.788954  2.778101  2.770577  2.760221
[29]  2.753860  2.745955  2.737186  2.734127  2.722202  2.712311  2.702478
[36]  2.700047  2.683850  2.679051  2.676908  2.673995  2.664854  2.657311
[43]  2.651177  2.641776  2.632970  2.629520  2.624529  2.618376  2.618003
[50]  2.601866
```

In [178]:

```
%%R
# relationship of cells, graphs
colPairs(sce.zell)$connectivities
```

```
SelfHits object with 41952 hits and 1 metadata column:
               from        to |         x
          <integer> <integer> | <numeric>
      [1]         1        61 |  0.114571
      [2]         1       109 |  0.303653
      [3]         1       475 |  0.124247
      [4]         1      1574 |  0.134454
      [5]         1      1981 |  0.210848
      ...       ...       ... .       ...
  [41948]      2638      1820 |  1.000000
  [41949]      2638      1879 |  0.167065
  [41950]      2638      2078 |  0.150742
  [41951]      2638      2573 |  0.314722
  [41952]      2638      2575 |  0.774504
  -------
  nnode: 2638
```

In [179]:

```
%%R
# relationship of cells, graphs
colPairs(sce.zell)$distances
```

```
SelfHits object with 23742 hits and 1 metadata column:
               from        to |         x
          <integer> <integer> | <numeric>
      [1]         1        61 |   9.75174
      [2]         1       109 |   9.46562
      [3]         1       475 |   9.72794
      [4]         1      1574 |   9.70476
      [5]         1      1981 |   9.57269
      ...       ...       ... .       ...
  [23738]      2638      1680 |   7.47523
  [23739]      2638      1879 |   8.69155
  [23740]      2638      2078 |   8.76144
  [23741]      2638      2573 |   8.26106
  [23742]      2638      2575 |   7.96147
  -------
  nnode: 2638
```

#### schard¶

**Retained information** (`AnnData -> SingleCellExperiemnt`):

- count matrix:
  - `use.raw = TRUE`: raw count matrix (`raw.X -> assays`)
  - `use.raw = FALSE`: scaled count matrix (`X -> assays`)
- cells' meta-information (`obs -> colData`)
- annotation of features (`var -> rowData`), missing dispersions, dispersions\_norm, highly\_variable, mean, means, std when `use.raw = TRUE`
- dimensional reduction results (`obsm -> reducedDim`)

In [180]:

```
%%R
sce.schard = AD2SCE(anndata.file = "./write/pbmc3k.h5ad",
                    method = "schard", use.raw = TRUE)
sce.schard
```

```
class: SingleCellExperiment 
dim: 13714 2638 
metadata(0):
assays(1): X
rownames(13714): AL627309.1 AP006222.2 ... PNRC2-1 SRSF10-1
rowData names(8): _index gene_ids ... pct_dropout_by_counts
  total_counts
colnames(2638): AAACATACAACCAC-1 AAACATTGAGCTAC-1 ... TTTGCATGAGAGGC-1
  TTTGCATGCCTCAC-1
colData names(7): _index n_genes ... pct_counts_mt leiden
reducedDimNames(2): X_pca X_umap
altExpNames(0):
```

In [181]:

```
%%R
suppressMessages(library(SingleCellExperiment))
assay(sce.schard, "X")[1:10, 1:15]
```

```
10 x 15 sparse Matrix of class "dgCMatrix"
```

```
R[write to console]:   [[ suppressing 15 column names ‘AAACATACAACCAC-1’, ‘AAACATTGAGCTAC-1’, ‘AAACATTGATCAGC-1’ ... ]]
```

```
AL627309.1    . . . . . . . . . . . . . . .
AP006222.2    . . . . . . . . . . . . . . .
RP11-206L10.2 . . . . . . . . . . . . . . .
RP11-206L10.9 . . . . . . . . . . . . . . .
LINC00115     . . . . . . . . . . . . . . .
NOC2L         . . . . . . . . . . . 1 . . .
KLHL17        . . . . . . . . . . . . . . .
PLEKHN1       . . . . . . . . . . . . . . .
RP11-54O7.17  . . . . . . . . . . . . . . .
HES4          . . . . . . . . . . . . . . .
```

In [182]:

```
%%R
# shape
dim(assay(sce.schard, "X"))
```

```
[1] 13714  2638
```

In [183]:

```
%%R
suppressMessages(library(tidyverse))
# cells' meta-information
colData(sce.schard) %>% head()
```

```
DataFrame with 6 rows and 7 columns
                           _index   n_genes n_genes_by_counts total_counts
                      <character> <integer>         <integer>    <numeric>
AAACATACAACCAC-1 AAACATACAACCAC-1       781               779         2419
AAACATTGAGCTAC-1 AAACATTGAGCTAC-1      1352              1352         4903
AAACATTGATCAGC-1 AAACATTGATCAGC-1      1131              1129         3147
AAACCGTGCTTCCG-1 AAACCGTGCTTCCG-1       960               960         2639
AAACCGTGTATGCG-1 AAACCGTGTATGCG-1       522               521          980
AAACGCACTGGTAC-1 AAACGCACTGGTAC-1       782               781         2163
                 total_counts_mt pct_counts_mt      leiden
                       <numeric>     <numeric> <character>
AAACATACAACCAC-1              73      3.017776           0
AAACATTGAGCTAC-1             186      3.793596           2
AAACATTGATCAGC-1              28      0.889736           0
AAACCGTGCTTCCG-1              46      1.743085           4
AAACCGTGTATGCG-1              12      1.224490           5
AAACGCACTGGTAC-1              36      1.664355           0
```

In [184]:

```
%%R
# annotation of features, missing dispersions, dispersions_norm, highly_variable, mean, means, std when use.raw = TRUE
rowData(sce.schard) %>% head()
```

```
DataFrame with 6 rows and 8 columns
                     _index        gene_ids   n_cells          mt
                <character>     <character> <integer> <character>
AL627309.1       AL627309.1 ENSG00000237683         9       FALSE
AP006222.2       AP006222.2 ENSG00000228463         3       FALSE
RP11-206L10.2 RP11-206L10.2 ENSG00000228327         5       FALSE
RP11-206L10.9 RP11-206L10.9 ENSG00000237491         3       FALSE
LINC00115         LINC00115 ENSG00000225880        18       FALSE
NOC2L                 NOC2L ENSG00000188976       258       FALSE
              n_cells_by_counts mean_counts pct_dropout_by_counts total_counts
                      <integer>   <numeric>             <numeric>    <numeric>
AL627309.1                    9  0.00333333               99.6667            9
AP006222.2                    3  0.00111111               99.8889            3
RP11-206L10.2                 5  0.00185185               99.8148            5
RP11-206L10.9                 3  0.00111111               99.8889            3
LINC00115                    18  0.00666667               99.3333           18
NOC2L                       258  0.10666667               90.4444          288
```

In [185]:

```
%%R
# dimensional reduction results of pca
reducedDim(sce.schard, "X_pca")[1:5, 1:5]
```

```
                      [,1]       [,2]       [,3]       [,4]        [,5]
AAACATACAACCAC-1 -5.556221 -0.2577271  0.1867943 -2.8000970  0.05072495
AAACATTGAGCTAC-1 -7.209527 -7.4820013 -0.1627175  8.0185165 -3.00661612
AAACATTGATCAGC-1 -2.694437  1.5836617  0.6631235 -2.2056429  1.78901792
AAACCGTGCTTCCG-1 10.143297  1.3685347 -1.2098237  0.7000697  2.90616465
AAACCGTGTATGCG-1  1.112813  8.1527987 -1.3323525  4.2524910 -1.96318078
```

In [186]:

```
%%R
# dimensional reduction results of umap
reducedDim(sce.schard, "X_umap")[1:5, 1:2]
```

```
                      [,1]       [,2]
AAACATACAACCAC-1  7.906657  3.5560911
AAACATTGAGCTAC-1  9.248348 12.5443316
AAACATTGATCAGC-1  7.629986  3.8347855
AAACCGTGCTTCCG-1  0.131578  5.5397391
AAACCGTGTATGCG-1 10.055341 -0.6474292
```

### SingleCellExperiemnt to AnnData¶

#### SingleCellExperiemnt¶

In [187]:

```
%%R
# test data (AD2SCE/zellkonverter)
pbmc3k.sce = sce.zell
pbmc3k.sce
```

```
class: SingleCellExperiment 
dim: 1838 2638 
metadata(7): hvg leiden ... rank_genes_groups umap
assays(3): scale.data logcounts rawcounts
rownames(1838): TNFRSF4 CPSF3L ... S100B PRMT2
rowData names(14): gene_ids n_cells ... std varm
colnames(2638): AAACATACAACCAC-1 AAACATTGAGCTAC-1 ... TTTGCATGAGAGGC-1
  TTTGCATGCCTCAC-1
colData names(6): n_genes n_genes_by_counts ... pct_counts_mt leiden
reducedDimNames(2): X_pca X_umap
altExpNames(1): raw
```

##### Count matrix (`assays` and `altExp`)¶

In [188]:

```
%%R
# raw count matrix
assay(pbmc3k.sce, "rawcounts")[1:10,1:15]
```

```
10 x 15 sparse Matrix of class "dgCMatrix"
```

```
R[write to console]:   [[ suppressing 15 column names ‘AAACATACAACCAC-1’, ‘AAACATTGAGCTAC-1’, ‘AAACATTGATCAGC-1’ ... ]]
```

```
TNFRSF4  . . . . . . . . 1 . . . . . .
CPSF3L   . . . . . . . . . . 1 . . . .
ATAD3C   . . . . . . . . . . . . . . .
C1orf86  . . . . . . . . 1 . . . . . .
RER1     . 1 1 1 . . . . . . . 1 . 1 .
TNFRSF25 2 . . . . . . 1 . . . . . . .
TNFRSF9  . . . . . . . . . . . . . . .
CTNNBIP1 . . . . 1 . . . . . . . 1 . .
SRM      . . . . . . 1 . . . . . . . .
UBIAD1   . . . . . . . . . . . . . . .
```

In [189]:

```
%%R
# log-normalized count matrix
assay(pbmc3k.sce, "logcounts")[1:10,1:15]
```

```
10 x 15 sparse Matrix of class "dgCMatrix"
```

```
R[write to console]:   [[ suppressing 15 column names ‘AAACATACAACCAC-1’, ‘AAACATTGAGCTAC-1’, ‘AAACATTGATCAGC-1’ ... ]]
```

```
TNFRSF4  .        .        .        .        .        . .        .       
CPSF3L   .        .        .        .        .        . .        .       
ATAD3C   .        .        .        .        .        . .        .       
C1orf86  .        .        .        .        .        . .        .       
RER1     .        1.111715 1.429744 1.566387 .        . .        .       
TNFRSF25 2.226555 .        .        .        .        . .        1.690977
TNFRSF9  .        .        .        .        .        . .        .       
CTNNBIP1 .        .        .        .        2.416278 . .        .       
SRM      .        .        .        .        .        . 1.722356 .       
UBIAD1   .        .        .        .        .        . .        .       
                                                         
TNFRSF4  2.179642 . .        .        .        .        .
CPSF3L   .        . 1.268336 .        .        .        .
ATAD3C   .        . .        .        .        .        .
C1orf86  2.179642 . .        .        .        .        .
RER1     .        . .        1.646272 .        1.457932 .
TNFRSF25 .        . .        .        .        .        .
TNFRSF9  .        . .        .        .        .        .
CTNNBIP1 .        . .        .        1.638876 .        .
SRM      .        . .        .        .        .        .
UBIAD1   .        . .        .        .        .        .
```

In [190]:

```
%%R
# scaled count matrix
assay(pbmc3k.sce, "scale.data")[1:10,1:4]
```

```
         AAACATACAACCAC-1 AAACATTGAGCTAC-1 AAACATTGATCAGC-1 AAACCGTGCTTCCG-1
TNFRSF4       -0.17146961      -0.21458235      -0.37688771      -0.28524107
CPSF3L        -0.28081229      -0.37265328      -0.29508454      -0.28173482
ATAD3C        -0.04667677      -0.05480441      -0.05752748      -0.05222671
C1orf86       -0.47516865      -0.68339121      -0.52097195      -0.48492861
RER1          -0.54402399       0.63395083       1.33264792       1.57267952
TNFRSF25       4.92849684      -0.33483663      -0.30936241      -0.27182469
TNFRSF9       -0.03802770      -0.04558870      -0.10310833      -0.07455204
CTNNBIP1      -0.28057277      -0.49826378      -0.27252606      -0.25887546
SRM           -0.34178808      -0.54191375      -0.50079864      -0.41675180
UBIAD1        -0.19536127      -0.20901665      -0.22022836      -0.20847099
```

In [191]:

```
%%R
# full raw count matrix
altExp(pbmc3k.sce)
```

```
class: SummarizedExperiment 
dim: 13714 2638 
metadata(0):
assays(1): X
rownames: NULL
rowData names(7): gene_ids n_cells ... pct_dropout_by_counts
  total_counts
colnames(2638): AAACATACAACCAC-1 AAACATTGAGCTAC-1 ... TTTGCATGAGAGGC-1
  TTTGCATGCCTCAC-1
colData names(0):
```

In [192]:

```
%%R
# full raw count matrix
assay(altExp(pbmc3k.sce))[1:10,1:15]
```

```
10 x 15 sparse Matrix of class "dgCMatrix"
```

```
R[write to console]:   [[ suppressing 15 column names ‘AAACATACAACCAC-1’, ‘AAACATTGAGCTAC-1’, ‘AAACATTGATCAGC-1’ ... ]]
```

```
 [1,] . . . . . . . . . . . . . . .
 [2,] . . . . . . . . . . . . . . .
 [3,] . . . . . . . . . . . . . . .
 [4,] . . . . . . . . . . . . . . .
 [5,] . . . . . . . . . . . . . . .
 [6,] . . . . . . . . . . . 1 . . .
 [7,] . . . . . . . . . . . . . . .
 [8,] . . . . . . . . . . . . . . .
 [9,] . . . . . . . . . . . . . . .
[10,] . . . . . . . . . . . . . . .
```

##### `colData` - cells' meta-information¶

In [193]:

```
%%R
colData(pbmc3k.sce) %>% head()
```

```
DataFrame with 6 rows and 6 columns
                   n_genes n_genes_by_counts total_counts total_counts_mt
                 <numeric>         <integer>    <numeric>       <numeric>
AAACATACAACCAC-1       781               779         2419              73
AAACATTGAGCTAC-1      1352              1352         4903             186
AAACATTGATCAGC-1      1131              1129         3147              28
AAACCGTGCTTCCG-1       960               960         2639              46
AAACCGTGTATGCG-1       522               521          980              12
AAACGCACTGGTAC-1       782               781         2163              36
                 pct_counts_mt   leiden
                     <numeric> <factor>
AAACATACAACCAC-1      3.017776        0
AAACATTGAGCTAC-1      3.793596        2
AAACATTGATCAGC-1      0.889736        0
AAACCGTGCTTCCG-1      1.743085        4
AAACCGTGTATGCG-1      1.224490        5
AAACGCACTGGTAC-1      1.664355        0
```

##### `rowData` - annotation of features and feature loadings¶

In [194]:

```
%%R
# feature loadings in column varm
rowData(pbmc3k.sce) %>% head()
```

```
DataFrame with 6 rows and 14 columns
                gene_ids   n_cells        mt n_cells_by_counts mean_counts
             <character> <numeric> <logical>         <numeric>   <numeric>
TNFRSF4  ENSG00000186827       155     FALSE               155  0.07740740
CPSF3L   ENSG00000127054       202     FALSE               202  0.09481481
ATAD3C   ENSG00000215915         9     FALSE                 9  0.00925926
C1orf86  ENSG00000162585       501     FALSE               501  0.22777778
RER1     ENSG00000157916       608     FALSE               608  0.29814816
TNFRSF25 ENSG00000215788       170     FALSE               170  0.08851852
         pct_dropout_by_counts total_counts highly_variable     means
                     <numeric>    <numeric>       <logical> <numeric>
TNFRSF4                94.2593          209            TRUE 0.2774103
CPSF3L                 92.5185          256            TRUE 0.3851941
ATAD3C                 99.6667           25            TRUE 0.0382519
C1orf86                81.4444          615            TRUE 0.6782825
RER1                   77.4815          805            TRUE 0.8148132
TNFRSF25               93.7037          239            TRUE 0.3026148
         dispersions dispersions_norm         mean       std
           <numeric>        <numeric>    <numeric> <numeric>
TNFRSF4      2.08605         0.665406 -3.67207e-10  0.424481
CPSF3L       4.50699         2.955005 -2.37244e-10  0.460416
ATAD3C       3.95349         4.352607  8.47299e-12  0.119465
C1orf86      2.71352         0.543183  3.38920e-10  0.685145
RER1         3.44753         1.582528  7.69630e-11  0.736050
TNFRSF25     3.19553         1.274278  3.13501e-10  0.430840
                                             varm
                                      <DataFrame>
TNFRSF4  -0.02601482: 0.00325417: 0.001897890:...
CPSF3L   -0.00827822: 0.00908316:-0.000781411:...
ATAD3C   -0.00331519: 0.00320968: 0.000279859:...
C1orf86   0.01065073:-0.00026814:-0.007008112:...
RER1      0.01371184: 0.02738791:-0.010785383:...
TNFRSF25 -0.02663716: 0.01085256: 0.002487649:...
```

##### `reducedDim` - dimensional reduction results¶

In [195]:

```
%%R
reducedDim(pbmc3k.sce, "X_pca")[1:5, 1:5]
```

```
                      [,1]       [,2]       [,3]       [,4]        [,5]
AAACATACAACCAC-1 -5.556221 -0.2577271  0.1867943 -2.8000970  0.05072495
AAACATTGAGCTAC-1 -7.209527 -7.4820013 -0.1627175  8.0185165 -3.00661612
AAACATTGATCAGC-1 -2.694437  1.5836617  0.6631235 -2.2056429  1.78901792
AAACCGTGCTTCCG-1 10.143297  1.3685347 -1.2098237  0.7000697  2.90616465
AAACCGTGTATGCG-1  1.112813  8.1527987 -1.3323525  4.2524910 -1.96318078
```

##### `colPairs` - relationship of cells, graphs¶

In [196]:

```
%%R
colPairs(pbmc3k.sce)
```

```
List of length 2
names(2): connectivities distances
```

In [197]:

```
%%R
colPairs(pbmc3k.sce)$connectivities
```

```
SelfHits object with 41952 hits and 1 metadata column:
               from        to |         x
          <integer> <integer> | <numeric>
      [1]         1        61 |  0.114571
      [2]         1       109 |  0.303653
      [3]         1       475 |  0.124247
      [4]         1      1574 |  0.134454
      [5]         1      1981 |  0.210848
      ...       ...       ... .       ...
  [41948]      2638      1820 |  1.000000
  [41949]      2638      1879 |  0.167065
  [41950]      2638      2078 |  0.150742
  [41951]      2638      2573 |  0.314722
  [41952]      2638      2575 |  0.774504
  -------
  nnode: 2638
```

In [198]:

```
%%R
colPairs(pbmc3k.sce)$distances
```

```
SelfHits object with 23742 hits and 1 metadata column:
               from        to |         x
          <integer> <integer> | <numeric>
      [1]         1        61 |   9.75174
      [2]         1       109 |   9.46562
      [3]         1       475 |   9.72794
      [4]         1      1574 |   9.70476
      [5]         1      1981 |   9.57269
      ...       ...       ... .       ...
  [23738]      2638      1680 |   7.47523
  [23739]      2638      1879 |   8.69155
  [23740]      2638      2078 |   8.76144
  [23741]      2638      2573 |   8.26106
  [23742]      2638      2575 |   7.96147
  -------
  nnode: 2638
```

##### `metadata` - unstructured annotation¶

In [199]:

```
%%R
# all annotation
names(metadata(pbmc3k.sce))
```

```
[1] "hvg"               "leiden"            "log1p"            
[4] "neighbors"         "pca"               "rank_genes_groups"
[7] "umap"
```

In [200]:

```
%%R
# pca variance
metadata(pbmc3k.sce)$pca$variance
```

```
 [1] 32.110455 18.718655 15.607329 13.235289  4.802269  3.985932  3.526233
 [8]  3.233445  3.121209  3.075261  2.998075  2.959521  2.951785  2.944248
[15]  2.913872  2.899030  2.880682  2.864685  2.843064  2.835751  2.831421
[22]  2.818236  2.803552  2.799987  2.788954  2.778101  2.770577  2.760221
[29]  2.753860  2.745955  2.737186  2.734127  2.722202  2.712311  2.702478
[36]  2.700047  2.683850  2.679051  2.676908  2.673995  2.664854  2.657311
[43]  2.651177  2.641776  2.632970  2.629520  2.624529  2.618376  2.618003
[50]  2.601866
```

In [201]:

```
%%R
# convert SingleCellExperiemnt to AnnData (use current conda environment)
# now integrated into GEfetch2R
SCE2AD = function(sce.obj, method = c("sceasy",	"scDIOR", "zellkonverter"), out.folder = NULL,
                  out.filename = NULL, slot = "counts"){
  # check parameters
  method <- match.arg(arg = method)
  # check folder
  if(is.null(out.folder)){
    out.folder = getwd()
  }
  if(! dir.exists(out.folder)){
    message(out.folder, " does not exist, create automatically!")
    dir.create(path = out.folder, showWarnings = FALSE)
  }
  # out name
  out.name = deparse(substitute(sce.obj))
  # conversion
  if(method == "sceasy"){
    if(is.null(out.filename)){
      sceasy.out.name = file.path(out.folder, paste0(out.name, "_sceasy.h5ad"))
    }else{
      sceasy.out.name = file.path(out.folder, out.filename)
    }
    sceasy.log = tryCatch(
      {
        # reticulate::use_condaenv("/Applications/anaconda3", required = TRUE)
        # or set RETICULATE_PYTHON = "/Applications/anaconda3/bin/python" in Renvion
        sceasy::convertFormat(sce.obj, from="sce", to="anndata", drop_single_values = FALSE,
                              outFile=sceasy.out.name, main_layer = slot)
      },
      error = function(cond) {
        message("There is an error when using sceasy: ", cond)
      }
    )
    return(sceasy.log)
  }else if(method == "scDIOR"){
    if(is.null(out.filename)){
      scdior.out.name = file.path(out.folder, paste0(out.name, "_scDIOR.h5"))
    }else{
      scdior.out.name = file.path(out.folder, out.filename)
    }
    scdior.log = tryCatch(
      {
        dior::write_h5(data = sce.obj, object.type = "singlecellexperiment",
                       file = scdior.out.name)
        # adata = diopy.input.read_h5(file = 'pbmc3k.sce.h5') # require diopy to load h5 to AnnData
      },
      error = function(cond) {
        message("There is an error when using scDIOR: ", cond)
      }
    )
    return(scdior.log)
  }else if(method == "zellkonverter"){
    if(is.null(out.filename)){
      zell.out.name = file.path(out.folder, paste0(out.name, "_zellkonverter.h5ad"))
    }else{
      zell.out.name = file.path(out.folder, out.filename)
    }
    zell.log = tryCatch(
      {
        anndata <- reticulate::import("anndata")
        adata <- zellkonverter::SCE2AnnData(sce.obj, X_name = slot)
        adata$write_h5ad(zell.out.name)
      },
      error = function(cond) {
        message("There is an error when using zellkonverter: ", cond)
      }
    )
    return(zell.log)
  }
}
```

#### sceasy¶

**Retained information** (`SingleCellExperiemnt -> AnnData`):

- count matrix: log-normalized data/scaled data/raw count matrix (`logcounts/scale.data/rawcounts -> X`) (`slot` parameter)
- cells' meta-information (`colData -> obs`)
- annotation of features and feature loadings (`rowData -> var`)
- dimensional reduction results (`reducedDim -> obsm`)

In [202]:

```
%%R
# use raw count matrix
SCE2AD(sce.obj = pbmc3k.sce, method = "sceasy", 
       out.folder = "./", slot = "rawcounts")
```

```
/Applications/anaconda3/lib/python3.7/site-packages/rpy2/rinterface.py:807: FutureWarning: X.dtype being converted to np.float32 from float64. In the next version of anndata (0.9) conversion will not be automatic. Pass dtype explicitly to avoid this warning. Pass `AnnData(X, dtype=X.dtype, ...)` to get the future behavour.
  error_occured)
```

```
AnnData object with n_obs × n_vars = 2638 × 1838
    obs: 'n_genes', 'n_genes_by_counts', 'total_counts', 'total_counts_mt', 'pct_counts_mt', 'leiden'
    var: 'gene_ids', 'n_cells', 'mt', 'n_cells_by_counts', 'mean_counts', 'pct_dropout_by_counts', 'total_counts', 'highly_variable', 'means', 'dispersions', 'dispersions_norm', 'mean', 'std', 'varm.PCs.V1', 'varm.PCs.V2', 'varm.PCs.V3', 'varm.PCs.V4', 'varm.PCs.V5', 'varm.PCs.V6', 'varm.PCs.V7', 'varm.PCs.V8', 'varm.PCs.V9', 'varm.PCs.V10', 'varm.PCs.V11', 'varm.PCs.V12', 'varm.PCs.V13', 'varm.PCs.V14', 'varm.PCs.V15', 'varm.PCs.V16', 'varm.PCs.V17', 'varm.PCs.V18', 'varm.PCs.V19', 'varm.PCs.V20', 'varm.PCs.V21', 'varm.PCs.V22', 'varm.PCs.V23', 'varm.PCs.V24', 'varm.PCs.V25', 'varm.PCs.V26', 'varm.PCs.V27', 'varm.PCs.V28', 'varm.PCs.V29', 'varm.PCs.V30', 'varm.PCs.V31', 'varm.PCs.V32', 'varm.PCs.V33', 'varm.PCs.V34', 'varm.PCs.V35', 'varm.PCs.V36', 'varm.PCs.V37', 'varm.PCs.V38', 'varm.PCs.V39', 'varm.PCs.V40', 'varm.PCs.V41', 'varm.PCs.V42', 'varm.PCs.V43', 'varm.PCs.V44', 'varm.PCs.V45', 'varm.PCs.V46', 'varm.PCs.V47', 'varm.PCs.V48', 'varm.PCs.V49', 'varm.PCs.V50'
    obsm: 'X_x_pca', 'X_x_umap'
```

In [203]:

```
scesceasy_ann = sc.read("./pbmc3k.sce_sceasy.h5ad")
scesceasy_ann
```

Out[203]:

```
AnnData object with n_obs × n_vars = 2638 × 1838
    obs: 'n_genes', 'n_genes_by_counts', 'total_counts', 'total_counts_mt', 'pct_counts_mt', 'leiden'
    var: 'gene_ids', 'n_cells', 'mt', 'n_cells_by_counts', 'mean_counts', 'pct_dropout_by_counts', 'total_counts', 'highly_variable', 'means', 'dispersions', 'dispersions_norm', 'mean', 'std', 'varm.PCs.V1', 'varm.PCs.V2', 'varm.PCs.V3', 'varm.PCs.V4', 'varm.PCs.V5', 'varm.PCs.V6', 'varm.PCs.V7', 'varm.PCs.V8', 'varm.PCs.V9', 'varm.PCs.V10', 'varm.PCs.V11', 'varm.PCs.V12', 'varm.PCs.V13', 'varm.PCs.V14', 'varm.PCs.V15', 'varm.PCs.V16', 'varm.PCs.V17', 'varm.PCs.V18', 'varm.PCs.V19', 'varm.PCs.V20', 'varm.PCs.V21', 'varm.PCs.V22', 'varm.PCs.V23', 'varm.PCs.V24', 'varm.PCs.V25', 'varm.PCs.V26', 'varm.PCs.V27', 'varm.PCs.V28', 'varm.PCs.V29', 'varm.PCs.V30', 'varm.PCs.V31', 'varm.PCs.V32', 'varm.PCs.V33', 'varm.PCs.V34', 'varm.PCs.V35', 'varm.PCs.V36', 'varm.PCs.V37', 'varm.PCs.V38', 'varm.PCs.V39', 'varm.PCs.V40', 'varm.PCs.V41', 'varm.PCs.V42', 'varm.PCs.V43', 'varm.PCs.V44', 'varm.PCs.V45', 'varm.PCs.V46', 'varm.PCs.V47', 'varm.PCs.V48', 'varm.PCs.V49', 'varm.PCs.V50'
    obsm: 'X_x_pca', 'X_x_umap'
```

In [204]:

```
# raw count matrix
scesceasy_ann.X.toarray()
# shape
scesceasy_ann.X.shape
```

Out[204]:

```
array([[0., 0., 0., ..., 0., 0., 0.],
       [0., 0., 0., ..., 0., 0., 0.],
       [0., 0., 0., ..., 0., 0., 1.],
       ...,
       [0., 0., 0., ..., 0., 0., 1.],
       [0., 0., 0., ..., 0., 0., 0.],
       [0., 0., 0., ..., 0., 0., 0.]], dtype=float32)
```

Out[204]:

```
(2638, 1838)
```

In [205]:

```
# cells' meta-information
scesceasy_ann.obs.head()
```

Out[205]:

|  | n\_genes | n\_genes\_by\_counts | total\_counts | total\_counts\_mt | pct\_counts\_mt | leiden |
| --- | --- | --- | --- | --- | --- | --- |
| AAACATACAACCAC-1 | 781.0 | 779 | 2419.0 | 73.0 | 3.017776 | 0 |
| AAACATTGAGCTAC-1 | 1352.0 | 1352 | 4903.0 | 186.0 | 3.793596 | 2 |
| AAACATTGATCAGC-1 | 1131.0 | 1129 | 3147.0 | 28.0 | 0.889736 | 0 |
| AAACCGTGCTTCCG-1 | 960.0 | 960 | 2639.0 | 46.0 | 1.743085 | 4 |
| AAACCGTGTATGCG-1 | 522.0 | 521 | 980.0 | 12.0 | 1.224490 | 5 |

In [206]:

```
# annotation of features and cell loadings (varm.PCs.V41...)
scesceasy_ann.var.head()
# shape
scesceasy_ann.var.shape
```

Out[206]:

|  | gene\_ids | n\_cells | mt | n\_cells\_by\_counts | mean\_counts | pct\_dropout\_by\_counts | total\_counts | highly\_variable | means | dispersions | ... | varm.PCs.V41 | varm.PCs.V42 | varm.PCs.V43 | varm.PCs.V44 | varm.PCs.V45 | varm.PCs.V46 | varm.PCs.V47 | varm.PCs.V48 | varm.PCs.V49 | varm.PCs.V50 |
| --- | --- | --- | --- | --- | --- | --- | --- | --- | --- | --- | --- | --- | --- | --- | --- | --- | --- | --- | --- | --- | --- |
| TNFRSF4 | ENSG00000186827 | 155.0 | False | 155.0 | 0.077407 | 94.259259 | 209.0 | True | 0.277410 | 2.086050 | ... | 0.002325 | -0.040325 | 0.038295 | 0.023172 | 0.045645 | 0.043366 | 0.004316 | -0.005188 | 0.014497 | -0.000667 |
| CPSF3L | ENSG00000127054 | 202.0 | False | 202.0 | 0.094815 | 92.518519 | 256.0 | True | 0.385194 | 4.506987 | ... | 0.041707 | -0.021910 | 0.017430 | 0.002010 | -0.014974 | 0.021934 | 0.002236 | 0.030873 | -0.008870 | -0.002881 |
| ATAD3C | ENSG00000215915 | 9.0 | False | 9.0 | 0.009259 | 99.666667 | 25.0 | True | 0.038252 | 3.953486 | ... | -0.001872 | 0.006610 | -0.003863 | -0.005954 | 0.000394 | 0.010592 | -0.009244 | 0.010148 | -0.000530 | 0.001508 |
| C1orf86 | ENSG00000162585 | 501.0 | False | 501.0 | 0.227778 | 81.444444 | 615.0 | True | 0.678283 | 2.713522 | ... | 0.001046 | 0.001486 | -0.019350 | 0.021935 | -0.028314 | 0.050388 | -0.021616 | 0.028711 | -0.018219 | -0.027101 |
| RER1 | ENSG00000157916 | 608.0 | False | 608.0 | 0.298148 | 77.481481 | 805.0 | True | 0.814813 | 3.447533 | ... | 0.003501 | 0.022283 | 0.002127 | 0.015502 | -0.026489 | 0.007630 | -0.008917 | 0.004199 | -0.017951 | 0.024898 |

5 rows × 63 columns

Out[206]:

```
(1838, 63)
```

In [207]:

```
# dimensional reduction results of pca
scesceasy_ann.obsm['X_x_pca'][0:5, 0:5]
# dimensional reduction results of umap
scesceasy_ann.obsm['X_x_umap'][0:5, 0:2]
```

Out[207]:

```
array([[-5.55622101, -0.25772715,  0.18679433, -2.80009699,  0.05072495],
       [-7.20952702, -7.4820013 , -0.16271746,  8.01851654, -3.00661612],
       [-2.69443727,  1.58366168,  0.66312349, -2.20564294,  1.78901792],
       [10.1432972 ,  1.36853468, -1.20982373,  0.70006967,  2.90616465],
       [ 1.11281347,  8.15279865, -1.33235252,  4.252491  , -1.96318078]])
```

Out[207]:

```
array([[ 7.90665722,  3.55609107],
       [ 9.24834824, 12.54433155],
       [ 7.62998581,  3.83478546],
       [ 0.13157795,  5.53973913],
       [10.05534077, -0.64742917]])
```

#### zellkonverter¶

**Retained information** (`SingleCellExperiemnt -> AnnData`):

- count matrix (`slot = "rawcounts"`): raw count matrix (`rawcounts -> X`), log-normalized data (`logcounts -> layers['logcounts']`), scaled count matrix (`scale.data -> layers['scale.data']`)
- cells' meta-information (`colData -> obs`)
- annotation of features (`rowData -> var`)
- dimensional reduction results (`reducedDim -> obsm`)
- relationship of cells, graphs (`colPairs -> obsp`)
- unstructured annotation (`metadata -> uns`)
- feature loadings (`rowData -> varm`)

In [208]:

```
%%R
# save rawcounts to X
SCE2AD(sce.obj = pbmc3k.sce, method = "zellkonverter", 
       out.folder = "./", slot = "rawcounts")
```

```
NULL
```

In [209]:

```
scezell_ann = sc.read("./pbmc3k.sce_zellkonverter.h5ad")
scezell_ann
```

Out[209]:

```
AnnData object with n_obs × n_vars = 2638 × 1838
    obs: 'n_genes', 'n_genes_by_counts', 'total_counts', 'total_counts_mt', 'pct_counts_mt', 'leiden'
    var: 'gene_ids', 'n_cells', 'mt', 'n_cells_by_counts', 'mean_counts', 'pct_dropout_by_counts', 'total_counts', 'highly_variable', 'means', 'dispersions', 'dispersions_norm', 'mean', 'std'
    uns: 'X_name', 'hvg', 'leiden', 'log1p', 'neighbors', 'pca', 'rank_genes_groups', 'umap'
    obsm: 'X_pca', 'X_umap'
    varm: 'PCs'
    layers: 'logcounts', 'scale.data'
    obsp: 'connectivities', 'distances'
```

In [210]:

```
# raw count matrix
scezell_ann.X.toarray()
```

Out[210]:

```
array([[0., 0., 0., ..., 0., 0., 0.],
       [0., 0., 0., ..., 0., 0., 0.],
       [0., 0., 0., ..., 0., 0., 1.],
       ...,
       [0., 0., 0., ..., 0., 0., 1.],
       [0., 0., 0., ..., 0., 0., 0.],
       [0., 0., 0., ..., 0., 0., 0.]])
```

In [211]:

```
# log-normalized count matrix
scezell_ann.layers['logcounts'].toarray()
```

Out[211]:

```
array([[0.        , 0.        , 0.        , ..., 0.        , 0.        ,
        0.        ],
       [0.        , 0.        , 0.        , ..., 0.        , 0.        ,
        0.        ],
       [0.        , 0.        , 0.        , ..., 0.        , 0.        ,
        1.42974401],
       ...,
       [0.        , 0.        , 0.        , ..., 0.        , 0.        ,
        1.93704844],
       [0.        , 0.        , 0.        , ..., 0.        , 0.        ,
        0.        ],
       [0.        , 0.        , 0.        , ..., 0.        , 0.        ,
        0.        ]])
```

In [212]:

```
# scaled count matrix
scezell_ann.layers['scale.data']
```

Out[212]:

```
array([[-0.17146961, -0.28081229, -0.04667677, ..., -0.09826882,
        -0.20909512, -0.53120333],
       [-0.21458235, -0.37265328, -0.05480441, ..., -0.266844  ,
        -0.31314582, -0.5966543 ],
       [-0.37688771, -0.29508454, -0.05752748, ..., -0.15865591,
        -0.17087644,  1.37899971],
       ...,
       [-0.20708963, -0.2504642 , -0.04639699, ..., -0.05114426,
        -0.16106427,  2.04149699],
       [-0.19032849, -0.2263338 , -0.04399936, ..., -0.00591774,
        -0.13521305, -0.48211104],
       [-0.33378935, -0.25358772, -0.05271561, ..., -0.07842438,
        -0.13032718, -0.47133783]])
```

In [213]:

```
# cells' meta-information
scezell_ann.obs.head()
```

Out[213]:

|  | n\_genes | n\_genes\_by\_counts | total\_counts | total\_counts\_mt | pct\_counts\_mt | leiden |
| --- | --- | --- | --- | --- | --- | --- |
| AAACATACAACCAC-1 | 781.0 | 779 | 2419.0 | 73.0 | 3.017776 | 0 |
| AAACATTGAGCTAC-1 | 1352.0 | 1352 | 4903.0 | 186.0 | 3.793596 | 2 |
| AAACATTGATCAGC-1 | 1131.0 | 1129 | 3147.0 | 28.0 | 0.889736 | 0 |
| AAACCGTGCTTCCG-1 | 960.0 | 960 | 2639.0 | 46.0 | 1.743085 | 4 |
| AAACCGTGTATGCG-1 | 522.0 | 521 | 980.0 | 12.0 | 1.224490 | 5 |

In [214]:

```
# annotation of features
scezell_ann.var.head()
# shape
scezell_ann.var.shape
```

Out[214]:

|  | gene\_ids | n\_cells | mt | n\_cells\_by\_counts | mean\_counts | pct\_dropout\_by\_counts | total\_counts | highly\_variable | means | dispersions | dispersions\_norm | mean | std |
| --- | --- | --- | --- | --- | --- | --- | --- | --- | --- | --- | --- | --- | --- |
| TNFRSF4 | ENSG00000186827 | 155.0 | False | 155.0 | 0.077407 | 94.259259 | 209.0 | True | 0.277410 | 2.086050 | 0.665406 | -3.672069e-10 | 0.424481 |
| CPSF3L | ENSG00000127054 | 202.0 | False | 202.0 | 0.094815 | 92.518519 | 256.0 | True | 0.385194 | 4.506987 | 2.955005 | -2.372437e-10 | 0.460416 |
| ATAD3C | ENSG00000215915 | 9.0 | False | 9.0 | 0.009259 | 99.666667 | 25.0 | True | 0.038252 | 3.953486 | 4.352607 | 8.472988e-12 | 0.119465 |
| C1orf86 | ENSG00000162585 | 501.0 | False | 501.0 | 0.227778 | 81.444444 | 615.0 | True | 0.678283 | 2.713522 | 0.543183 | 3.389195e-10 | 0.685145 |
| RER1 | ENSG00000157916 | 608.0 | False | 608.0 | 0.298148 | 77.481481 | 805.0 | True | 0.814813 | 3.447533 | 1.582528 | 7.696297e-11 | 0.736050 |

Out[214]:

```
(1838, 13)
```

In [215]:

```
# dimensional reduction results of pca
scezell_ann.obsm['X_pca'][0:5, 0:5]
# dimensional reduction results of umap
scezell_ann.obsm['X_umap'][0:5, 0:2]
```

Out[215]:

```
array([[-5.55622101, -0.25772715,  0.18679433, -2.80009699,  0.05072495],
       [-7.20952702, -7.4820013 , -0.16271746,  8.01851654, -3.00661612],
       [-2.69443727,  1.58366168,  0.66312349, -2.20564294,  1.78901792],
       [10.1432972 ,  1.36853468, -1.20982373,  0.70006967,  2.90616465],
       [ 1.11281347,  8.15279865, -1.33235252,  4.252491  , -1.96318078]])
```

Out[215]:

```
array([[ 7.90665722,  3.55609107],
       [ 9.24834824, 12.54433155],
       [ 7.62998581,  3.83478546],
       [ 0.13157795,  5.53973913],
       [10.05534077, -0.64742917]])
```

In [216]:

```
# unstructured annotation
scezell_ann.uns.keys()
```

Out[216]:

```
dict_keys(['X_name', 'hvg', 'leiden', 'log1p', 'neighbors', 'pca', 'rank_genes_groups', 'umap'])
```

In [217]:

```
# pca variance
scezell_ann.uns['pca']['variance']
```

Out[217]:

```
array([32.11045456, 18.71865463, 15.60732937, 13.23528862,  4.80226898,
        3.98593235,  3.52623272,  3.23344541,  3.12120867,  3.07526112,
        2.99807477,  2.95952106,  2.95178485,  2.94424772,  2.913872  ,
        2.89903021,  2.88068199,  2.86468506,  2.84306359,  2.83575082,
        2.83142138,  2.81823635,  2.80355239,  2.79998732,  2.78895402,
        2.77810097,  2.77057672,  2.76022053,  2.75386024,  2.74595523,
        2.73718643,  2.73412681,  2.72220206,  2.71231079,  2.70247769,
        2.70004725,  2.68385029,  2.67905068,  2.67690778,  2.67399454,
        2.66485381,  2.65731144,  2.65117669,  2.64177561,  2.63297033,
        2.6295197 ,  2.62452936,  2.61837602,  2.61800337,  2.60186577])
```

In [218]:

```
# relationship of cells, graphs
scezell_ann.obsp
# RNA_nn graph
scezell_ann.obsp['distances'].toarray()
# RNA_snn graph
scezell_ann.obsp['connectivities'].toarray()
```

Out[218]:

```
PairwiseArrays with keys: connectivities, distances
```

Out[218]:

```
array([[0., 0., 0., ..., 0., 0., 0.],
       [0., 0., 0., ..., 0., 0., 0.],
       [0., 0., 0., ..., 0., 0., 0.],
       ...,
       [0., 0., 0., ..., 0., 0., 0.],
       [0., 0., 0., ..., 0., 0., 0.],
       [0., 0., 0., ..., 0., 0., 0.]])
```

Out[218]:

```
array([[0., 0., 0., ..., 0., 0., 0.],
       [0., 0., 0., ..., 0., 0., 0.],
       [0., 0., 0., ..., 0., 0., 0.],
       ...,
       [0., 0., 0., ..., 0., 0., 0.],
       [0., 0., 0., ..., 0., 0., 0.],
       [0., 0., 0., ..., 0., 0., 0.]])
```

In [219]:

```
# feature loadings
scezell_ann.varm['PCs']
```

Out[219]:

```
array([[-2.60148179e-02,  3.25416843e-03,  1.89788977e-03, ...,
        -5.18770702e-03,  1.44968908e-02, -6.67473301e-04],
       [-8.27822462e-03,  9.08316299e-03, -7.81411130e-04, ...,
         3.08727100e-02, -8.86981003e-03, -2.88053416e-03],
       [-3.31518659e-03,  3.20968428e-03,  2.79858650e-04, ...,
         1.01477914e-02, -5.30328136e-04,  1.50829612e-03],
       ...,
       [ 8.34176037e-03, -1.24651939e-03, -4.12195362e-03, ...,
        -1.01806019e-02,  9.22558550e-03,  2.79657058e-02],
       [-1.64065659e-02,  4.41013835e-02, -2.13347375e-05, ...,
         9.99553967e-03, -4.50964272e-03, -1.36533342e-02],
       [-1.51882619e-02,  4.00086790e-02,  5.41223399e-03, ...,
        -3.72782419e-03,  2.11074371e-02,  3.59644145e-02]])
```

#### scDIOR¶

**Retained information** (`SingleCellExperiemnt -> AnnData`):

- count matrix: scaled data (`scale.data -> X`), log-normalized data (`logcounts -> layers['logcounts']`), raw count matrix (`rawcounts -> layers['rawcounts']`)
- cells' meta-information (`colData -> obs`)
- annotation of features (`rowData -> var`)
- dimensional reduction results (`reducedDim -> obsm`)

scDIOR requires diopy to read `.h5` file.

In [220]:

```
%%R
# scDIOR does not support varm in rowData, diopy.input.read_h5 error: Error: "['varm'] not in index"
rowData(pbmc3k.sce)$varm = NULL
SCE2AD(sce.obj = pbmc3k.sce, method = "scDIOR", out.folder = "./")
```

```
[1] "The first 'assayNames' defaults to 'X'"
[1] "RNA"
```

In [221]:

```
# scDIOR require diopy to read h5
import diopy
```

In [222]:

```
scescdior_ann = diopy.input.read_h5(file = "./pbmc3k.sce_scDIOR.h5")
scescdior_ann
```

Out[222]:

```
AnnData object with n_obs × n_vars = 2638 × 1838
    obs: 'n_genes', 'n_genes_by_counts', 'total_counts', 'total_counts_mt', 'pct_counts_mt', 'leiden'
    var: 'gene_ids', 'n_cells', 'mt', 'n_cells_by_counts', 'mean_counts', 'pct_dropout_by_counts', 'total_counts', 'highly_variable', 'means', 'dispersions', 'dispersions_norm', 'mean', 'std'
    obsm: 'X_x_pca', 'X_x_umap'
    layers: 'logcounts', 'rawcounts'
```

In [223]:

```
# scaled count matrix
scescdior_ann.X
```

Out[223]:

```
array([[-0.17146961, -0.2808123 , -0.04667677, ..., -0.09826882,
        -0.20909512, -0.5312033 ],
       [-0.21458235, -0.37265328, -0.05480441, ..., -0.266844  ,
        -0.31314582, -0.5966543 ],
       [-0.3768877 , -0.29508454, -0.05752748, ..., -0.15865591,
        -0.17087644,  1.3789997 ],
       ...,
       [-0.20708963, -0.2504642 , -0.04639699, ..., -0.05114426,
        -0.16106427,  2.041497  ],
       [-0.1903285 , -0.2263338 , -0.04399936, ..., -0.00591774,
        -0.13521305, -0.48211104],
       [-0.33378935, -0.25358772, -0.05271561, ..., -0.07842438,
        -0.13032718, -0.47133783]], dtype=float32)
```

In [224]:

```
# log-normalized count matrix
scescdior_ann.layers['logcounts'].toarray()
```

Out[224]:

```
array([[0.       , 0.       , 0.       , ..., 0.       , 0.       ,
        0.       ],
       [0.       , 0.       , 0.       , ..., 0.       , 0.       ,
        0.       ],
       [0.       , 0.       , 0.       , ..., 0.       , 0.       ,
        1.429744 ],
       ...,
       [0.       , 0.       , 0.       , ..., 0.       , 0.       ,
        1.9370484],
       [0.       , 0.       , 0.       , ..., 0.       , 0.       ,
        0.       ],
       [0.       , 0.       , 0.       , ..., 0.       , 0.       ,
        0.       ]], dtype=float32)
```

In [225]:

```
# raw count matrix
scescdior_ann.layers['rawcounts'].toarray()
```

Out[225]:

```
array([[0., 0., 0., ..., 0., 0., 0.],
       [0., 0., 0., ..., 0., 0., 0.],
       [0., 0., 0., ..., 0., 0., 1.],
       ...,
       [0., 0., 0., ..., 0., 0., 1.],
       [0., 0., 0., ..., 0., 0., 0.],
       [0., 0., 0., ..., 0., 0., 0.]], dtype=float32)
```

In [226]:

```
# cells' meta-information
scescdior_ann.obs.head()
```

Out[226]:

|  | n\_genes | n\_genes\_by\_counts | total\_counts | total\_counts\_mt | pct\_counts\_mt | leiden |
| --- | --- | --- | --- | --- | --- | --- |
| index |  |  |  |  |  |  |
| AAACATACAACCAC-1 | 781.0 | 779 | 2419.0 | 73.0 | 3.017776 | 0 |
| AAACATTGAGCTAC-1 | 1352.0 | 1352 | 4903.0 | 186.0 | 3.793596 | 2 |
| AAACATTGATCAGC-1 | 1131.0 | 1129 | 3147.0 | 28.0 | 0.889736 | 0 |
| AAACCGTGCTTCCG-1 | 960.0 | 960 | 2639.0 | 46.0 | 1.743085 | 4 |
| AAACCGTGTATGCG-1 | 522.0 | 521 | 980.0 | 12.0 | 1.224490 | 5 |

In [227]:

```
# annotation of features
scescdior_ann.var.head()
# shape
scescdior_ann.var.shape
```

Out[227]:

|  | gene\_ids | n\_cells | mt | n\_cells\_by\_counts | mean\_counts | pct\_dropout\_by\_counts | total\_counts | highly\_variable | means | dispersions | dispersions\_norm | mean | std |
| --- | --- | --- | --- | --- | --- | --- | --- | --- | --- | --- | --- | --- | --- |
| index |  |  |  |  |  |  |  |  |  |  |  |  |  |
| TNFRSF4 | ENSG00000186827 | 155.0 | False | 155.0 | 0.077407 | 94.259259 | 209.0 | True | 0.277410 | 2.086050 | 0.665406 | -3.672069e-10 | 0.424481 |
| CPSF3L | ENSG00000127054 | 202.0 | False | 202.0 | 0.094815 | 92.518519 | 256.0 | True | 0.385194 | 4.506987 | 2.955005 | -2.372437e-10 | 0.460416 |
| ATAD3C | ENSG00000215915 | 9.0 | False | 9.0 | 0.009259 | 99.666667 | 25.0 | True | 0.038252 | 3.953486 | 4.352607 | 8.472988e-12 | 0.119465 |
| C1orf86 | ENSG00000162585 | 501.0 | False | 501.0 | 0.227778 | 81.444444 | 615.0 | True | 0.678283 | 2.713522 | 0.543183 | 3.389195e-10 | 0.685145 |
| RER1 | ENSG00000157916 | 608.0 | False | 608.0 | 0.298148 | 77.481481 | 805.0 | True | 0.814813 | 3.447533 | 1.582528 | 7.696297e-11 | 0.736050 |

Out[227]:

```
(1838, 13)
```

In [228]:

```
# dimensional reduction results of pca
scescdior_ann.obsm['X_x_pca'][0:5, 0:5]
# dimensional reduction results of umap
scescdior_ann.obsm['X_x_umap'][0:5, 0:2]
```

Out[228]:

```
array([[-5.55622101, -0.25772715,  0.18679433, -2.80009699,  0.05072495],
       [-7.20952702, -7.4820013 , -0.16271746,  8.01851654, -3.00661612],
       [-2.69443727,  1.58366168,  0.66312349, -2.20564294,  1.78901792],
       [10.1432972 ,  1.36853468, -1.20982373,  0.70006967,  2.90616465],
       [ 1.11281347,  8.15279865, -1.33235252,  4.252491  , -1.96318078]])
```

Out[228]:

```
array([[ 7.90665722,  3.55609107],
       [ 9.24834824, 12.54433155],
       [ 7.62998581,  3.83478546],
       [ 0.13157795,  5.53973913],
       [10.05534077, -0.64742917]])
```

In [ ]:
